# Genome-wide association study, network analysis, and reverse genetics pinpoint novel genes associated with seedling root growth variation of *Arabidopsis thaliana* under drought

**DOI:** 10.1101/2025.09.18.676994

**Authors:** Debankona Marik, Surbhi Vilas Tajane, Rishabh Kumar, Sucharita Dey, Ayan Sadhukhan

**Affiliations:** Department of Bioscience and Bioengineering, Indian Institute of Technology Jodhpur, Jodhpur, Rajasthan, India

**Keywords:** drought, GWAS, PEG, PLP salvage, root, stress granule

## Abstract

Development of drought-resilient crops requires a precise understanding of molecular signaling in the root, the primary organ encountering drought. This study unraveled novel genetic loci contributing to drought tolerance by exploiting the natural variation in seedling root growth of *Arabidopsis thaliana* under PEG-induced drought stress. Through a genome-wide association study (GWAS) of 207 worldwide *Arabidopsis thaliana* ecotypes from regions with varied rainfall, 68 protein-coding genes were identified, associated with the top 50 single-nucleotide polymorphisms (*P* < 10^− 3^), explaining 63% of the observed variation in root length. Subsequent network and functional enrichment analyses of the GWAS-delineated genes demarcated key biological processes crucial for maintaining root growth under drought, including DNA repair, tRNA editing, protein folding and quality control, cell cycle regulation, stress granule assembly, and the pyridoxal 5′-phosphate (PLP) salvage pathway regulating oxidative stress in roots. Expression level polymorphisms, promoter *cis*-element variations, and amino acid substitutions affecting predicted protein dynamics, with phenotype and climate associations, were identified. Finally, reverse genetic evaluation using T-DNA insertion knockout/knockdown mutants confirmed a direct association of the identified candidate genes, AT1G06690 (PLP pathway), AT4G26990 and *RBP45C* (stress granule assembly), *ACD55.5* (protein folding), *PCMP-A4* (RNA modification), *SKS6* and *ANAC094* (cell wall remodeling), and *INCENP* (cell cycle regulation), with seedling drought tolerance. Furthermore, the knockdown of AT1G06690 led to higher accumulation of hydrogen peroxide in root tissues, inhibiting growth. Future translation of the current findings into crops will provide new tools for the improvement of drought tolerance by modulating root traits through biotechnology and breeding.

## 1. Introduction

Drought, one of the most disastrous abiotic stresses, can be described as a state of exceptionally low precipitation for a prolonged period, resulting in hydrological imbalance in plants (Ahluwalia et al., 2021; Hooshyaripor et al., 2022). Drought decreases crop production due to physiological alterations in plants, such as stomatal closure, reduced photosynthesis, oxidative stress, and impaired cellular division and elongation, ultimately resulting in stunted growth (Hemati et al., 2022). Over the past few years, drought has led to a substantial loss of crops worldwide. In the year 2024 alone, the USA experienced a 37% reduction in wheat production due to an unprecedented drought (Science Advances, 2024), while in Italy, agricultural production dropped by 25% (European Commission Joint Research Centre, 2024), and India is no exception to it (https://agriwelfare.gov.in). Hence, it has become an absolute exigency to develop stress-resilient crops (Agho et al., 2024) through contemporary breeding methods that involve the introgression of quantitative trait loci (QTLs) regulating drought tolerance (Cooper & Messina, 2023), biotechnology (Şimşek et al., 2024), and emerging technologies such as gene editing (Pixley et al., 2023), epi-breeding (Schmid et al., 2018), and nanotechnology (Zhao et al., 2022). A precise understanding of the molecular signaling processes in different plant organs under drought is paramount to engineering tolerant plant varieties.

The root is the primary organ that encounters drought, and it senses the reduction in water potential of its environment through osmo- and mechano-sensors. This detection triggers an influx of ions, leading to the hyperpolarization of root cells, the signal being subsequently transduced by proteins including receptor-like kinases (RLKs), mitogen-activated protein kinases (MAPKs), and Ca^2+^-dependent protein kinases (CDPKs or CPKs) to the nucleus, resulting in the phosphorylation of transcription factors (TFs) that activate the expression of effector drought-tolerance genes in the root (Marik et al., 2025). A complex interaction among plant hormones: abscisic acid (ABA), auxin, cytokinin, and jasmonic acid (JA) regulates root development, meristem function, and cell division during dehydration stress. A decrease in water potential is transformed into hydraulic and electrical signals by plants, which, along with hormones like ABA, mRNAs, peptides, and osmolytes, move towards the shoot to produce a coordinated organismal response (Mittler et al., 2015). Moreover, plants adjust to drought conditions by altering their root plasticity and seeking water through tropic responses (Voothuluru et al., 2024).

Identification of drought-resistant genes utilizes forward genetic techniques such as QTL mapping and genome-wide association studies (GWAS). The natural allelic variation within a species is exploited in GWAS, rapidly identifying genome-wide candidate genes underlying complex quantitative traits like drought through statistical association (Kobayashi et al., 2016; Sadhukhan et al., 2017; Uffelmann et al., 2021; He and Gai, 2023). Extensive genetic variation exists in *Arabidopsis thaliana* natural ecotypes associated with drought adaptation (Exposito-Alonso et al., 2018). Previously, GWAS on shoot traits like leaf growth under mild drought stress in *A. thaliana* unraveled tolerance genes like *trehalase1* (*TRE1*), regulating the levels of the osmolyte trehalose (Clauw et al., 2016). Another GWAS on the variation in drought-induced ABA levels identified tonoplast transporters affecting ABA accumulation, signaling RLKs, and a START domain-containing lipid-binding protein regulating leaf water loss, having alleles associated with climate variation at the native geographical locations of the *A. thaliana* ecotypes (Kalladan et al., 2017). In recent years, genetic studies have started mapping important root traits underlying drought tolerance. Although rhizotron-based root phenotyping methods can be adapted to study drought in soil culture (Deja-Muylle et al., 2022), most studies relied on the simulation of drought conditions using polyethylene glycol (PEG) under hydroponic culture systems to observe root traits easily. PEG is a high-molecular-weight hydrophilic chemical, regularly used as an osmotic stressor to mimic drought in plant research (Khakwani et al., 2011). Because of its size, it cannot penetrate plant cells but induces water loss through cytorrhysis instead of plasmolysis. Hydroponic solutions infused with PEG result in decreased oxygen diffusion to the roots, and PEG buildup in tissues causes gradual dehydration (Rajeshwar et al., 2021). Hydroponics is a reproducible system where many genotypes can be cultivated in a compact area, allowing easy observation of root traits and making it a phenotyping platform of choice. A GWAS combined with co-expression network analysis conducted on maize inbred lines identified a TF, *ZmGRAS15*, of the gibberellin signaling pathway, whose overexpression improved drought tolerance by enhancing primary root growth (Wang et al., 2025). Another GWAS on rice root characteristics, viz., primary root length, root surface area, root diameter, etc., identified novel QTLs regulating drought tolerance (Sakhare et al., 2024). GWAS on root traits of sesame unveiled a single-nucleotide polymorphism (SNP) associated with root number and dry weight on the promoter of the *big root biomass* (*BRB*) gene, whose overexpression in Arabidopsis resulted in a decrease in root biomass and drought sensitivity (Dossa et al., 2021). Hence, GWAS on root traits can potentially identify yet unknown loci crucial for drought tolerance.

Motivated by these promising recent studies, we conducted a GWAS on the root growth variation of approximately 200 natural *A. thaliana* accessions distributed across diverse ecoregions of the globe with varying drought occurrences. The GWAS results were combined with transcriptome and gene network analyses, identifying key processes associated with natural variation in root growth of *A. thaliana*. Following analyses of gene expression and protein-level polymorphisms and reverse genetic validation of GWAS-identified genes by phenotyping T-DNA insertion mutants, previously studied and novel genes for drought tolerance involved in DNA repair, tRNA editing, protein quality control, cell cycle regulation, and stress granule assembly were unraveled. Our study revealed molecular signaling networks responsible for root growth maintenance under drought, using the natural variation in A. *thaliana*.

## 2. Materials and Methods

### 2.1. Plant materials and phenotyping for drought tolerance

Two hundred seven worldwide ecotypes of *A. thaliana* procured from the Nottingham Arabidopsis Stock Centre (NASC), UK, and the RIKEN Bioresource Research Centre (BRC), Japan, were maintained using the single-seed descent method, and fresh seeds were harvested before conducting GWAS. The seeds underwent imbibition and stratification at 4°C for three days in deionized water, ensuring synchronized germination, and were planted on nylon meshes mounted on photographic frames floated on ¼-Hoagland’s media, pH 5.8 (Hoagland & Arnon, 1938; Kobayashi et al., 2016). A total of 20 seedlings per genotype were cultivated hydroponically in ¼-Hoagland’s media alone or supplemented with 2.5% PEG-6000 (pH 5.8), imposing drought conditions. The growth room maintained 22-24°C and 55-60% humidity with a 12 h photoperiod and a photon flux density of 40 μmol m^−^² supplied using surface-mount device-type T8 LED lamps. The media was changed at two-day intervals. After 14 days, plants were carefully picked up with forceps and positioned on Petri plates with solidified agar. Equidistant imaging of the plates was performed, and the root length was determined by LIA 3.0 software (Sadhukhan et al., 2017). Finally, the percent relative root length (RRL; %) was determined by averaging the top eight longest root length values from both the control and drought conditions, then dividing the drought by the control and multiplying by 100. The broad-sense heritability (H ^2^) was determined by the ratio of the genetic variance to the sum of genetic and residual variance (Sadhukhan et al., 2017; Agrahari et al., 2021). The coordinates of the native geographical locations of the ecotypes were obtained from public databases (Atwell et al., 2010; Cao et al., 2011; Horton et al., 2012). The available yearly average precipitation data (in millimeters) between 1901 and 2022 of the exact locations were mined from a public database (https://climatecharts.net/) and correlated with the RRL values (Deja-Muylle et al., 2022). A map of geographical locations of *A. thaliana* ecotypes against a background of average annual precipitation was created using QGIS 3.4 Madeira (https://qgis.org/).

### 2.2. Genome-wide association study

GWAS calculations were conducted in TASSEL ver 5.0 (Bradbury et al., 2007) using the RRL phenotypes of a total of 207 *A. thaliana* ecotypes and genotype information containing a total of 105,856 single nucleotide polymorphisms (SNPs), after removing the missing data and minor alleles (< 10%) extracted from public databases, using a mixed linear model (MLM) taking into account the kinship and population structure (Kobayashi et al., 2016; Sadhukhan et al., 2017). A quantile-quantile (Q-Q) plot of observed versus expected *P*-value distribution was used to visually inspect the GWAS results using TASSEL (Agrahari et al., 2021). A ridge regression analysis was conducted to observe the cumulative effect of top-ranking SNPs identified by the GWAS using the glmnet R package after five-fold cross-validation repeated 100 times (Friedman et al., 2010). The mean correlation coefficient (*r^2^*) between the observed and expected RRL of 100 repeats was plotted against the number of top-ranking SNPs delineated by the GWAS. Haploblocks were estimated around the most significant SNPs (*P* < 10^−3^) by assessing the linkage disequilibrium (LD) of the neighboring SNPs within the LD decay range of 10 Kb (Buckler and Gore, 2007) using TASSEL (*r^2^* ≥ 0.6), and genes within such haploblocks were listed using gene coordinates from TAIR (https://www.arabidopsis.org/) by in-house Microsoft Excel macro programs. The GWAS-identified genes were subjected to an overrepresentation test in Panther (https://pantherdb.org/) and ShineyGO 0.85 (https://bioinformatics.sdstate.edu/go/) using Fisher’s exact test corrected with Benjamini-Hochberg *false discovery rate* (*FDR*) to identify the enriched Gene Ontology (GO) biological processes.

### 2.3. Network analysis

Network analysis of the GWAS-delineated genes was carried out using the STRING database (https://string-db.org/) (Szklarczyk et al., 2023). Up to 50 interactors in the first shell of the GWAS-identified genes were selected, and a high confidence score of 0.7 was used to filter meaningful interactions. Both gene co-expression and experimentally determined protein-protein interactions were considered for network construction. The results were visualized using Cytoscape version. 3.10.3 (Shannon et al., 2003). The hub genes were identified by Cytohubba using the maximal clique centrality (MCC) algorithm (Ono et al., 2025). This was followed by cluster analysis using MCODE version 2.0.3 to shortlist hub genes with node degree ≥ 2. The OMICS visualizer plugin was used to visualize functional enrichment analysis of the networking genes and represented as colored rings around the nodes (Legeay et al., 2020). The Plant Cistrome database hosting DNA-affinity purification sequencing (DAP-seq) data (http://neomorph.salk.edu/dap_web/pages/index.php) was used to fetch the target genes of GWAS-identified TFs.

### 2.4. Transcriptome analysis

For the transcriptome analysis, Col-0 seedlings were grown in ¼-Hoagland’s media for 12 days, and thereafter transferred to media with 5% PEG for 3 days, followed by snap freezing in liquid nitrogen. The CTAB method-based RNA isolation followed by cDNA library preparation using the NEBNext^®^ Ultra^TM^ II RNA library kit (New England Biolabs, New Delhi, India) was carried out before sequencing in the Illumina Nova-Seq 6000 V1.5 (Illumina Inc., Gurgaon, India). The raw data were submitted to NCBI SRA with accession number PRJNA1086208. Raw data quality control, adapter removal, and alignment with the reference *A. thaliana* genome were conducted as described previously (Dey et al., 2025). Differential gene expression was estimated using the edgeR version 4.1.2 (Chen et al., 2025) and subjected to Fisher’s exact test corrected with Benjamini-Hochberg *FDR*. The differential gene expression (*FDR* < 0.05, *N* = 3 biological replicates) data were mapped to the GWAS results.

### 2.5. *In silico* expression analysis

RNAseq datasets for various seedling tissues of *A. thaliana* were extracted from the Expression Atlas (https://www.ebi.ac.uk/gxa/home) (Mergner et al., 2020). Each gene’s transcripts per million (TPM) values, averaged separately for seedling root and shoot samples, were further normalized using a logarithm base two transformation to stabilize the variance and prepare the data for visualization. The log_2_(TPM+1) values were used to generate a hierarchical clustering using the TBtools software, using the Euclidean distance metric and a complete linkage method (Chen et al., 2020). In addition, we mined the NCBI Gene Expression Omnibus for the expression levels of the GWAS-identified genes under osmotic stress, high salt, and ABA in other studies. Expression data of the wild-type (WT) *A. thaliana* (Col-0) seedlings treated with 100 mM NaCl (GSE176433; Wang et al., 2021), 100 μM ABA (GSE127910), and 300 mM mannitol (GSE157435; Chen et al., 2020) versus non-stressful conditions were extracted and used for comparison with our in-house transcriptome data.

### 2.6. Sequence polymorphism analysis and local association study

Denser polymorphism information in the GWAS-identified genes was available for 142 out of the 207 ecotypes in the 1001 Genomes database (https://tools.1001genomes.org/pseudogenomes/) (Alonso-Blanco et al., 2016). The POLYMORPH 1001 genome variants web tool (http://tools.1001genomes.org/polymorph/) was used to extract polymorphism information. For the analysis of promoter and 5’-UTR polymorphisms, nucleotide sequences up to 1 kb upstream from the start codon, and for amino acid polymorphisms, the cDNA sequences were retrieved and analyzed using GENETYX ver. 11 software (GENETYX, Tokyo, Japan). Introns were removed, the nucleotides were converted to amino acids, and all polymorphism positions were verified carefully. A local association study (LAS) was conducted in TASSEL using denser polymorphisms in the GWAS-identified genes in 142 ecotypes using both the general linear model (GLM) and MLM, separately for different genomic regions (Sadhukhan et al., 2021).

### 2.7. Expression level polymorphism analysis

To analyze the expression level polymorphism (ELP) of the genes with promoter polymorphisms identified in the LAS, 40 randomly selected ecotypes used in the LAS were grown for 10 days in control ¼-Hoagland’s media to grow sufficient root biomass and then transferred to media with 2.5% PEG for five days before harvesting the roots in liquid nitrogen and RNA isolation. The RevertAid reverse transcriptase kit (K1691; Thermo-scientific, Mumbai, India) and random hexamer primers were used for cDNA preparation. Real-time quantitative PCR (qPCR) was carried out using TB Green master mix (RR420; Takara, India) in a QuantStudio™ 5 Real-Time PCR System (Thermo). The Primer3 webtool (Kõressaar et al., 2018) was used to design primers, and the relative expression levels were analyzed by the standard curve method, using *AtUBQ1* as an internal control. The expression levels were further normalized with respect to the levels in Col-0 using the 2^−ΔΔCt^ method (Livak & Schmittgen, 2001).

### 2.8. Analysis of promoter *cis*-elements

The sequences up to 1 kb upstream of the start codon of each gene were mined from TAIR and separated into stretches of eight nucleotides or octamers. The frequency of occurrence of each octamer was calculated in a list of up-(log_2_FC > 1) and downregulated (log_2_FC < -1) genes under drought (in-house transcriptome; PRJNA1086208), osmotic (GSE157435), and high salt (GSE176433) stresses, and ABA treatment (GSE127910) to WT seedlings. The relative appearance ratio (RAR) of each octamer in the promoter of genes up- or downregulated under a particular condition versus *A. thaliana* genome-wide genes was calculated as described earlier (Yamamoto et al., 2011; Sadhukhan et al., 2014) using in-house Python scripts and plotted against promoter positions. Octamers with RAR > 2 and Fisher’s test *P*-value < 0.05 were considered overrepresented. Promoter SNPs were mapped to the RAR plot. Known promoter *cis* elements were predicted using the PLANTCARE webtool (https://bioinformatics.psb.ugent.be/webtools/plantcare/html/).

### 2.9. Protein structure prediction and molecular dynamics simulations

Homology searches against the amino acid sequences of Col-0 were performed using NCBI BLASTP (Altschul et al., 1990) against the Protein Data Bank (PDB) (https://www.rcsb.org/; Berman, 2000) and UniProtKB/Swiss-Prot (https://www.uniprot.org/) databases. The search was filtered to include homologous proteins with 40–100% sequence identity and 80–100% query coverage to ensure relevant and high-confidence homolog retrieval for further structural and functional analyses. In the absence of homologous proteins in the PDB, the protein structures were modeled using the AlphaFold 3 web server (Jumper et al., 2021). GO analysis was performed using InterPro (Blum et al., 2025) to identify the molecular functions, biological processes, and cellular components associated with each protein. To determine the presence of functional domains and ligand/cofactor binding sites, relevant information was retrieved from the UniProt database (UniProt Consortium, 2025). All-atom molecular dynamics (MD) simulations of the apo proteins were performed using the GROMACS 2024 package with the Optimized Potentials for Liquid Simulations all-atom force field (Ponder & Case, 2003). To ensure better sampling, production runs of 200 ns each were carried out for four protein systems in explicit solvent. For each titratable residue, the default protonation state was used. After positioning the complex in the middle of a cubic box, TIP3P water molecules were used to solvate it. Na^+^ and Cl^−^ ions were added to the system to neutralize its net charge, and the steepest descent algorithm was used to minimize energy and eliminate any unfavorable interactions. The system was gradually heated to 300 K over 100 ps under constant number of particles, volume, and temperature (NVT) conditions, followed by 100 ps equilibration under constant number of particles, pressure, and temperature (NPT). Subsequently, a 200-ns final MD production run was performed with each step of 2 fs time. The simulations were repeated twice for each protein, and the trajectory with better stability metrics was selected for analysis. The Particle Mesh Ewald (PME) approach was used to identify electrostatic interactions, with periodic boundary constraints in all directions. Trajectory analyses were performed using GROMACS analysis tools and in-house Python scripts. HBPLUS was used to estimate hydrogen bonds (McDonald & Thornton, 1994). Ionic interactions were defined as contacts between side-chain atoms of positively (Lys, Arg, His) and negatively (Asp, Glu) charged residues within 4.0 Å (Gowri Shankar et al., 2007), and hydrophobic interactions as contacts between side-chain atoms of hydrophobic residues within 4.0 Å.

### 2.10. Confirmation of homozygosity and phenotyping T-DNA insertion mutants

T-DNA insertion mutants of the GWAS-identified genes were mined from the Salk T-DNA Express database (http://signal.salk.edu/cgi-bin/tdnaexpress) and procured from NASC. Plants were grown from individual seeds hydroponically for two weeks, followed by culture in a mixture of garden soil, soilrite, and vermiculite 1:2:1 in two-inch pots with Arasystem (https://www.arasystem.com/). Leaves from each plant were harvested by snap-freezing in liquid nitrogen, crushed by bead beating, and then DNA was isolated by the CTAB method. The DNA from both WT (Col-0; NASC stock no. N60000) and T-DNA insertion lines was used for two PCRs, one using a left genomic primer (LP) and a right genomic primer (RP), and the other with a T-DNA left border (LB) and RP, for checking homozygosity. SALK-recommended LB primers, SALK LBb1.3, WISC, and SAIL LB1 were used (http://signal.salk.edu/tdnaprimers.2.html). Individual homozygous lines were multiplied by the single-seed descent method for phenotyping. The degree of gene knockdown in T-DNA insertions in the promoter and 5’-untranslated region (5’-UTR) was analyzed by comparing the gene expression level between the T-DNA insertion lines and the WT, after a 5-day drought treatment of 10-day-old seedlings, by qPCR, as described in section 2.7. Independent homozygous lines of each T-DNA insertion, along with the WT, were grown in hydroponics and phenotyped under the same conditions as the GWAS phenotyping described in section 2.1. Additional phenotyping was carried out in ½-strength Murashige Skoog (MS) media, with 1.5% sucrose, pH 5.8, supplemented with 100 and 150 mM NaCl, and 2.5 and 5 µM ABA, solidified with 0.7% agar. Seedlings were grown for two weeks from germination before photography. The increase or decrease in RRL was calculated as (RRL of mutant - RRL of WT)/RRL of WT, and expressed as a percentage. A one-way analysis of variance (ANOVA) followed by a post hoc Tukey’s honestly significant difference (HSD) test was used to determine significant differences between multiple samples at *P* < 0.05.

### 2.11. Quantification of reactive oxygen species in roots

The levels of the reactive oxygen species (ROS), viz., hydrogen peroxide (H_2_O_2_), were quantified spectrophotometrically as described earlier by Sagisaka (1976). Twenty-four randomly selected ecotypes used in the ELP analysis, along with independent Col-0 and T-DNA insertion lines, were grown for 10 days in control ¼-Hoagland’s media and then transferred to media with 2.5% PEG for five days before harvesting the roots and snap freezing in liquid nitrogen. The roots were crushed by bead beating with the immediate addition of 1 mL 5% perchloric acid. Samples were mixed by inverting the tubes and centrifuged at 14,000 rpm for 5 min. To the supernatant, 200 µl of 2.5 M potassium thiocyanate, 400 µl of 50 % trichloroacetic acid, and 400 µl of 10 mM ferrous ammonium sulphate were added. Samples were centrifuged again, and the absorbance was measured at 480 nm. The standard curve was prepared by gradually increasing the concentration of pure H_2_O_2_ from 0 mM to 120 mM and measuring the absorbance at 480 nm. Finally, the H_2_O_2_ levels were expressed as µmol g^−1^ tissue fresh weight (FW).

## 3. Results

### 3.1. Natural variation of *Arabidopsis thaliana* in seedling root growth under drought

A diversity panel comprising 207 worldwide *A. thaliana* accessions from arid, semi-arid, and wet ecoregions (Fig. 1a) showed a wide variation in seedling root growth under PEG-induced drought stress in the laboratory. The RRL varied from 3.4% to 150.9% with a median of 38.6%. The H_b_^2^ was estimated at 0.97 (Fig. 1b). Among the ecotypes, Rsch-0, Pla-3, Tu-0, Ts-1, Yeg-1, Bsch-0, Ra-0, Tu-SB30-3, Buckhorn Pass, and Star-8 demonstrated high sensitivity, while Da(1)-12, El-0, Lp2-2, Mnz-0, Ha-0, Dr-0, Kin-0, Col-0, Hl-3, and Eil-0 exhibited high drought tolerance (Table S1). A slight negative correlation in drought tolerance with the precipitation received by the native geographical locations of the ecotypes (Zepner et al., 2020) was observed (Fig. 1c). The ecotypes from humid areas (precipitation > 1000 mm/year) showed sensitivity to drought, as observed for Buckhorn Pass (10.3%), Vezzano-2 (32.9%), Zu-1 (25.8%), Alst-1 (16.3%), Nz-1 (21.9%), Pog-0 (19.3%), Boot-1 (38.0%), Ba-1 (31.7%), and Ty-0 (11.9%). Most of the highly tolerant (RRL > 60%) ecotypes came from moderate precipitation areas of Europe (600-800 mm/year). Ecotypes from the semi-arid regions (500-600 mm/year) showed high tolerance, e.g., Eil-0 (150.9%), Lp2-2 (99%), Jl-3 (94.2%), Mer-6 (81.6%), and Borsk-2 (64.2%). While a few ecotypes from hyper-arid regions of Asia and Africa (<500 mm/year) showed tolerance, e.g., Kondara (92%), others showed sensitivity, e.g., Aitba-2 (31.3%), Cdm-0 (27.6%), Se-0 (26%), Kz-9 (24.3%), Toufl-1 (17.3%), Stepn-2 (15.3%), and Yeg-1 (9.7%).

**Fig. 1.**
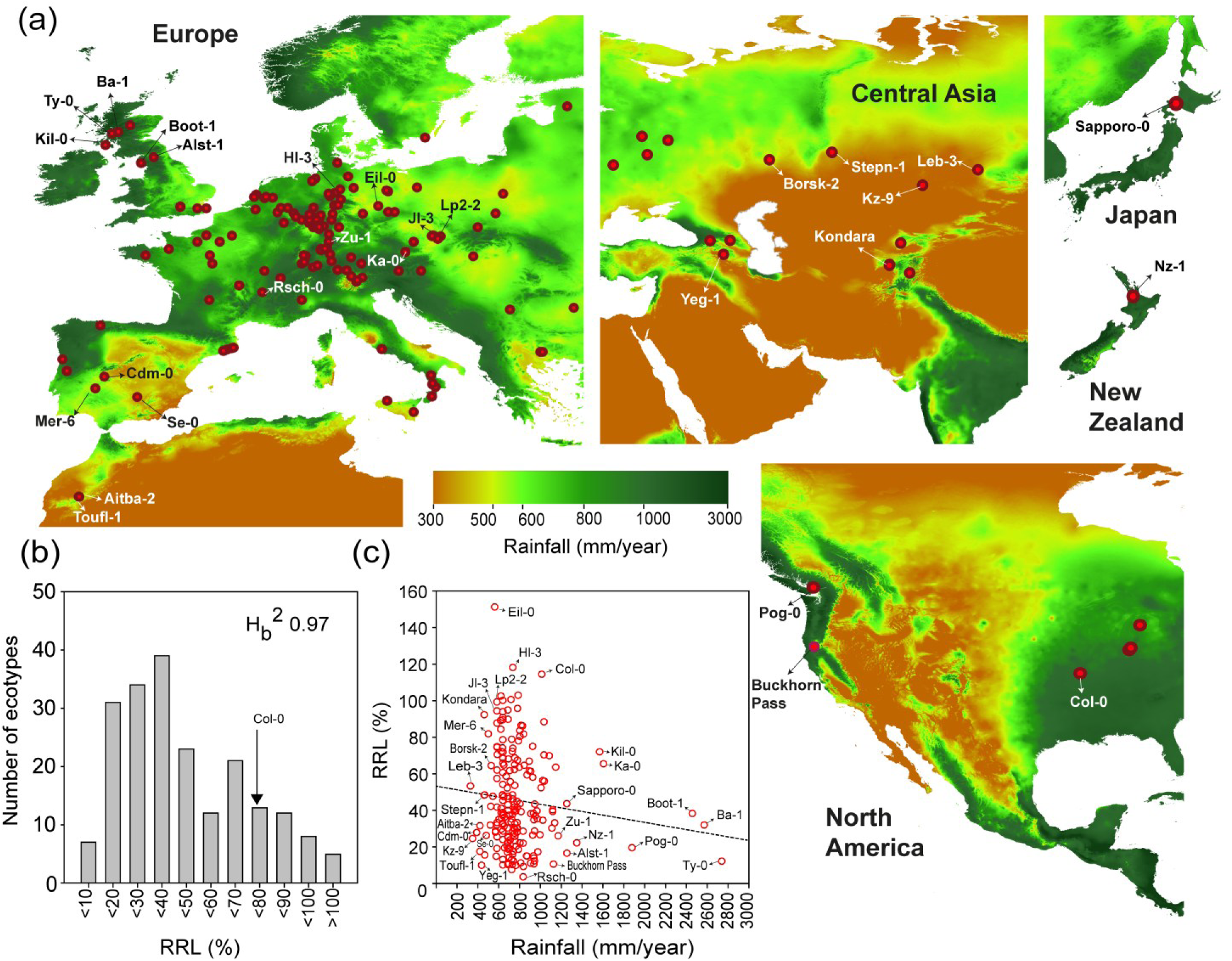
Natural variation of seedling drought tolerance in *Arabidopsis thaliana*. Two hundred seven *A. thaliana* ecotypes were grown hydroponically from seed in modified ¼-Hoagland’s solution for two weeks without (control) or with 2.5% PEG-6000, placed on solid media, and photographed. The relative root length (RRL; %) in PEG/control media was used as a phenotype. (**a**) The native geographical distribution of the 207 *A. thaliana* ecotypes used for the GWAS is shown (https://www.google.com/maps). The maps show parts of Eurasia, North America, New Zealand, and Japan (not to scale). The color scale indicates yearly average rainfall in millimetres between 1901 and 2022 (https://climatecharts.net/). Red dots show the locations of the ecotypes. The average data of 10 seedlings for each ecotype were used. (**b**) A histogram of the frequency distribution of RRL phenotypes in 207 *A. thaliana* ecotypes is shown. An arrow shows the reference genotype, Columbia (Col-0). The broad sense heritability (H_b_^2^), a ratio of the genetic variance to the sum of genetic and residual variance, is shown. (**c**) A correlation plot of RRL versus average annual rainfall of the exact native geographical locations of the ecotypes is shown, with a linear regression denoted by a dotted line. Ecotypes with extreme phenotypes and possible climate associations are indicated in the map and correlation plot.

### 3.2. A genome-wide association study unravels loci for seedling drought tolerance in *Arabidopsis thaliana*

A GWAS of root length variation in 207 global *A. thaliana* ecotypes identified loci associated with seedling drought tolerance. The QQ plot showed a deviation from the expected *P*-values at the tail region at *P*-value < 10^−3^, suggesting significance of association (Fig. 2a). A ridge regression analysis of the GWAS results showed that ∼63% of the observed variation in RRL could be explained by the top 50 SNPs having an MLM *P*-value < 10^−3^ (Fig. 2b). Within haploblocks at a distance of 10 kb from these SNPs, 68 protein-coding genes were localized (Table 1). The topmost ranking SNPs, *Chr1: 2051839* and *Chr 1:2051659*, having *P-values* of 2.26 × 10^−6^ and 5.89 × 10^−6^, respectively, were located on the exons of *NAD(P)-linked oxidoreductase superfamily protein* (AT1G06690) (Fig. 2c, Table 1), a chloroplast stroma, thylakoid membrane, and plastoglobule-localized aldo-keto reductase involved in the pyridoxal 5′-phosphate (PLP)/ vitamin B_6_ salvage pathway and tolerance to osmotic stress through antioxidant regulation (Herrero et al., 2011; Lundquist et al., 2012; Yu et al., 2020). Analysis of gene expression in various seedling tissues of *A. thaliana* in public databases indicated that the majority of GWAS-detected genes were expressed in both shoots and roots. The gene expression data indicated that the topmost ranking gene, AT1G06690, is also localized in roots and possibly is a component of the stroma or plastoglobules in root plastids (Ytterberg et al., 2006; Eugeni Piller et al., 2012; Lundquist et al., 2012). However, *trehalose-6-phosphate phosphatase I* (*TPPI*; AT5G10100), *response to low sulfur 4* (*LSU4*; AT5G24655), *Embryo defective 2788* (*EMB2788*; AT4G27010), and *NAC domain-containing protein 94* (*ANAC094*; AT5G39820) showed more than 2-fold higher expression in roots. On the other hand, AT2G22942, inner centromere protein (*INCENP*; AT5G55820), and *weak chloroplast movement under blue light protein* (*DUF827*; AT4G17210) were shoot-specific (Fig. 2d). These results suggested a synergistic role of both shoot- and root-expressed genes in the regulation of seedling root growth under drought in *A. thaliana*.

**Fig. 2.**
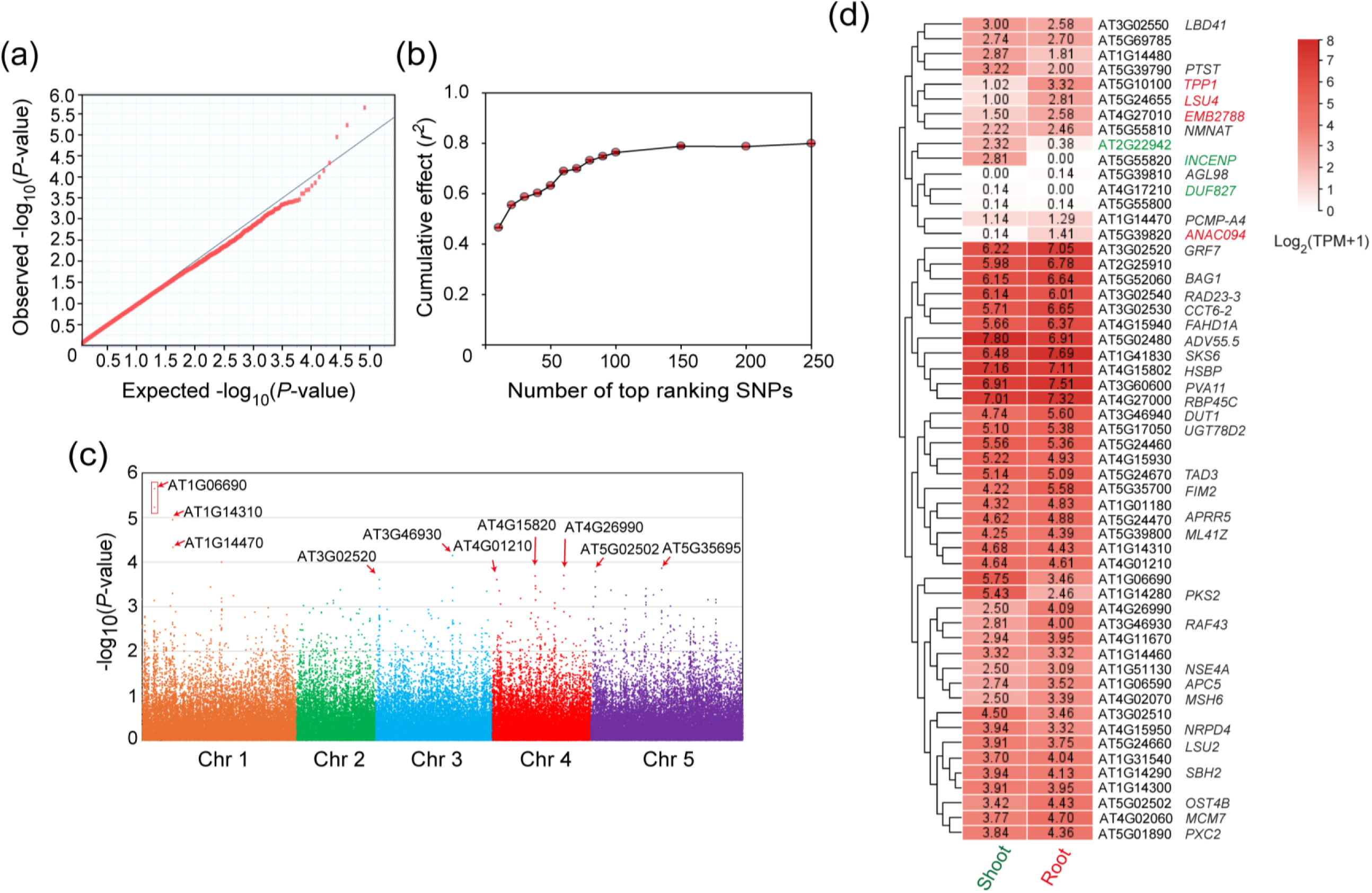
Genome-wide association study of drought tolerance in *Arabidopsis thaliana*. The results of a genome-wide association study (GWAS) involving the drought tolerance phenotypes of 207 worldwide *A. thaliana* ecotypes, single-nucleotide polymorphisms (SNPs), and population structure. (**a**) A quantile-quantile (Q-Q) plot shows the observed versus expected *P*-value distribution of SNPs in the GWAS. The straight line indicates the null distribution of *P*-values. (**b**) Results of a ridge regression analysis are shown. The mean correlation coefficient (*r^2^*) between predicted and observed RRLs with standard deviation for 100 repeats plotted against the number of top-ranking SNPs identified by the GWAS. (**c**) GWAS results are shown as a Manhattan plot of the significance of association (-log_10_*P*-value) vs. physical positions on the five *A. thaliana* chromosomes. Arrows indicate the locations of the protein-coding genes nearest to the suggestive SNPs. (**d**) A heatmap showing a hierarchical clustering of genes within haploblocks of the most significant SNPs (*P* < 10^−3^) detected by the GWAS based on their expression levels in seedling shoot and root tissues. Darker shades of red indicate higher expression levels in transcripts per million (TPM) normalized using a logarithm base 2 transformation to stabilize the variance (see Methods).

**Table 1.**
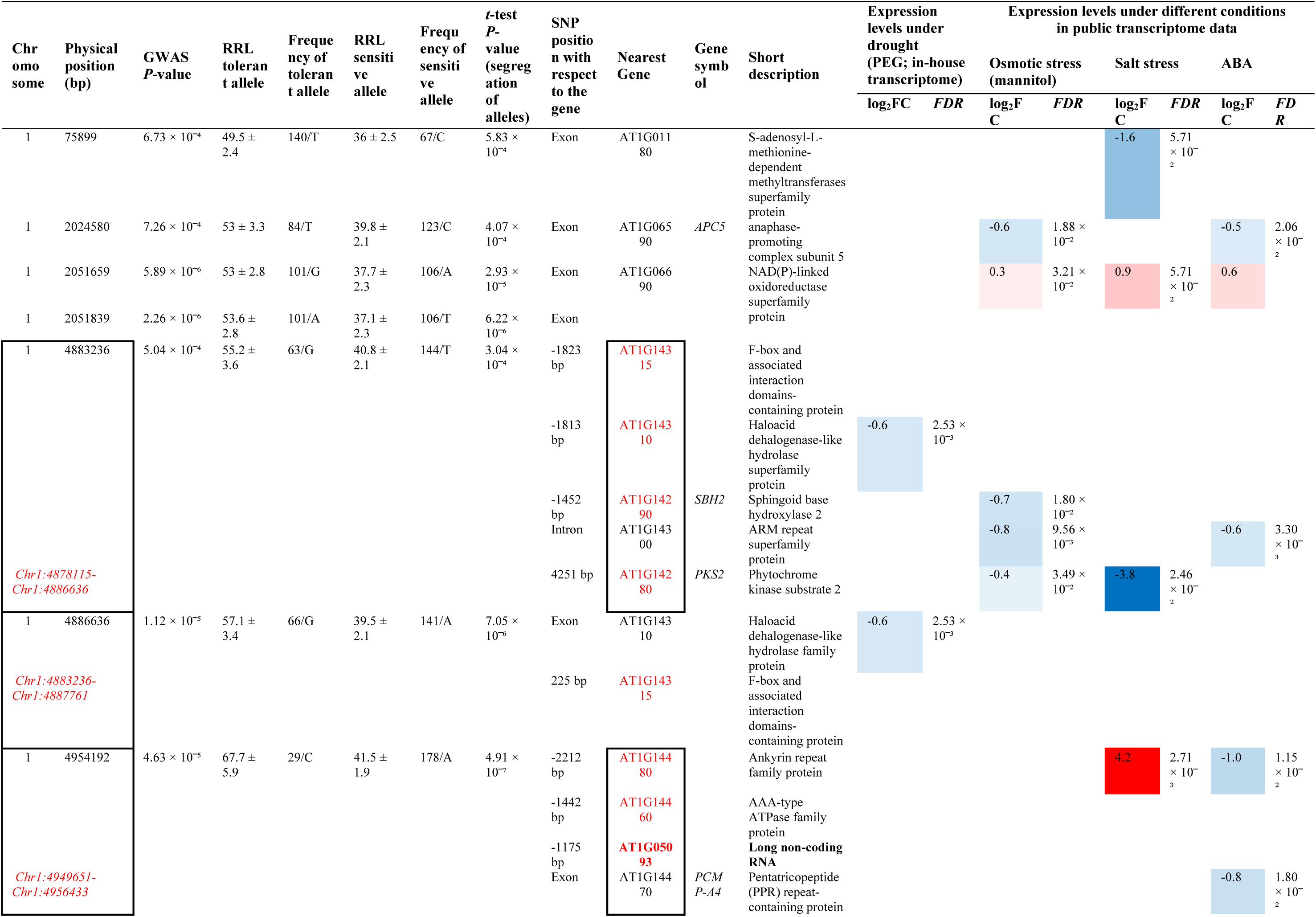
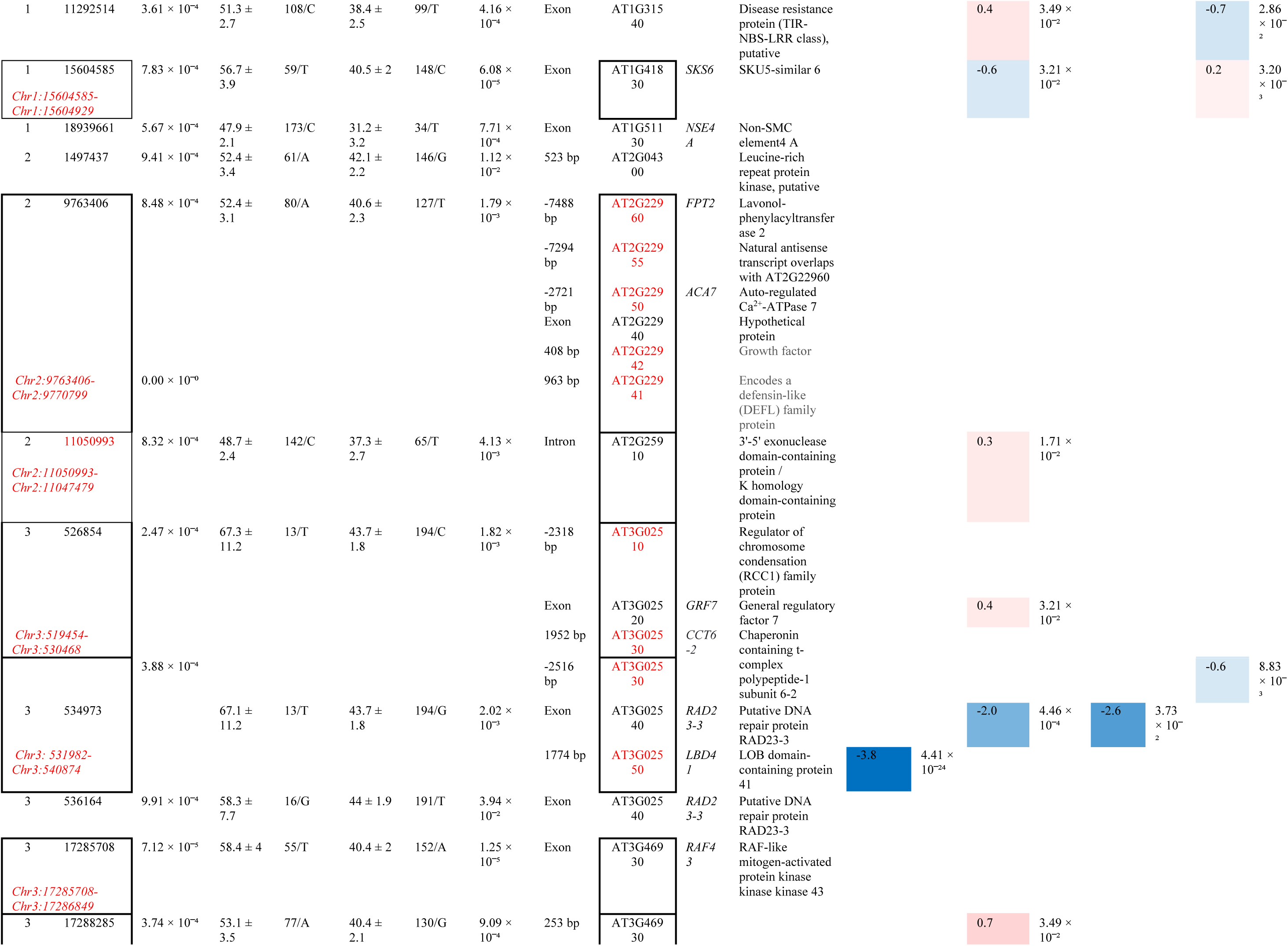
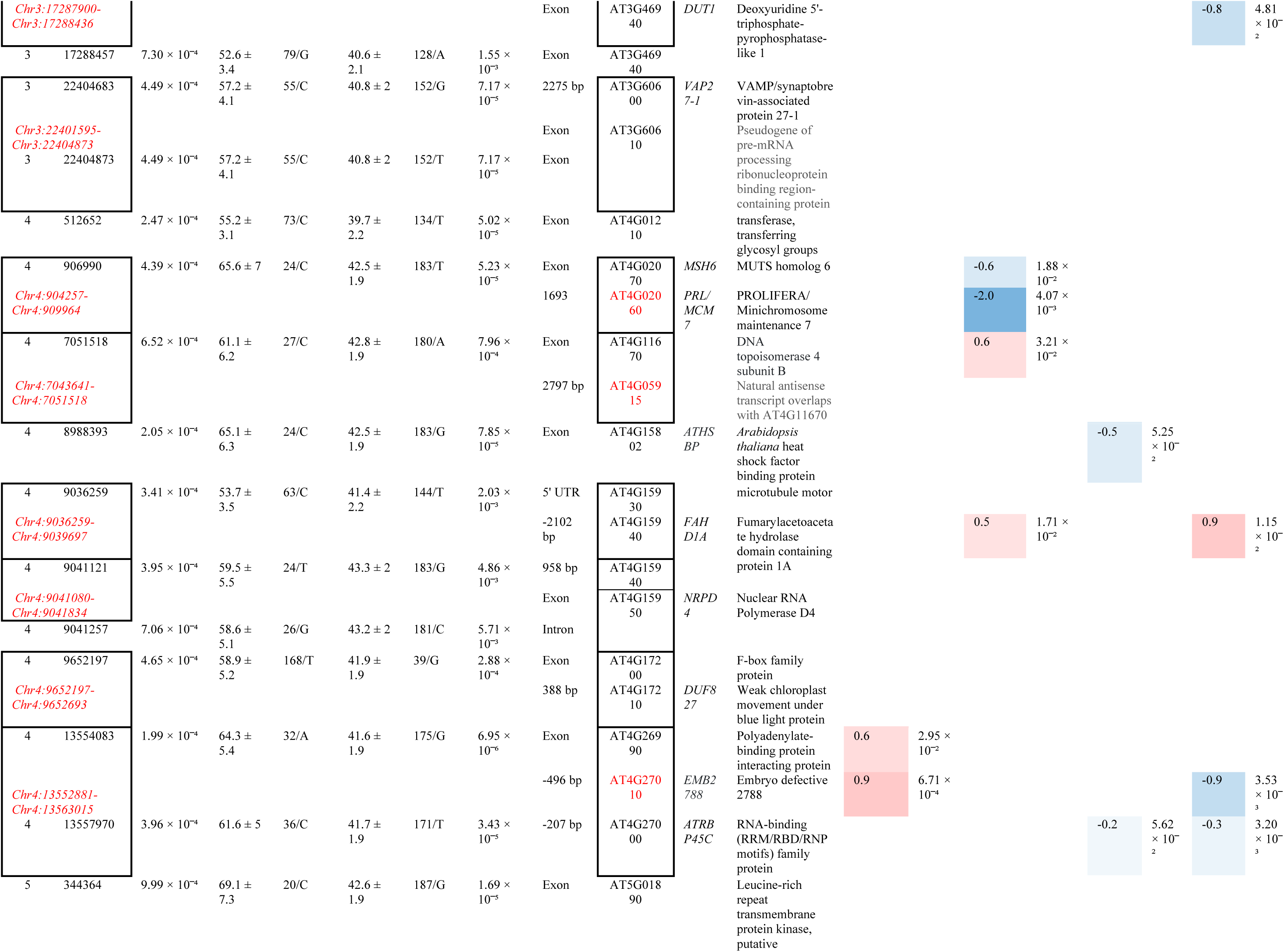
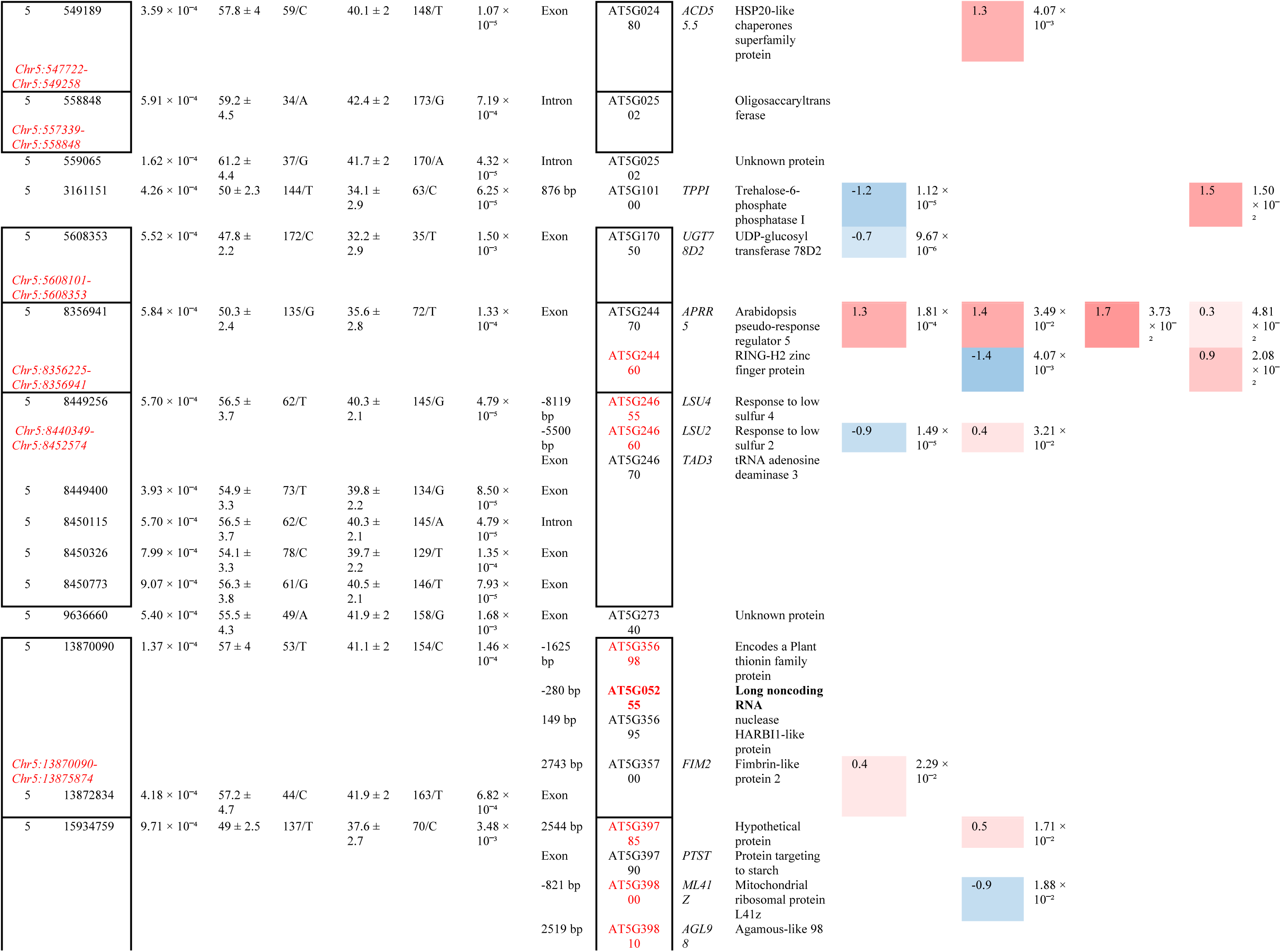
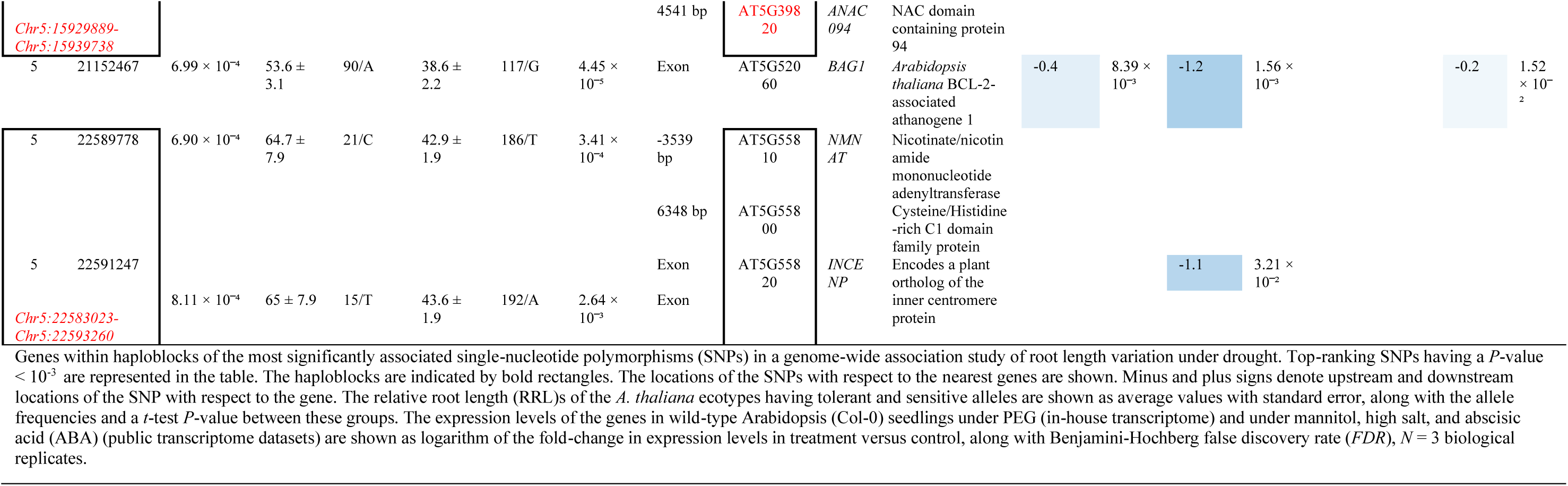
Genes associated with the most significant SNPs in a genome-wide association study of drought tolerance in *Arabidopsis thaliana*.

### 3.3. Stress-responsive genes involved in various biological processes underlie seedling drought tolerance variation in *Arabidopsis thaliana*

A GO enrichment analysis of the 68 protein-coding genes associated with the most significant SNPs (*P* < 10^−3^) showed that ten genes belonged to the enriched category of ‘cellular response to stress,’ indicating that the GWAS has identified genes of various stress-adaptive biological processes. Out of these, the majority of six genes functioned in ‘DNA repair,’ possibly protecting the DNA from drought stress-induced damage (Table 2). These included an *AAA-type ATPase family protein* (AT1G14460), *non-SMC element 4A* (*NSE4A*; AT1G51130), *radiation-sensitive 23-3* (*RAD23-3*; AT3G02540), *deoxyuridine 5’-triphosphate-pyrophosphatase-like 1* (*DUT1*; AT3G46940), *prolifera/minichromosome maintenance 7* (*PRL*/*MCM7*; AT4G02060), and *MUTS homolog 6* (*MSH6*; AT4G02070). *DUT1* is also involved in the ‘pyrimidine deoxyribonucleoside monophosphate biosynthetic process,’ possibly supplying nucleotides for damage repair under stress. *Arabidopsis thaliana heat shock factor binding protein* (*ATHSBP*; AT4G15802) is a corepressor of heat shock TFs involved in ‘cellular heat acclimation.’ A *polyadenylate-binding protein interacting protein* (AT4G26990) was involved in ‘stress granule assembly’ (Marondedze et al., 2020). Another gene within the same haploblock, *RNA-binding (RRM/RBD/RNP motifs) protein 45C* (*RBP45C*; AT4G27000), is also localized in stress granules (Yan et al., 2022). *EMB2788*, a ribosome biogenesis protein, is involved in the enriched process, ‘maturation of 5.8S rRNA from tricistronic rRNA transcript (small subunit-rRNA, 5.8S rRNA, large subunit-rRNA),’ indicating the role of ribosome heterogeneity in drought tolerance (Dias-Fields & Adamala, 2022). Two genes, *response to low sulfur 2* (*LSU2*; AT5G24660) and *LSU4* (AT5G24655), were involved in ‘cellular response to sulfur starvation’ as well as ‘cellular oxidant detoxification,’ indicating the importance of ROS scavenging in drought tolerance. *RAD23-3*, *anaphase-promoting complex subunit 5* (*APC5*; AT1G06590), and *BCL-2-associated athanogene 1* (*BAG1*; AT5G52060) were involved in ‘proteasome-mediated ubiquitin-dependent protein catabolic process’ and ‘cytoplasm protein quality control,’ possibly clearing aggregated or misfolded proteins under stress, highlighting the importance of these processes in drought stress tolerance. *ATHSBP* and *APC5* are also involved in ‘positive regulation of mitotic cell cycle,’ possibly contributing to root cell growth under drought conditions. Other enriched processes may have significant roles in drought stress adaptation. These included *TPPI* and *UDP-glucosyl transferase 78D2* (*UGT78D2*; AT5G17050), *sphingoid base hydroxylase 2* (*SBH2*; AT1G14290) involved in ‘trehalose,’ ‘coumarin,’ and ‘sphingoid’ biosynthesis processes, respectively, indicating the involvement of osmolyte sugars, secondary metabolites, membrane stabilization, and ROS and sphingolipid signaling in maintaining root growth under drought. Association of *VAMP/synaptobrevin-associated protein 27-1* (*VAP27-1*; AT3G60600), involved in ‘endoplasmic reticulum–plasma membrane (ER–PM) tethering,’ indicated the crucial role of protein trafficking under drought stress (Sampaio et al., 2022). A gene of the enriched functional category ‘tRNA wobble adenosine to inosine editing,’ *tRNA adenosine deaminase 3* (*TAD3*; AT5G24670), suggests the role of RNA editing for maintaining growth under drought (Zhou et al., 2013). *DUF827* of the WEB1/PMI2-related (WPR) protein family (Kodama et al., 2011), involved in ‘chloroplast accumulation movement,’ regulating photorelocation response, possibly optimizing photosynthetic activity or avoiding photodamage. Finally, two genes involved in ‘photomorphogenesis,’ *Arabidopsis pseudo-response regulator 5* (*APRR5*; AT5G24470) and *SBH2*, highlight the cross-talk of light and drought signaling for seedling growth under drought stress (Mukherjee et al., 2023).

**Table 2.**
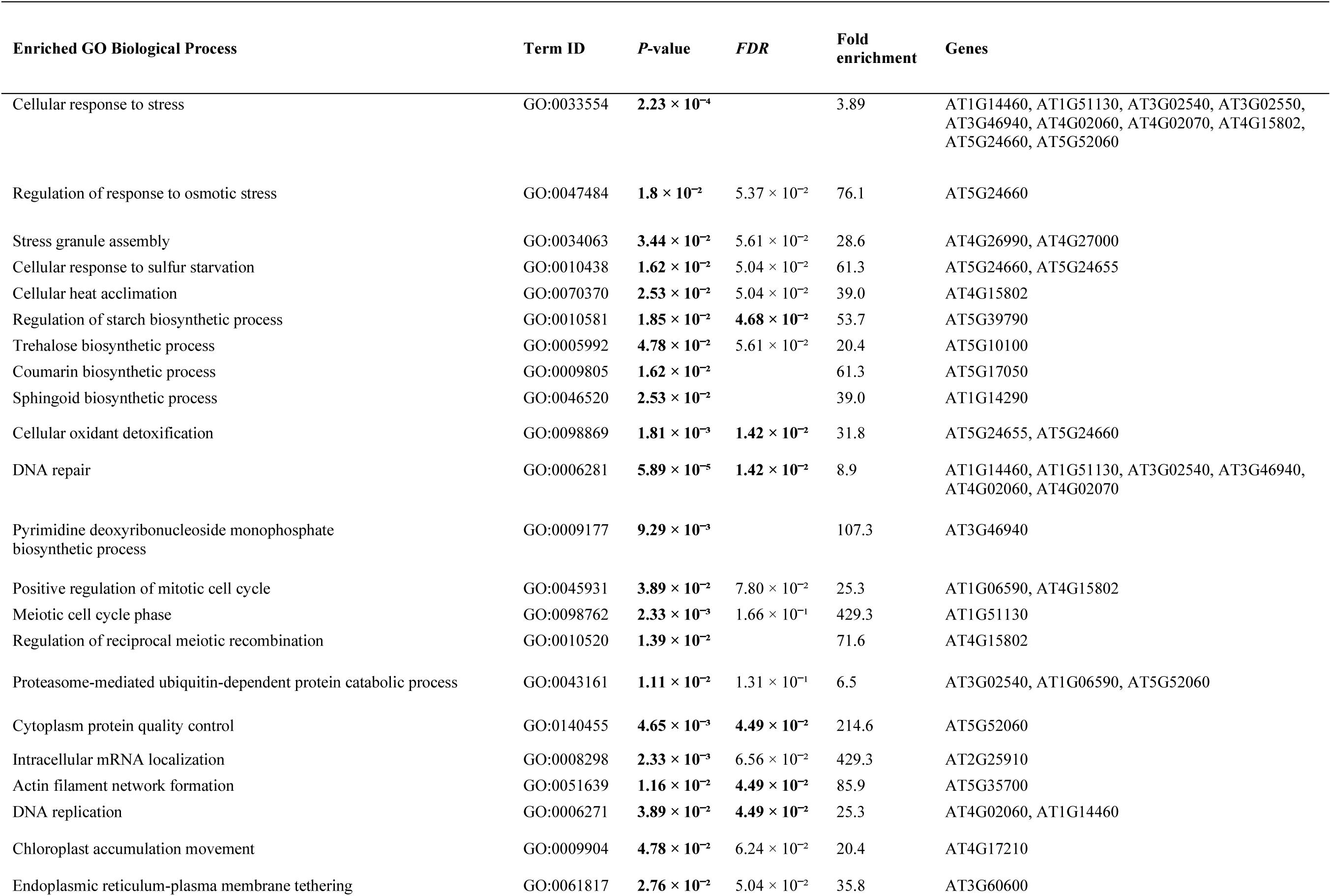
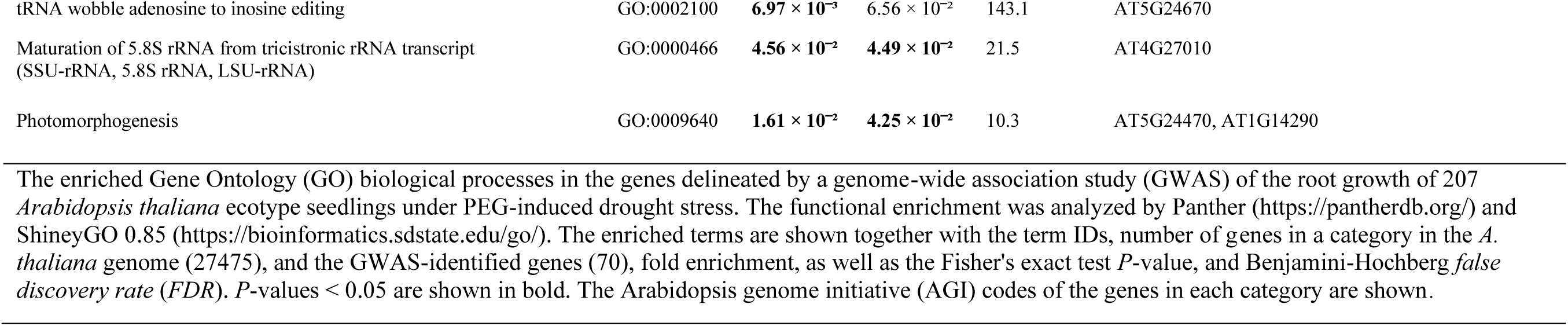
Functional enrichment of genes identified by genome-wide association study of drought tolerance in *Arabidopsis thaliana*.

| Enriched GO Biological Process | Term ID | P-value | FDR | Fold enrichment | Genes |
| --- | --- | --- | --- | --- | --- |
| Cellular response to stress | GO:0033554 | $2.23 \times 10^{-4}$ | | 3.89 | AT1G14460, AT1G51130, AT3G02540, AT3G02550, AT3G46940, AT4G02060, AT4G02070, AT4G15802, AT5G24660, AT5G52060 |
| Regulation of response to osmotic stress | GO:0047484 | $1.8 \times 10^{-2}$ | $5.37 \times 10^{-2}$ | 76.1 | AT5G24660 |
| Stress granule assembly | GO:0034063 | $3.44 \times 10^{-2}$ | $5.61 \times 10^{-2}$ | 28.6 | AT4G26990, AT4G27000 |
| Cellular response to sulfur starvation | GO:0010438 | $1.62 \times 10^{-2}$ | $5.04 \times 10^{-2}$ | 61.3 | AT5G24660, AT5G24655 |
| Cellular heat acclimation | GO:0070370 | $2.53 \times 10^{-2}$ | $5.04 \times 10^{-2}$ | 39.0 | AT4G15802 |
| Regulation of starch biosynthetic process | GO:0010581 | $1.85 \times 10^{-2}$ | $4.68 \times 10^{-2}$ | 53.7 | AT5G39790 |
| Trehalose biosynthetic process | GO:0005992 | $4.78 \times 10^{-2}$ | $5.61 \times 10^{-2}$ | 20.4 | AT5G10100 |
| Coumarin biosynthetic process | GO:0009805 | $1.62 \times 10^{-2}$ | | 61.3 | AT5G17050 |
| Sphingoid biosynthetic process | GO:0046520 | $2.53 \times 10^{-2}$ | | 39.0 | AT1G14290 |
| Cellular oxidant detoxification | GO:0098869 | $1.81 \times 10^{-3}$ | $1.42 \times 10^{-2}$ | 31.8 | AT5G24655, AT5G24660 |
| DNA repair | GO:0006281 | $5.89 \times 10^{-5}$ | $1.42 \times 10^{-2}$ | 8.9 | AT1G14460, AT1G51130, AT3G02540, AT3G46940, AT4G02060, AT4G02070 |
| Pyrimidine deoxyribonucleoside monophosphate biosynthetic process | GO:0009177 | $9.29 \times 10^{-3}$ | | 107.3 | AT3G46940 |
| Positive regulation of mitotic cell cycle | GO:0045931 | $3.89 \times 10^{-2}$ | $7.80 \times 10^{-2}$ | 25.3 | AT1G06590, AT4G15802 |
| Meiotic cell cycle phase | GO:0098762 | $2.33 \times 10^{-3}$ | $1.66 \times 10^{-1}$ | 429.3 | AT1G51130 |
| Regulation of reciprocal meiotic recombination | GO:0010520 | $1.39 \times 10^{-2}$ | | 71.6 | AT4G15802 |
| Proteasome-mediated ubiquitin-dependent protein catabolic process | GO:0043161 | $1.11 \times 10^{-2}$ | $1.31 \times 10^{-1}$ | 6.5 | AT3G02540, AT1G06590, AT5G52060 |
| Cytoplasm protein quality control | GO:0140455 | $4.65 \times 10^{-3}$ | $4.49 \times 10^{-2}$ | 214.6 | AT5G52060 |
| Intracellular mRNA localization | GO:0008298 | $2.33 \times 10^{-3}$ | $6.56 \times 10^{-2}$ | 429.3 | AT2G25910 |
| Actin filament network formation | GO:0051639 | $1.16 \times 10^{-2}$ | $4.49 \times 10^{-2}$ | 85.9 | AT5G35700 |
| DNA replication | GO:0006271 | $3.89 \times 10^{-2}$ | $4.49 \times 10^{-2}$ | 25.3 | AT4G02060, AT1G14460 |
| Chloroplast accumulation movement | GO:0009904 | $4.78 \times 10^{-2}$ | $6.24 \times 10^{-2}$ | 20.4 | AT4G17210 |
| Endoplasmic reticulum-plasma membrane tethering | GO:0061817 | $2.76 \times 10^{-2}$ | $5.04 \times 10^{-2}$ | 35.8 | AT3G60600 |
| tRNA wobble adenosine to inosine editing | GO:0002100 | **$6.97 \times 10^{-3}$** | $6.56 \times 10^{-2}$ | 143.1 | AT5G24670 |
| Maturation of 5.8S rRNA from tricistronic rRNA transcript<br>(SSU-rRNA, 5.8S rRNA, LSU-rRNA) | GO:0000466 | **$4.56 \times 10^{-2}$** | **$4.49 \times 10^{-2}$** | 21.5 | AT4G27010 |
| Photomorphogenesis | GO:0009640 | **$1.61 \times 10^{-2}$** | **$4.25 \times 10^{-2}$** | 10.3 | AT5G24470, AT1G14290 |

The enriched Gene Ontology (GO) biological processes in the genes delineated by a genome-wide association study (GWAS) of the root growth of 207 *Arabidopsis thaliana* ecotype seedlings under PEG-induced drought stress. The functional enrichment was analyzed by Panther (https://pantherdb.org/) and ShinyGO 0.85 (https://bioinformatics.sdstate.edu/go/). The enriched terms are shown together with the term IDs, number of genes in a category in the *A. thaliana* genome (27475), and the GWAS-identified genes (70), fold enrichment, as well as the Fisher's exact test *P*-value, and Benjamini-Hochberg *false discovery rate* (*FDR*). *P*-values < 0.05 are shown in bold. The Arabidopsis genome initiative (AGI) codes of the genes in each category are shown.

To understand the roles of the GWAS-delineated genes in drought, their expression levels were examined in Col-0 seedlings under PEG-induced drought stress. Eleven genes showed significant (*FDR* < 0.05) changes between drought and control conditions (Table 1). Among the up-regulated genes, *APRR5*, *EMB2788*, and AT4G26990 were induced 2.52-, 1.87-, and 1.50-fold, respectively, under drought. This suggests positive roles of these genes for seedling growth under drought stress. On the other hand, *LBD41* was strongly suppressed by 0.07-fold, followed by *TPPI*, *LSU2*, *UGT78D2*, and *haloacid dehalogenase-like hydrolase superfamily protein* (AT1G14310), suppressed 0.44-, 0.55-, 0.64-, and 0.66-fold, respectively, indicative of their negative roles in stress tolerance or cellular prioritization to conserve resources under drought. The expression levels were compared with public transcriptome datasets under osmotic stress (mannitol), high salt, and ABA in *A. thaliana* seedlings of similar stages (1-2 weeks) as used in this study (Table 1). The results indicated that out of the genes perturbed in our transcriptome, *APRR5* was consistently upregulated under osmotic and high salt stresses in all the studies, *ACD55.5* was upregulated by osmotic stress, *ML41Z*, *INCENP*, *BAG1*, *MCM7*, *RAD23-3* was suppressed by osmotic and high salt stresses, *PCMP-A4* and *DUT1* was suppressed by ABA, and *TPPI* was induced by ABA (log_2_FC > 1 for up and log_2_FC < 0.8 for down; *FDR* < 0.05). The results indicated that the genes identified by our GWAS are responsive to drought, salt, and the drought hormone ABA.

The GWAS-identified genes were further analyzed by examining their physically interacting proteins and co-expressing genes through network analysis, leading to clear insights into the biological pathways participating in root growth maintenance under drought. Altogether, 14 GWAS-detected genes were found to participate in eight interaction networks (Fig. 3a-h). The network ‘a’ included 43 densely interconnected genes, including four GWAS-identified genes: *DUT1*, *MCM7*, *MSH6*, and AT1G14460 (Fig. 3a; Table S2). Additional interacting genes included paralogs/subunits of *MCM*, *proliferating cell nuclear antigen* (*PCNA*), *origin recognition complex* (*ORC*), *DNA polymerase* (*POL*), *replication factor C* (*RFC*), *synthetic lethality with DPB11-1 5* (*SLD5*), *partner of SLD five 1* (*PSF1*), etc. These genes participated in the enriched processes ‘cellular response to stress,’ ‘DNA metabolic process,’ ‘DNA repair,’ and ‘cell cycle.’ Further, *MCM7* was identified as the most interconnected network hub gene (Fig. 3). The network ‘b’ included three GWAS-associated genes: *mitochondrial ribosomal protein L41Z* (*ML41Z*; AT5G39800), nicotinate/nicotinamide mononucleotide adenyltransferase (*NMNAT*; AT5G55810), and *nuclear RNA polymerase D4* (*NRPD4*; AT4G15950), with additional ribosomal proteins. *ML41Z* and *NMNAT* encode mitochondrial or chloroplast ribosome-associated proteins involved in translation within organelles, linking them to primary growth-enabling processes like energy metabolism, photosynthesis, and mitochondrial function. On the other hand, *NRPD4* encodes a non-catalytic subunit of RNA Polymerase IV and V, plant-specific RNA polymerases required for epigenetic gene silencing through RNA-directed DNA methylation (RdDM) to suppress mobile genetic elements in the heterochromatin, conferring genome stability (He et al., 2009). Network ‘c’ included *TAD* paralogs, *guanosine deaminase* (*GSDA*; AT5G28050), and *pyrimidine deaminase* (*PYRD-2*; AT4G20960) involved in tRNA modification, specifically in adenosine to inosine editing at the wobble position. Genes of network ‘d’ included *RAD23-3*, *peptide-N-glycanase 1* (*PNG1*; AT5G49570), and *Golgi-to-ER traffic-like protein* (*MDC12.19*; AT5G63220), functioning in the catabolic process of misfolded or incompletely synthesized proteins, facilitating their recognition and removal from the endoplasmic reticulum (ER). Network ‘e’ comprised *EMB2788*, *ARM repeat superfamily protein* (AT1G14300), and *RNA-binding (RRM/RBD/RNP motifs) family protein* (AT2G21440), regulating ribosome assembly, mRNA fate and function, and stress signaling through protein interactions, respectively, potentially linking stress signals to transcriptional or translational responses. All three genes are implicated in stress adaptation. EMB2788 is part of the ABA-regulated proteome, where ribosome biogenesis is modulated under stress (Zhang et al., 2019). Similarly, RRM proteins like AT2G21440 often respond to environmental cues by altering RNA metabolism. Network ‘f’ included four genes regulating mitosis through the anaphase-promoting complex/cyclosome (APC/C), with a GWAS-detected gene, *anaphase-promoting complex subunit 5* (*APC5*). Network ‘g’ genes, *2A phosphatase associated protein of 46 kD* (*TAP46*; AT5G53000), *chaperonin containing t-complex polypeptide-1 subunit 6-2* (*CCT6-2*; AT3G02530), and *thioredoxin superfamily protein* (*F18O22.30*; AT5G14240) encode chaperonins and their regulators, involved in protein folding. TAP46 is a regulatory subunit associated with a protein phosphatase 2A family protein in Arabidopsis and is a key effector in the target of rapamycin (TOR) signaling pathway, which regulates growth under normal and stress conditions (Ahn et al., 2011). TAP46 also plays a key role in ABA-mediated stress responses. TAP46 regulates PPX1, the catalytic subunit of protein phosphatase 4, which interacts with the CCT chaperonin complex for its own folding (Ahn et al., 2019). CCT6-2 is a subunit of the CCT complex, mediating folding of key cytosolic proteins, including tubulins, which are critical for microtubule formation and thus for cell structure and division (Ahn et al., 2019). AT5G14240, a thiol oxidoreductase, is also involved in protein folding (Lu & Holmgren, 2014). All three genes of the network ‘h’, *EMB1379* (AT5G21140), *methyl methanesulfonate sensitivity 21* (*MMS21*; AT3G15150), also known as *high ploidy 2* (*HPY2*), encoding a SUMO E3 ligase, and *NSE4A*, are components of the SMC5/6 complex, functioning in chromosome maintenance and DNA repair. To summarize, the network analysis demarcated different processes, viz., DNA repair, tRNA editing, protein quality control, and cell cycle regulation, among GWAS-identified genes and their interaction partners, functioning synergistically to maintain root growth under drought.

**Fig. 3.**
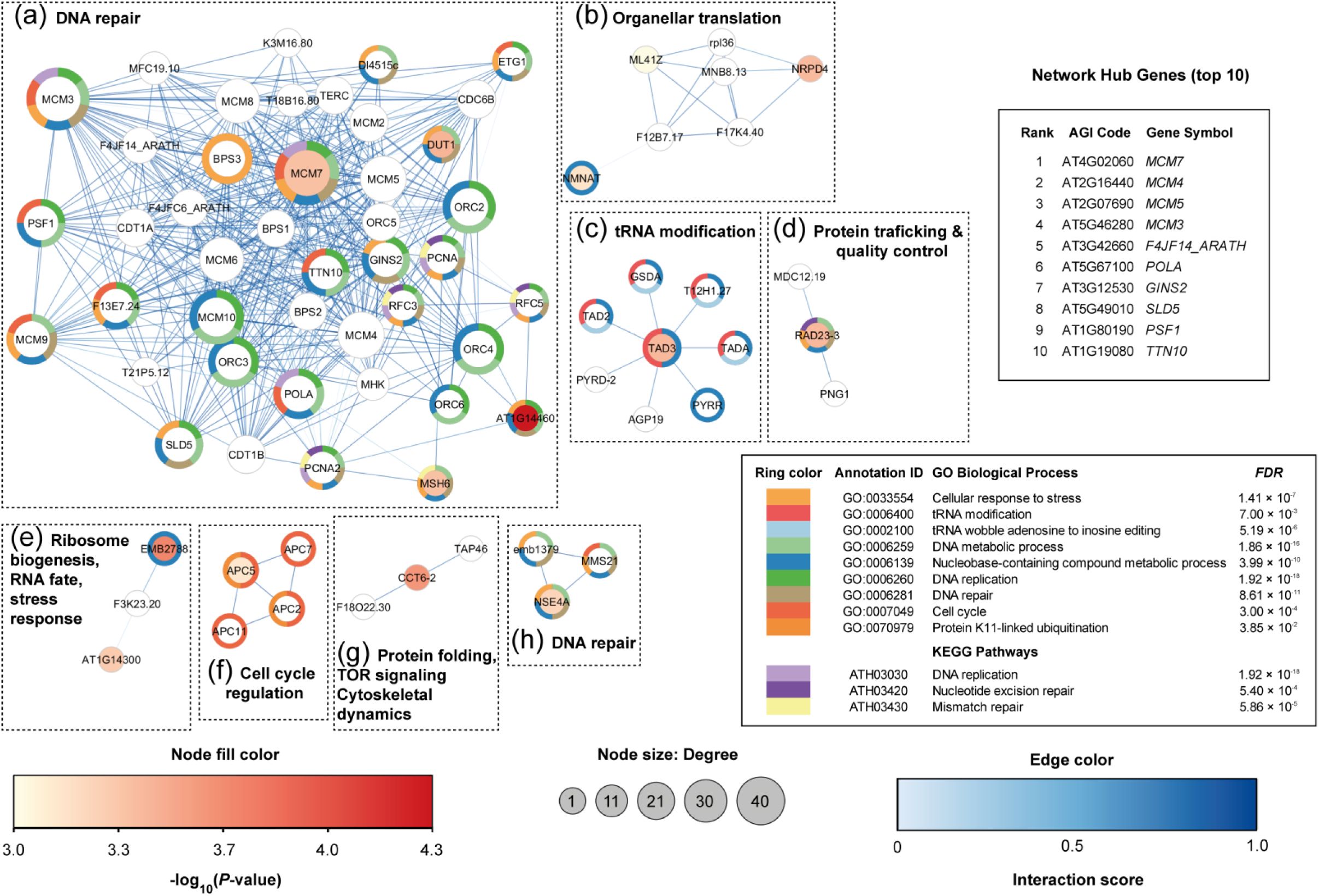
Network of genes identified by a genome-wide association study of drought tolerance in *Arabidopsis thaliana*. Genes within haploblocks of single-nucleotide polymorphisms (SNPs) with a *P*-value < 10^−3^ identified by the GWAS (see Table 1) were subjected to gene co-expression and protein-protein interaction network construction, using the STRING web server with a high confidence score (0.7). Circles indicate nodes, and lines indicate edges of the network. Node size represents the degree of prediction of relatedness of the genes by the software. Genes detected by the GWAS are colored in shades of red, indicating -log_10_(GWAS *P*-value), whereas the additional genes included, based on coexpression or physical interaction, are in white. Edge color in different blue shades indicates the interaction score. The rings around the nodes indicate enriched gene ontology (GO) biological processes and Kyoto Encyclopaedia of Genes and Genomes (KEGG) pathways of the genes in the network, with the color key shown in the table. The table shows enriched GO categories along with Fisher’s exact test *P*-values with Benjamini-Hochberg *false discovery rate* (*FDR*) correction.

Our GWAS identified *SKU5-similar 6* (*SKS6*; AT1G41830), belonging to the *SKS* multicopper oxidase family, implicated in cell wall remodeling, root growth, and oxidative stress response. This gene was down-regulated during a photosynthetic transition from C3 to crassulacean acid metabolism (CAM) under drought (Brilhaus et al., 2015). Interestingly, *SKS6* is a direct transcriptional target of our GWAS-identified TF, *ANAC094*, as indicated by public DAP-seq data in the Cistrome database.

### 3.4. GWAS-identified genes show expression-level and protein polymorphisms associated with drought tolerance

Polymorphisms in the coding sequences and upstream regulatory regions of the genes associated with the top 10 ranking GWAS-detected SNPs (Table 1) were analyzed with the denser SNP information for 142 out of the 207 ecotypes available in the 1001 genomes database. Seventeen genes showed various polymorphisms in the promoters and 5’-UTRs, along with several amino acid substitutions with ‘moderate’ impact, and eight genes showed ‘high’ impact polymorphisms, viz., stop codon gain and loss and frameshift mutations. Altogether, 14 genes exhibited amino acid polymorphisms, 13 displayed promoter variants, and nine had 5’-UTR variants (Table 3). Seven tolerant ecotypes gained a premature start codon due to a polymorphism at *Chr3:524543* on the 5’-UTR of the *regulator of chromosome condensation (RCC1) family protein* (AT3G02510). Similarly, start codon gain variants were observed at *Chr4:13558005* and *Chr4:13558113* on the 5’-UTR of *EMB2788*. *Raf-like mitogen-activated protein kinase kinase kinase 43* (*RAF43*; AT3G46930) showed a 5’-UTR splice region variant at *Chr3:17285818*. LAS calculations using both GLM and MLM for local genomic regions indicated significant associations of these polymorphisms with the drought tolerance phenotypes (Table 3). These included significant promoter variants in AT1G06690, AT1G14310, *F-box and associated interaction domains-containing protein* (AT1G14315), an *oligosaccaryltransferase* (AT5G02502), *ATHSBP*, *CCT6-2*, *RAF43*, *Fimbrin-like protein 2* (*FIM2*; AT5G35700), and *nuclease HARBI1-like protein* (AT5G35695), and significant 5’-UTR variants in AT3G02510, *EMB2788*, and a *glycosyl transferase family 1 protein* (AT4G01210).

**Table 3.**
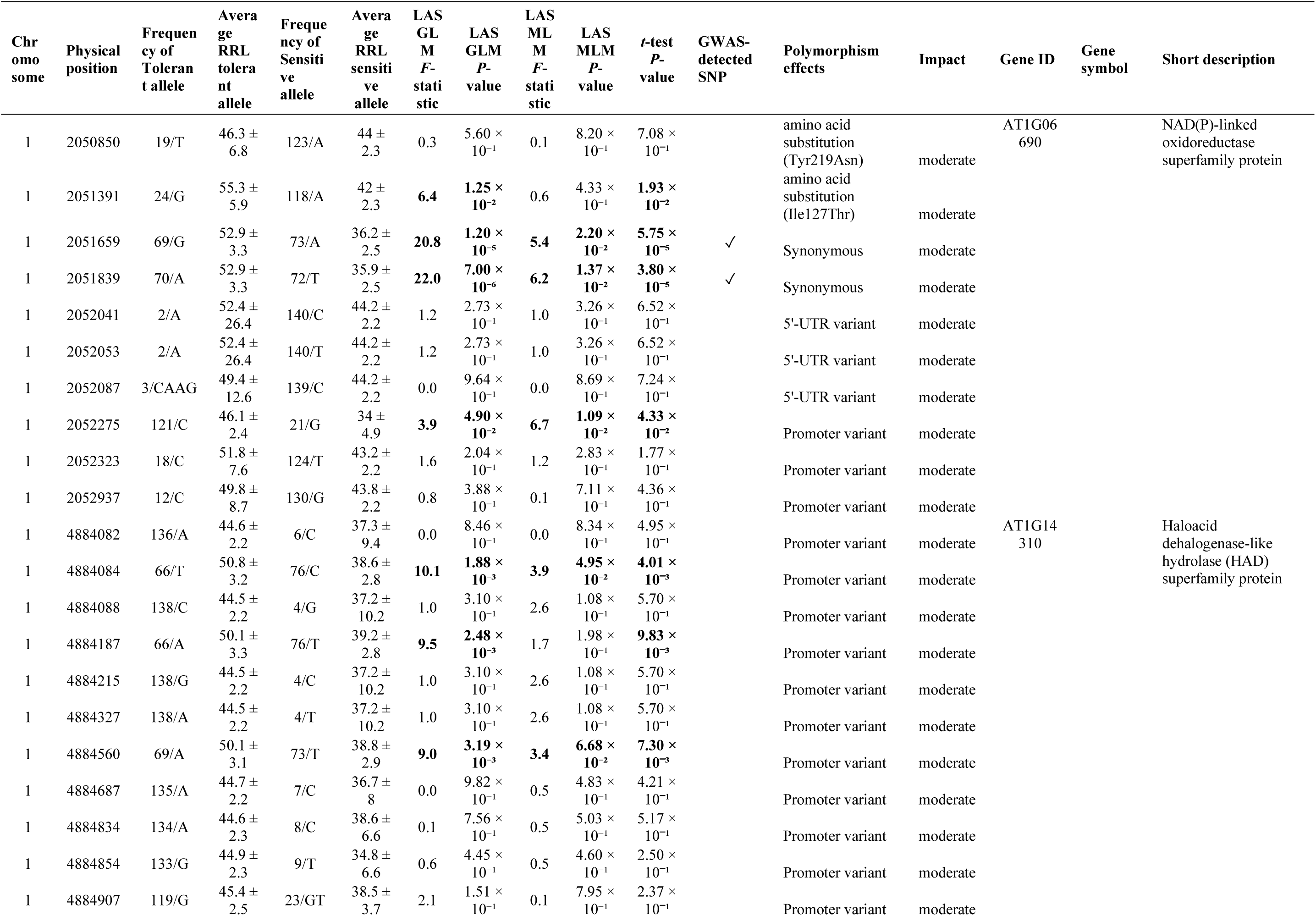
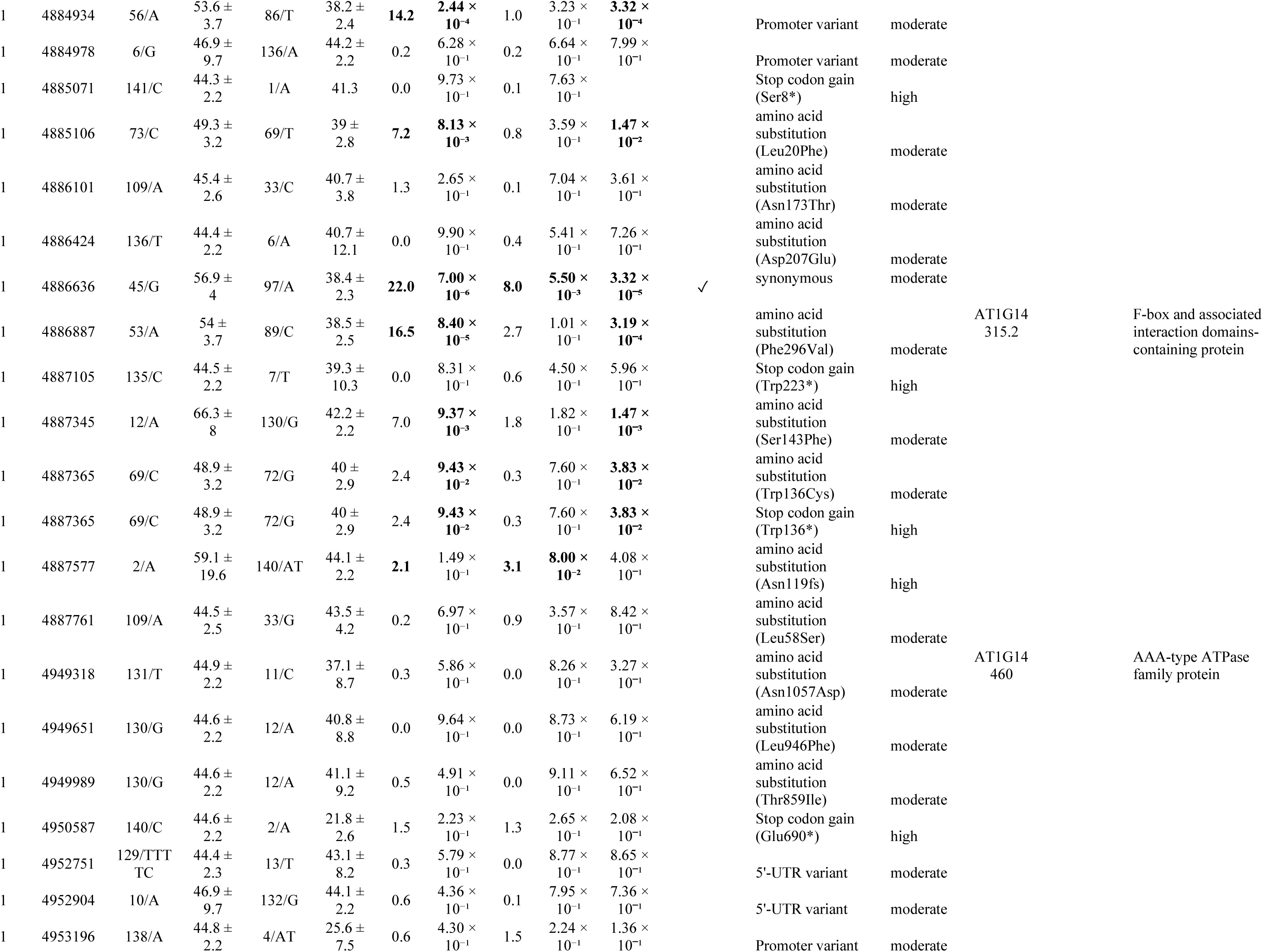
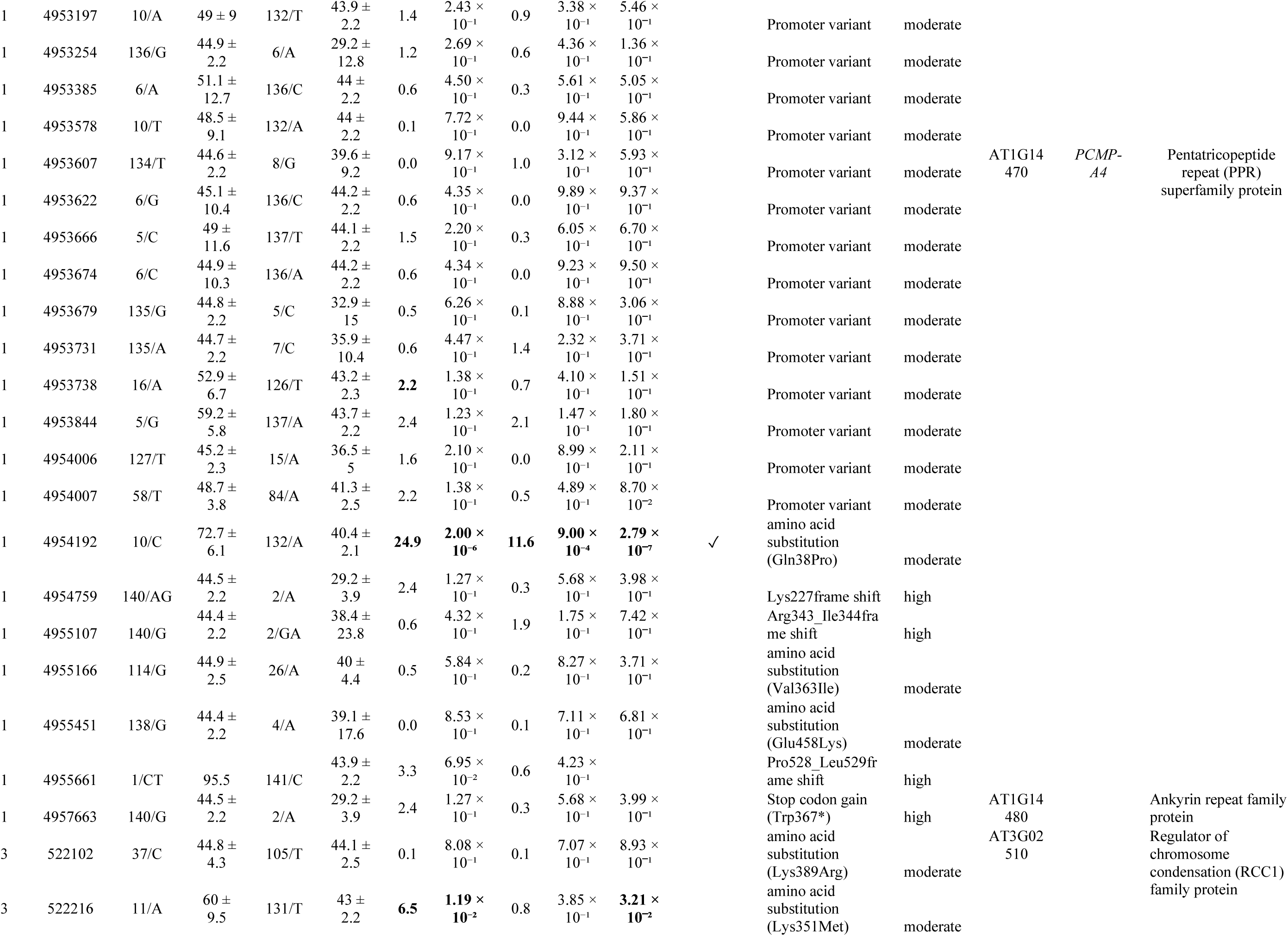
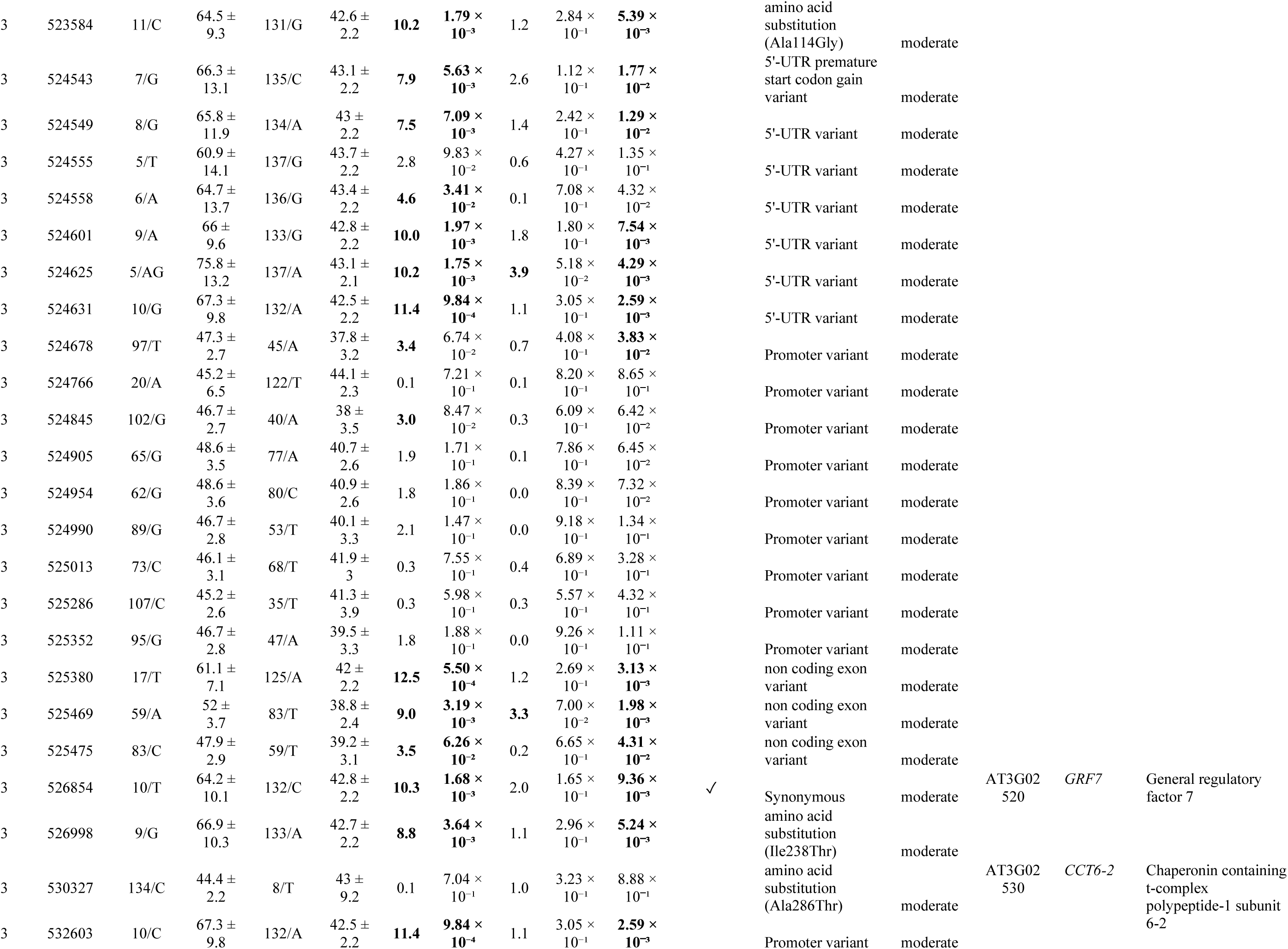
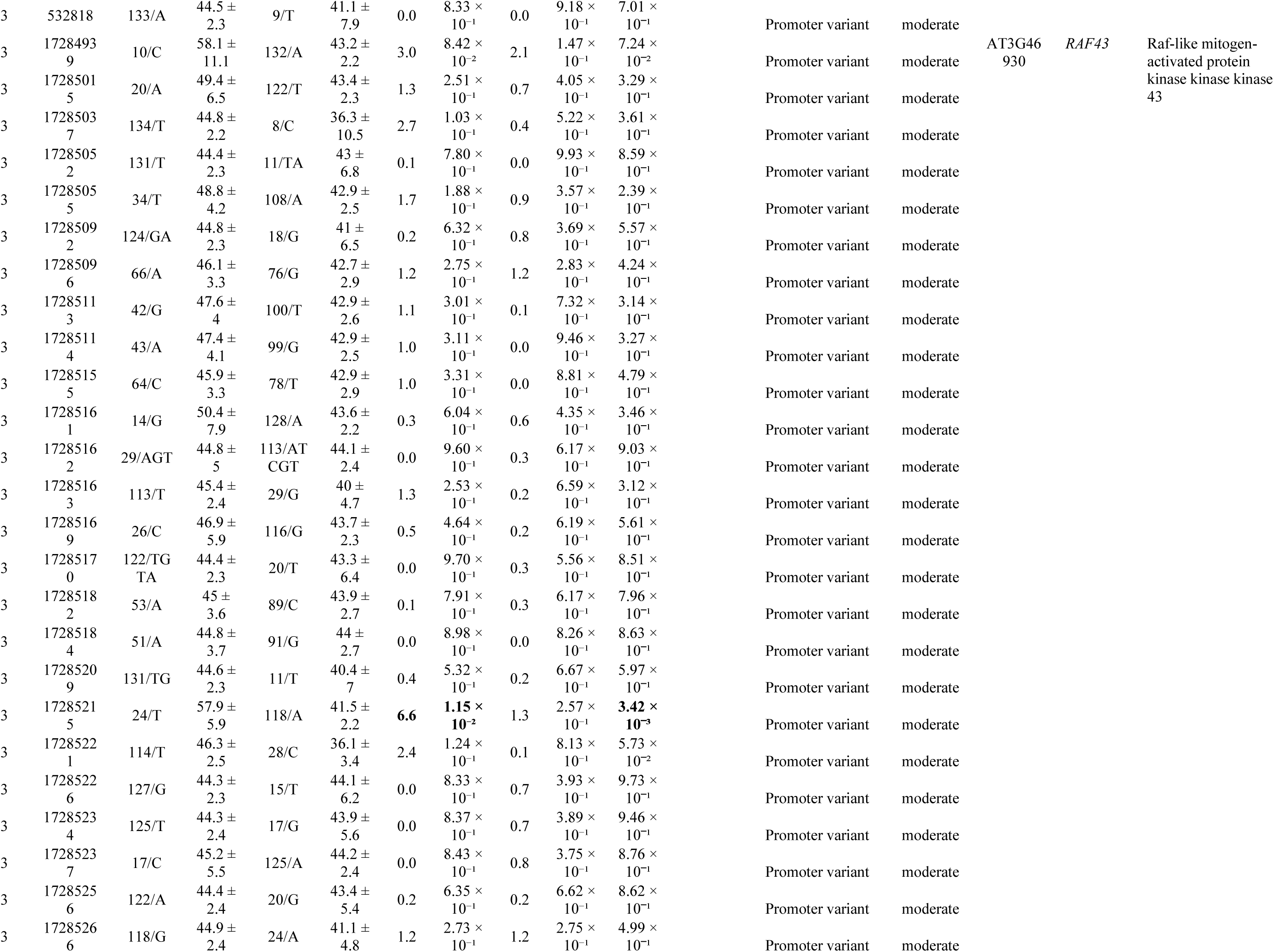
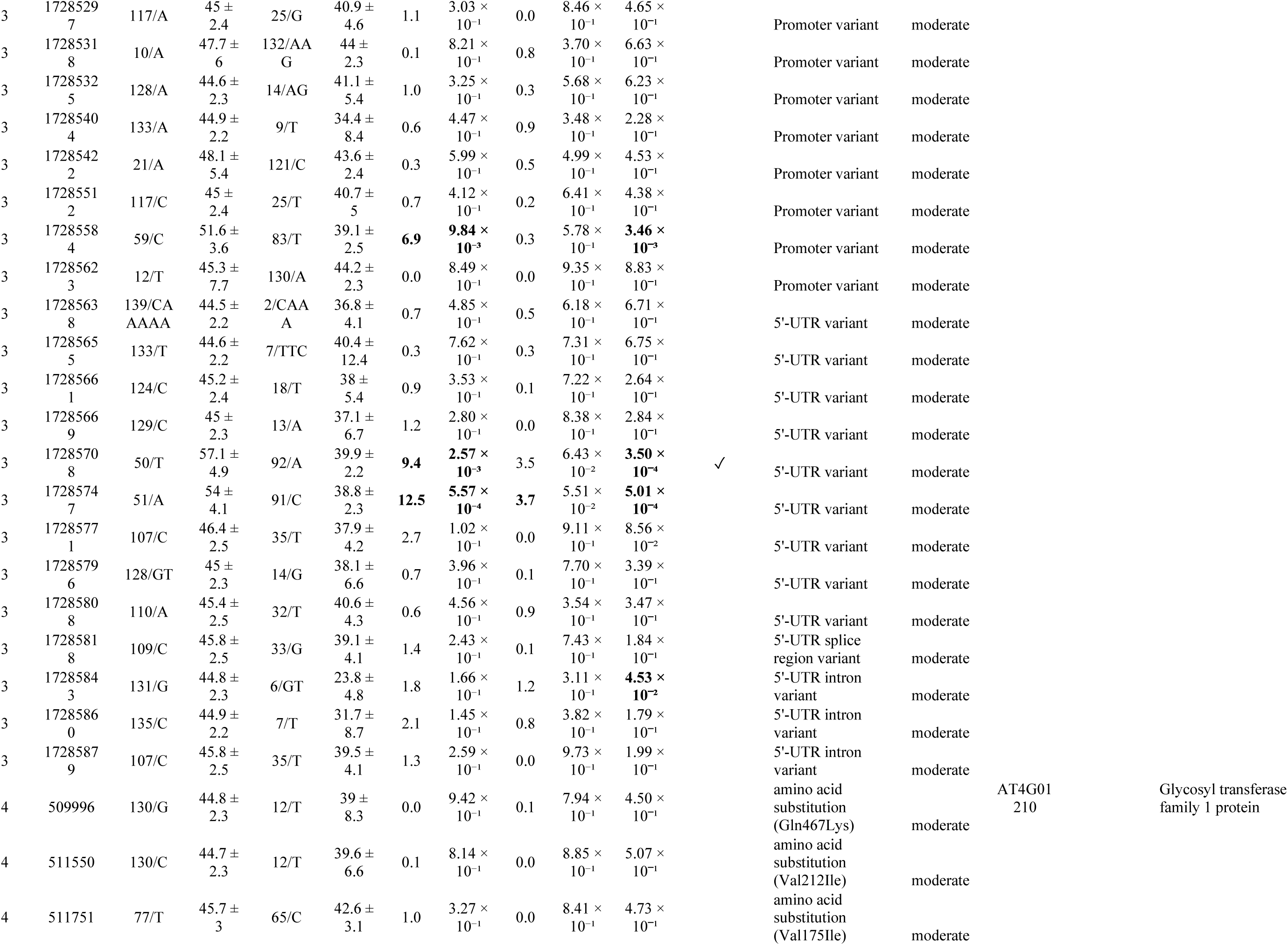
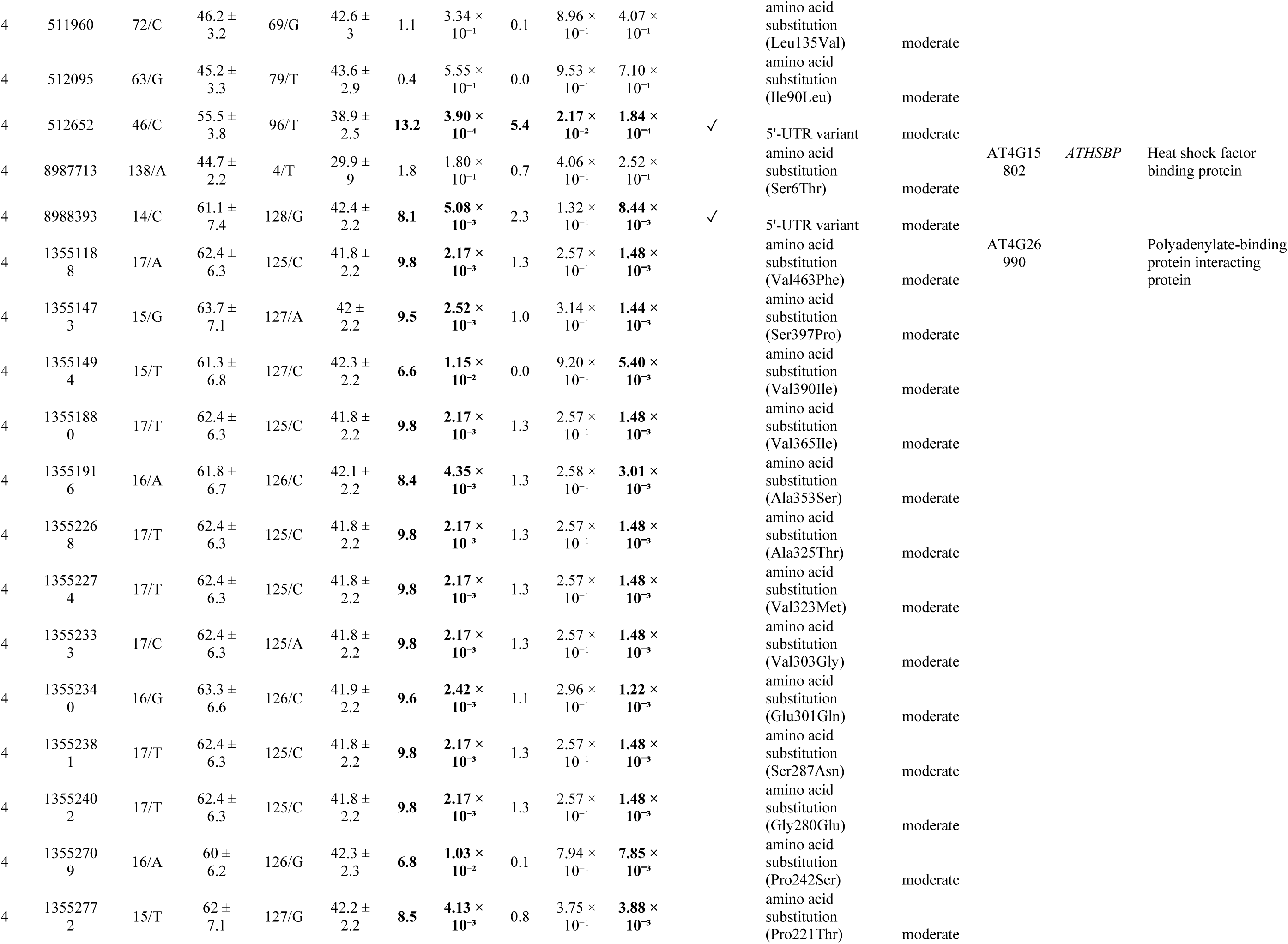
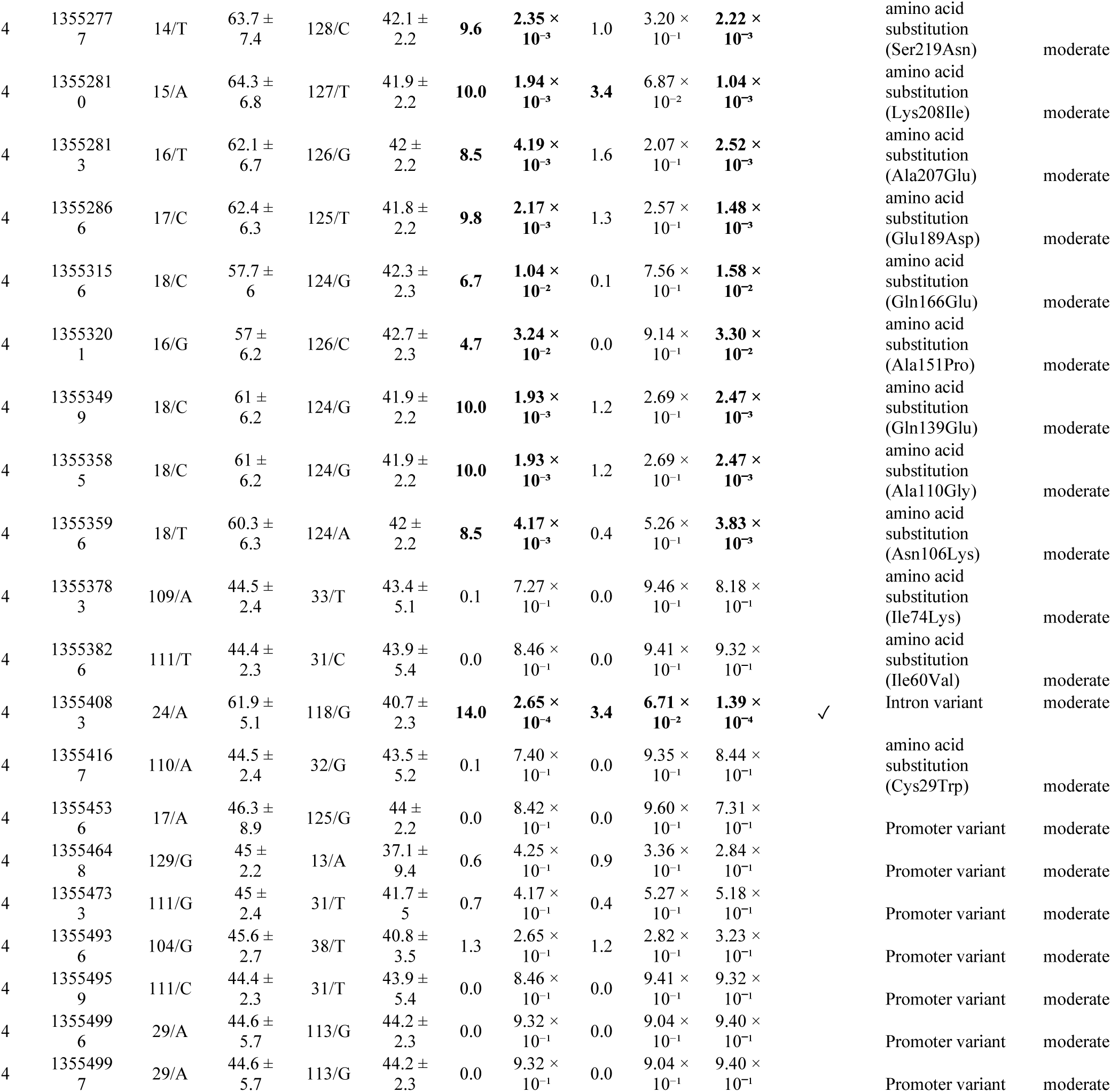
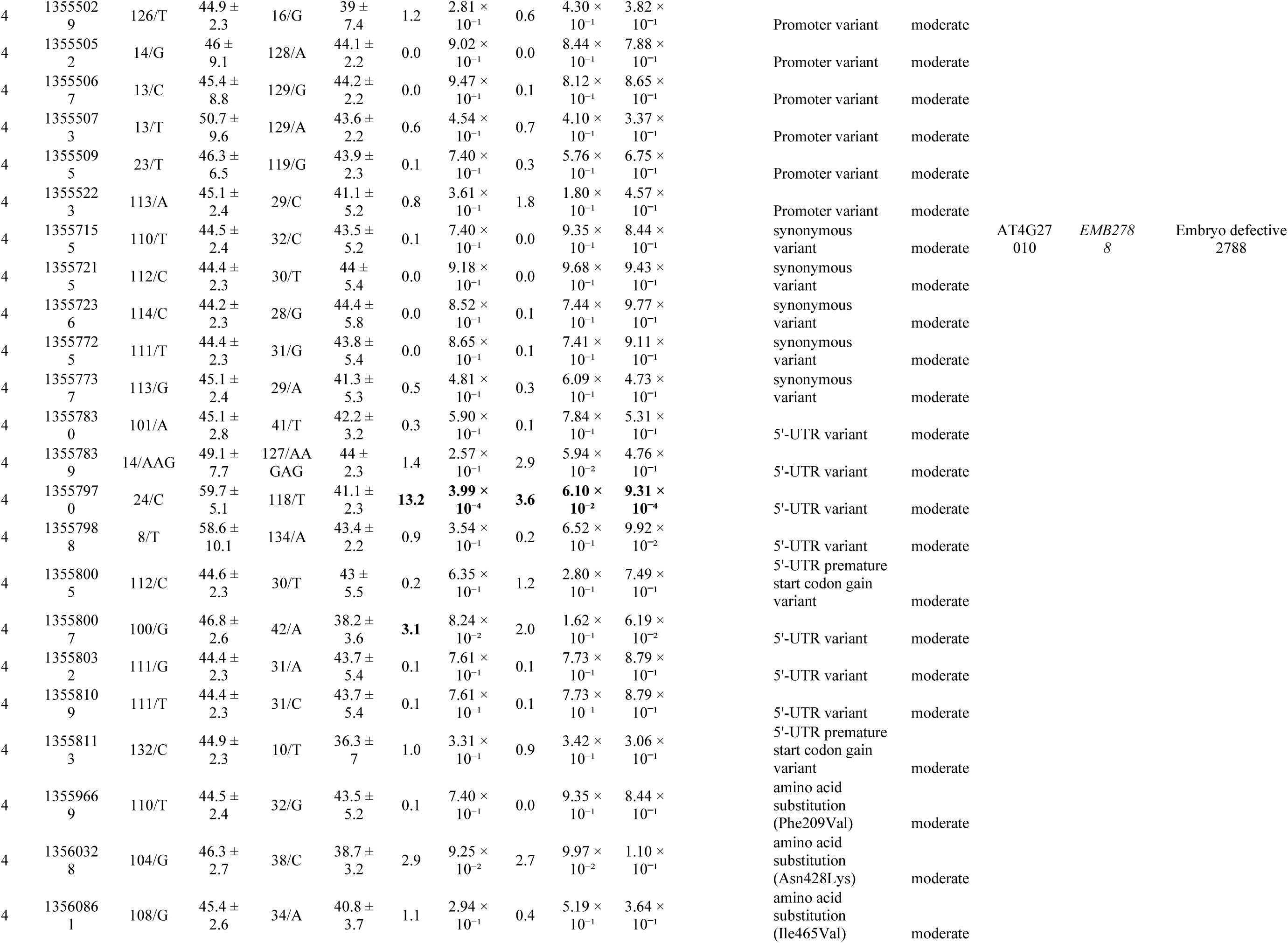
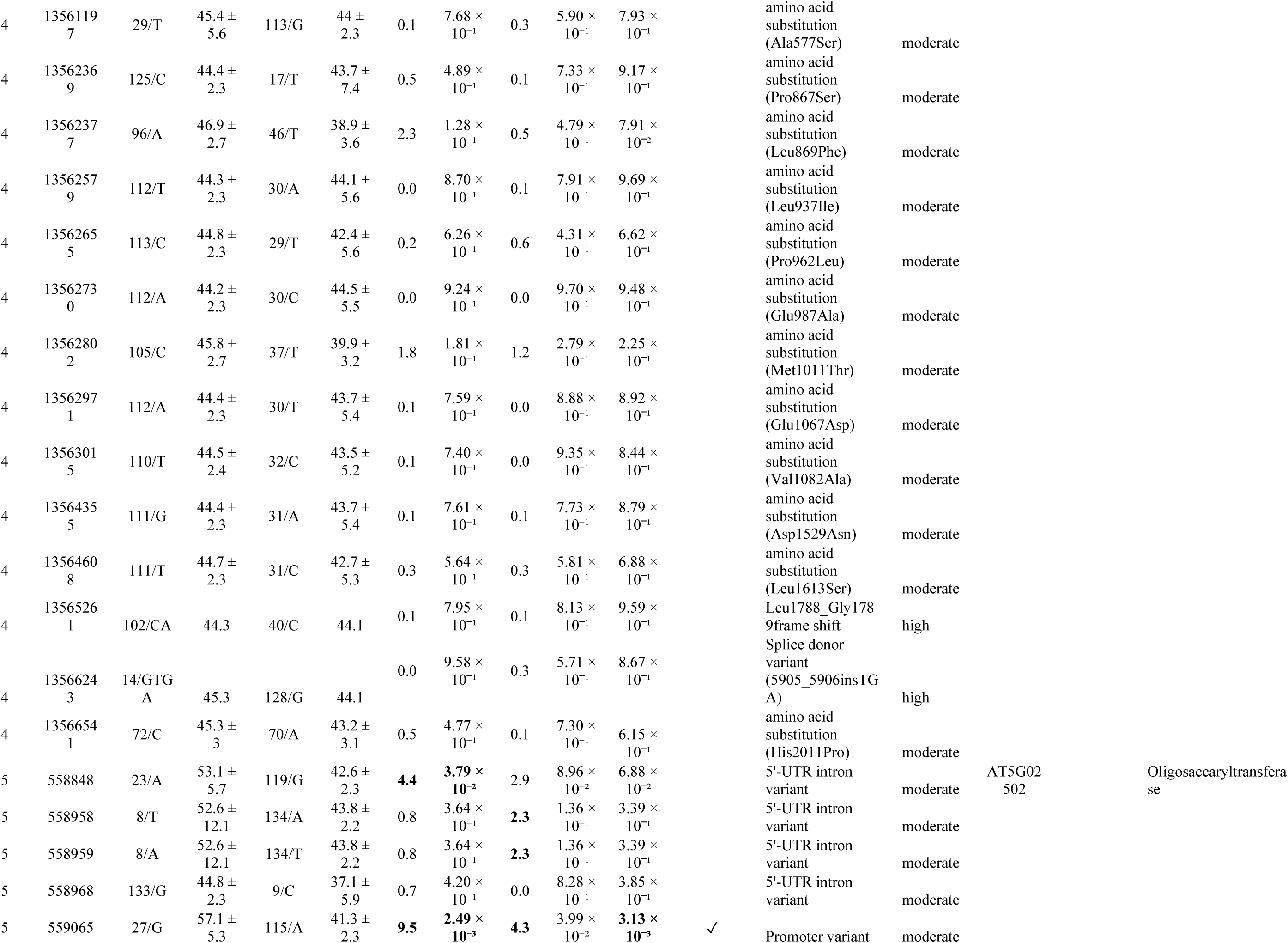
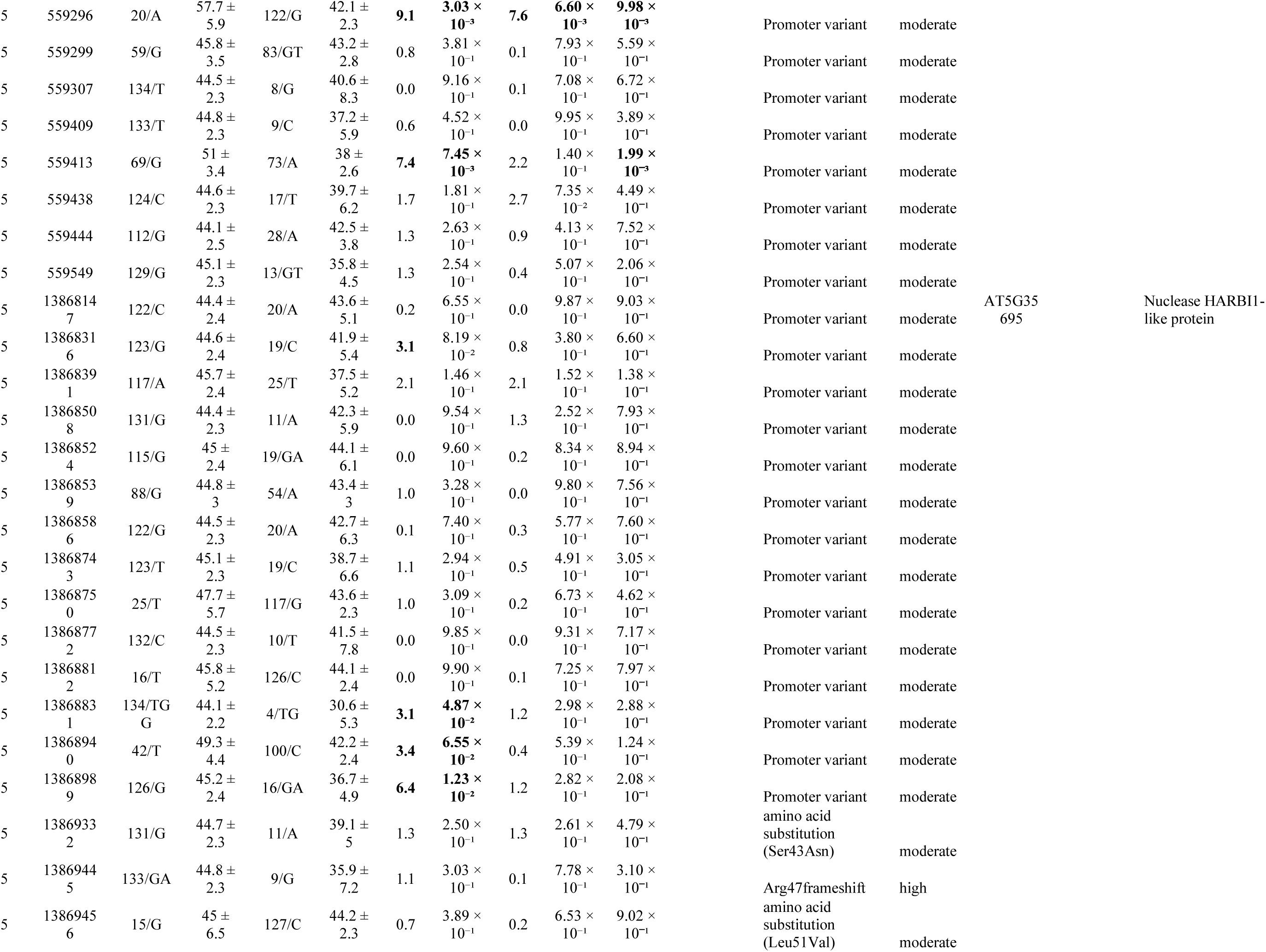
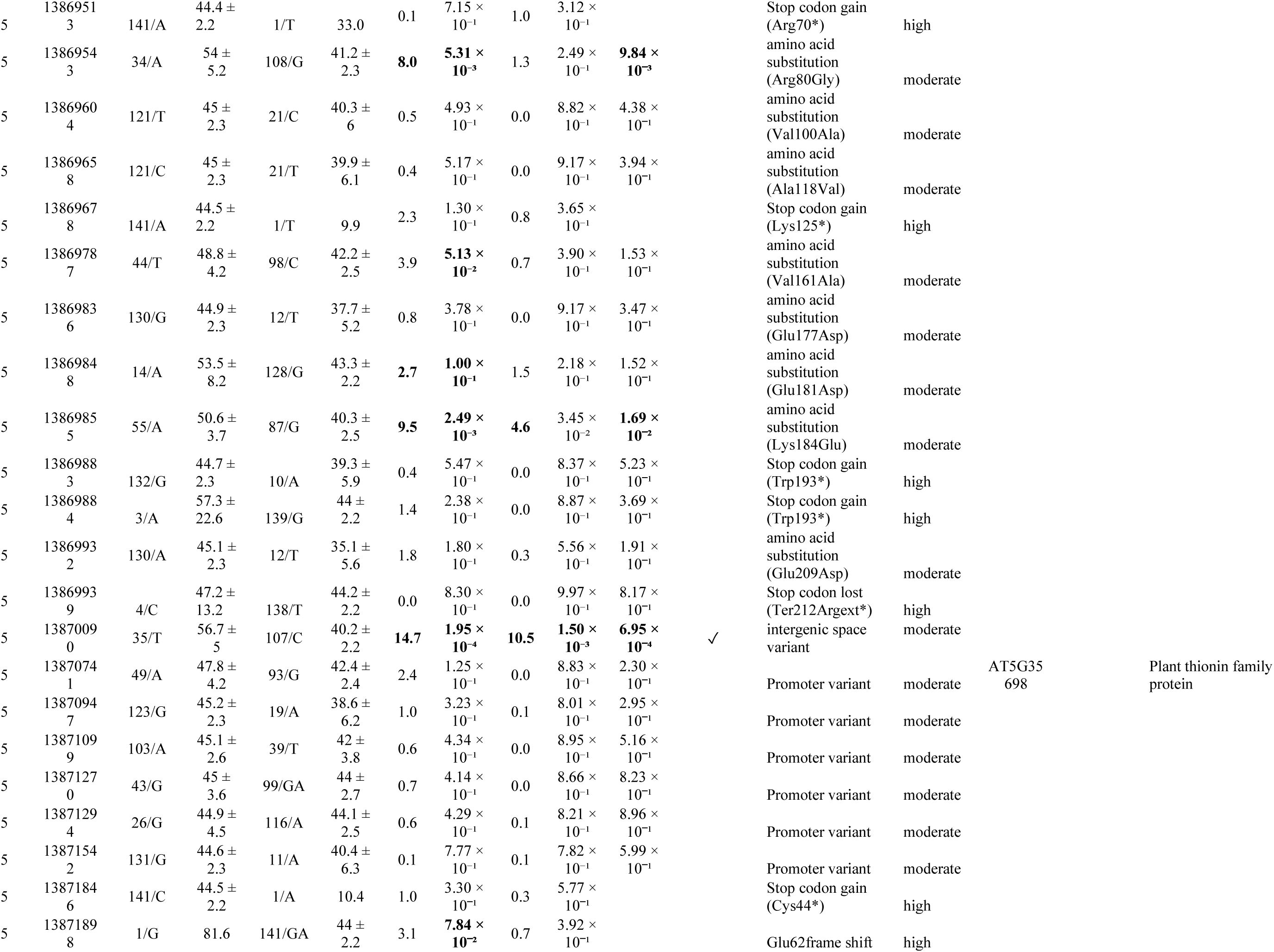
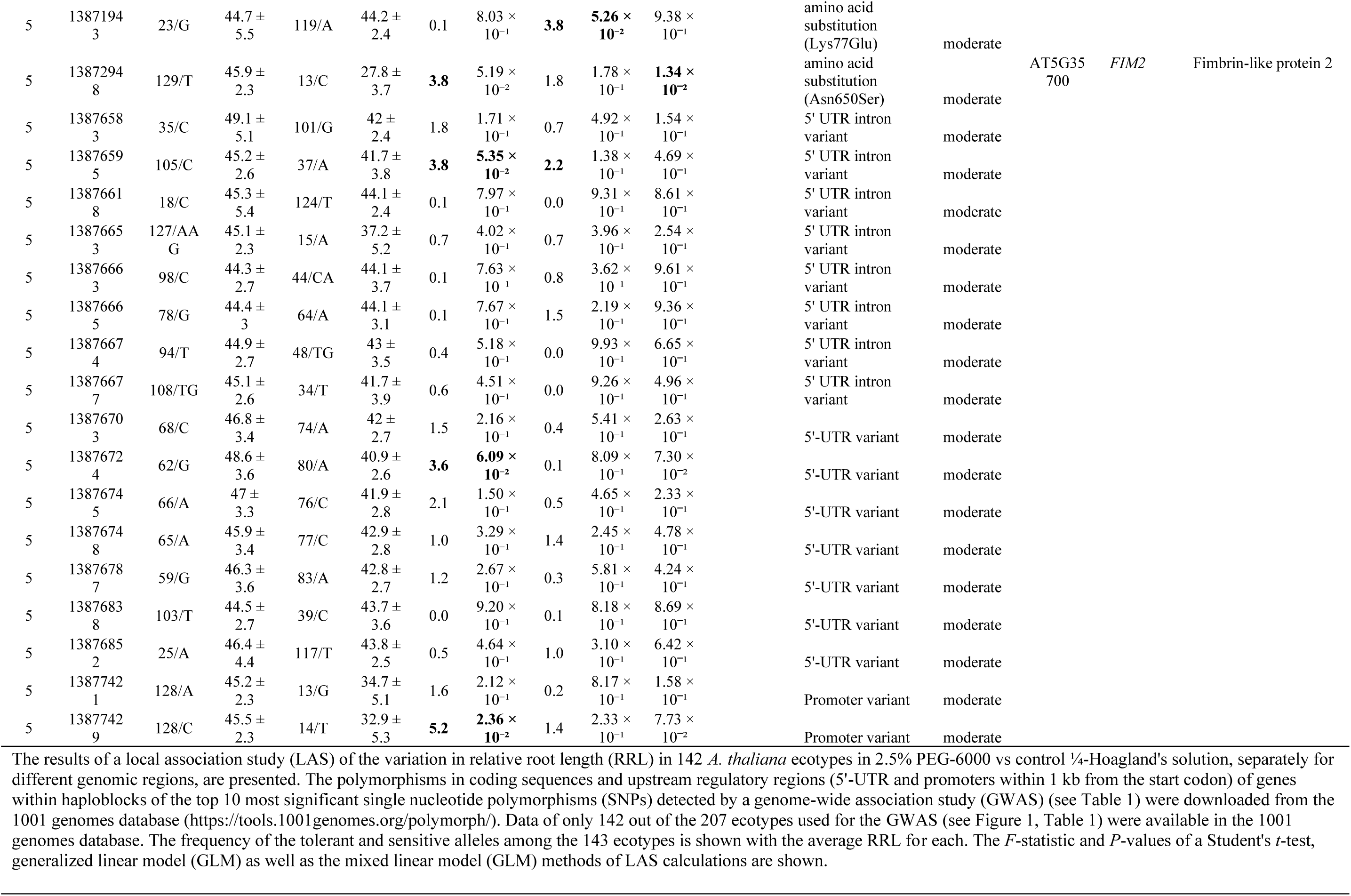
Local association study of root length variation in 142 worldwide *Arabidopsis thaliana* ecotypes under drought stress.

#### 3.4.1. GWAS-identified genes show expression level polymorphisms

We examined the variation in expression levels of genes nearest to the top 10 ranking SNPs (*P* < 10^−3.5^) with significant promoter polymorphisms detected by LAS (Table 3) among 40 randomly chosen ecotypes out of our diversity panel under drought. These included AT1G06690, AT1G14310, AT3G02520, AT5G02502, and AT5G35695 (for qPCR primer sequences, see Table S3). We could not generate suitable qPCR primers for the 521 bp gene AT5G02502. A significant association was detected at *Chr1:2052323* (*t*-test *P*-value = 0.03) and *Chr1:2052275* (*P* = 0.07) on the promoter of AT1G06690, with tolerant alleles associated with higher expression levels than sensitive ones. *Chr1:2052275* was found significantly associated (MLM *F* = 6.2, *P* = 0.001) with drought tolerance in the LAS (Table 3). The analysis of promoter *cis* elements indicated that *Chr1:2052323* (-284 bp from TSS) was localized within a poly A stretch 17 bp downstream to a MYC-binding site overlapping with an octamer “TGTCGTCT,” overrepresented (RAR = 2.3, Fisher’s test *P* = 0.006) in the promoters of genes induced by ABA (Fig. 4). *Chr1:2052275* (-236 bp from TSS) was localized at an octamer “CCAACTGA,” overrepresented (RAR = 3.1, *P* = 0.02) in the promoters of genes suppressed by ABA. This octamer also contained a MYB-binding site (MBS; CAACTG), a known drought-responsive *cis*-element. An opposite trend was observed at *Chr1:4884082* (*P* = 4.7×10^−5^) and *Chr1:4884687* (*P* = 4.8×10^−4^) on the promoter of AT1G14310, with higher expression associated with drought sensitivity in minor SNP alleles. *Chr1:4884082* (-967 bp from TSS) was localized at the octamer “ATTACACA,” overrepresented (RAR = 2.1, *P* = 9.7×10^−9^) in the promoters of genes suppressed under drought and ABA. *Chr1:4884687* (-362 bp from TSS) was at an octamer “CACCATAT,” overrepresented (RAR = 2.1, *P* = 0.02) in the promoters of genes induced under drought and 2 bp downstream to “CGTCACCA”, overrepresented (RAR = 2.2, *P* = 0.03) in the promoters of genes induced under ABA. Again, higher expression was associated with drought sensitivity for minor alleles of *Chr5:13868391* (*P* = 0.02) and *Chr5:13868743* (*P* = 0.009) on the promoter of AT5G35695 (Fig. 4). *Chr5:13868391* (-729 bp from TSS) was localized at an octamer “CGGATATG,” overrepresented in the promoters of genes suppressed under osmotic and high salt stresses. *Chr5:13868743* was 16 bp away from an MBS (Fig. 4). The promoter analysis indicated that SNPs within *cis* elements related to drought and osmotic stresses underlined the natural variation in gene expression of the GWAS-identified genes under drought.

**Fig. 4.**
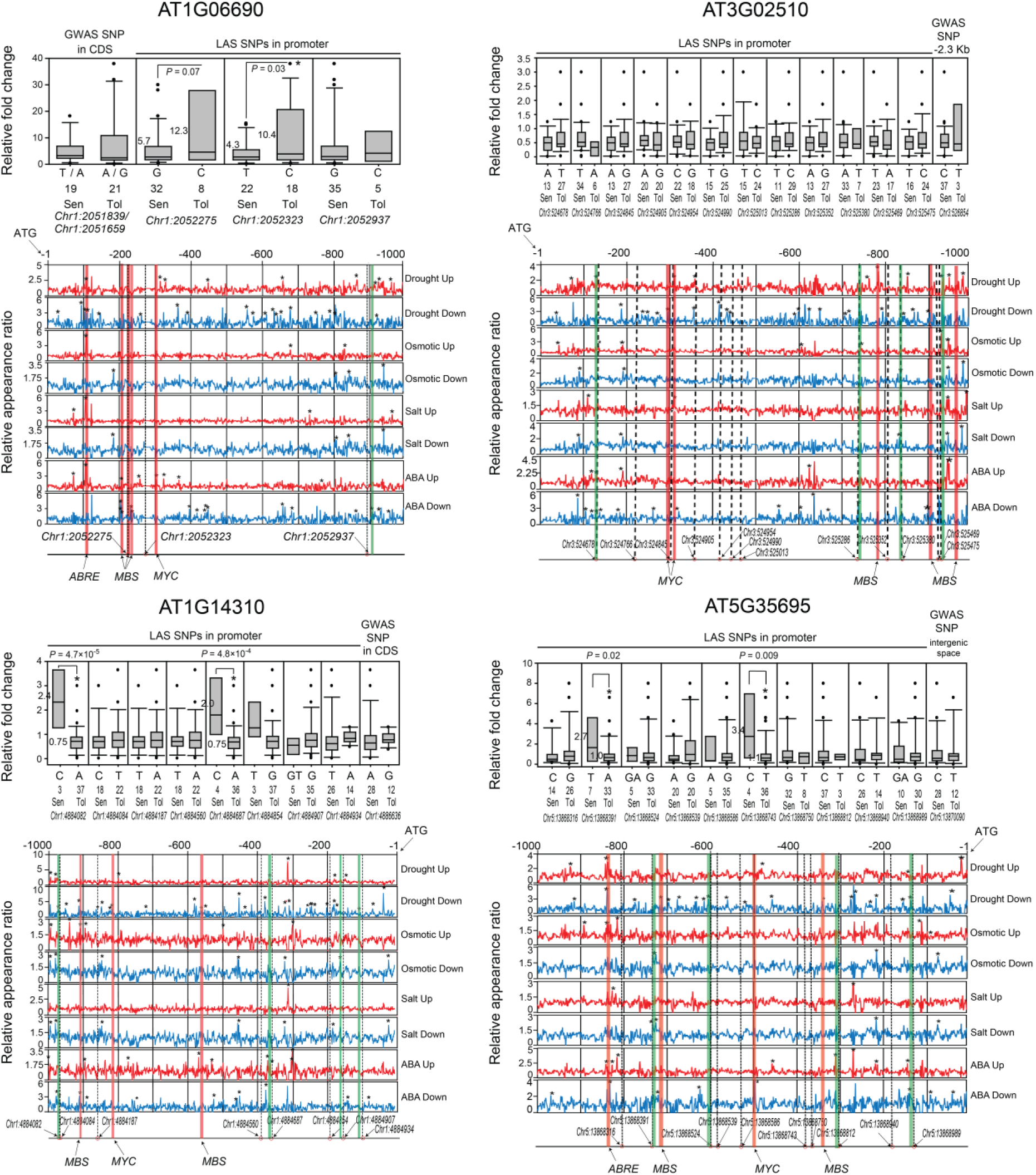
Expression level polymorphisms in GWAS-delineated genes. The expression level variations of genes nearest to the top 10 significant single-nucleotide polymorphisms (SNPs) in the genome-wide association study (GWAS) (Fig. 1, Table 1). The genes with significantly associated promoter polymorphisms in a local association study (LAS) conducted in 40 randomly selected ecotypes among the 207 ecotypes used in the GWAS are shown as box plots. Twenty plants of each ecotype were grown in ¼ Hoagland’s solution and treated with 2.5% PEG-6000 for 24 h. The expression levels in the roots were determined by quantitative real-time PCR and normalized with respect to *AtUBQ1* (AT3G52590) expression levels. The expression levels were further normalized relative to the Columbia (Col-0) ecotype using the 2^−ΔΔCt^ method. An average of three technical replicates is presented. In the X-axis of each plot, the ecotypes have been grouped according to the alleles at the respective SNP loci, mentioned below as chromosome numbers and physical positions in base pairs. “N” indicates the number of ecotypes containing the SNP allele. The asterisks indicate significant differences (Student’s *t*-test; *P* < 0.05). At significant positions, the numbers to the left of each box indicate the average relative fold change for ecotypes containing the SNP allele. Below the box plots, the curves show an overrepresentation of promoter elements (split into stretches of 8-nucleotides or octamers) in genes induced (up) or repressed (down) by drought, osmotic stress, high salt, and ABA (see Methods), along with mapping the promoter SNPs. The relative appearance ratio refers to the frequency of occurrence of an octamer in the promoters of stress-inducible or stress-repressed genes relative to the promoters of *Arabidopsis thaliana* genome-wide genes. Asterisks against the peaks indicate significant overrepresentation of octamers (Fisher’s test *P* < 0.05). Overrepresented octamers nearest to the promoter SNPs are indicated by green rectangles, and known drought-related *cis* elements from the PLANTCARE database by red.

#### 3.4.2. Polymorphic proteins show drastic structural variations

Altenb-2 (RRL 41.3%) from a moderate rainfall region (600-700 mm/year) gained a stop codon (Ser8*) in AT1G14310, leading to a truncated protein (Table 3; Table S4). Notably, the haplotype H7, comprising Kondara (92%), Sorbo (48.1%), Stepn-1 (48.2%), and Yeg-1 (9.7%) from low rainfall regions (<500 mm/year), and three, Ms-0 (19.2%), Sij-1 (29.8%), and Sij-2 (27.8%) from moderate rainfall regions, gained stop codons at *Chr1:4887105* (Trp223*) in AT1G14315. Another ecotype, Aitba-2 (31.3%) from a low rainfall region, gained a stop codon at *Chr1:4887365* (Trp136*) in AT1G14315 (Fig. 5). Kondara showed high drought tolerance; Sorbo and Stepn-1 moderate tolerance; while Yeg-1, Ms-0, Sij-1, and Sij-2 were sensitive to drought in our study (Table S4). Two ecotypes from Spain, Ll-0 (78.6%) and Bla-1 (39.4%), had a frameshift at *Chr1:4887577*, leading to an altered amino acid sequence from Asn119 and a premature stop downstream at 159. In AT5G35695, mostly sensitive ecotypes from moderate-to high-rainfall regions were found to gain stop codons at various amino acid positions, e.g., Rovero-1 (33%) at 70, Bsch-0 (9.9%) at 125, and Pog-0 (19.3%), An-1 (30.1%), CIBC-5 (41.6%), and Cnt-1 (48.2%) at 193. In haplotype H10, nine ecotypes from humid regions of Europe, viz., HKT2-4 (10.9%), Ag-0 (17%), Ema-1 (21.9%), NFA-8 (25.4%), Gy-0 (36%), Boot-1 (38%), Hh-0 (39%), NFA-10 (53.4%), Hn-0 (81.5%), had a frameshift at *Chr5:13869445* (Arg47), leading to a truncated AT5G35695 protein downstream at 63. In a pentatricopeptide repeat (PPR)-containing protein (PCMP-A4; AT1G14470), haplotype H3 consisting of two sensitive ecotypes, Pu2-23 (33%) and Zdr-1 (25.2%) had a frameshift at *Chr1:4954759* (Lys227) and haplotype H5 containing Sp-0 (14.6%) had a frameshift at *Chr1:4955107* (Agr343), leading to a short altered amino acid sequence followed by premature stop codons, leading to truncated proteins. Significantly associated (LAS *P* < 0.05) moderate-impact amino acid substitutions included Ile127Thr in AT1G06690; Leu20Phe in AT1G14310; Gln38Pro in PCMP-A4; Lys351Met and Ala114Gly in AT3G02510; Ile238Thr in AT3G02520; Arg80Gly, Val161Ala, Glu181Asp, and Lys184Glu in AT5G35695; and Lys77Glu in AT5G35698, which may influence the local protein structure with possible implications in protein functions (Table 3). A maximum of 22 amino acid substitutions were detected in AT4G26990, significantly associated with the drought tolerance phenotype in the LAS, indicating a prominent role of this protein in explaining the natural variation of seedling drought tolerance in *A. thaliana*.

**Fig. 5.**
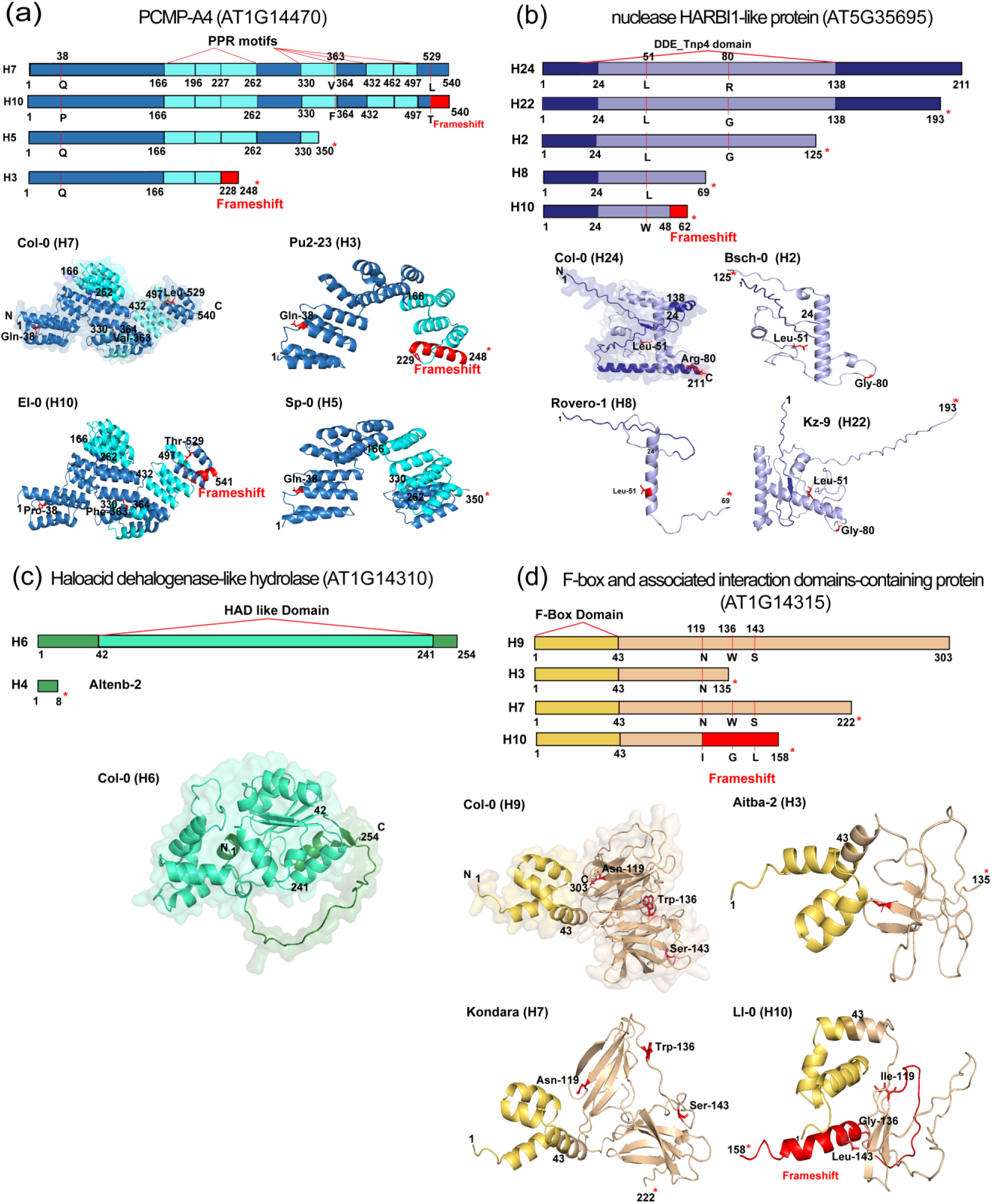
Drastic protein polymorphisms in GWAS-delineated genes. The high-impact protein polymorphisms n the top-ranking GWAS-delineated genes were identified by the POLYMORPH 1000 genome variants web ool (http://tools.1001genomes.org/polymorph/), and amino acid sequence haplotypes (denoted by H1, H2, etc) were constructed (Table S4). Representative haplotypes with drastic differences in protein structures are presented, together with the AlphaFold (https://alphafold.ebi.ac.uk/)-predicted three-dimensional structures. Defined InterPro (https://www.ebi.ac.uk/interpro/) domains are indicated in light colors. **(a)** pentatricopeptide epeat (PPR)-containing protein (PCMP-A4; AT1G14470) **(b)** nuclease harbinger transposase-like 1HARBI1)-like protein (AT5G35695) containing the DDE tnp4 domain **(c)** halo acid dehydrogenase (HAD)-ike hydrolase (AT1G14310), featuring the HAD domain **(d)** F-box, and associated interaction domains-containing protein (AT1G14315), displaying the F-box domain. Vertical red bars indicate amino acid ubstitution positions, asterisks indicate premature stop gains, and red patches indicate altered amino acid equences due to frameshifts.

BLAST searches of the candidate proteins against the PDB database did not provide any homologous proteins having solved structure passing the criteria of 40–100% sequence identity and 80–100% query coverage. Hence, we modeled the proteins *ab initio* using AlphaFold3 (Fig. 5, 6, Fig. S1). Haplotypes for further structure analysis (Table S4) were chosen based on amino acid substitutions that involve significant changes in amino acid properties, including polarity, charge, or drastic protein structural changes, caused by frameshifts and truncations. In the PPR protein PCMP-A4, the frameshift and premature stop codons were localized within its PPR motifs. The PPR motif of 35 amino acids is associated with RNA modification functions in organelles (Small & Peeters, 2000; Kotera et al., 2005). A similar motif, tetratricopeptide repeat (TPR), is known to facilitate protein–protein interactions that are essential for regulating stress signaling and transcriptional networks (D’Andrea, 2003). Loss and disruption of some of the PPR motifs in PCMP-A4 haplotypes H3 and H5 due to frameshift and truncation are likely to impair the protein’s structural integrity and ability to form functional complexes. In AT5G35695, a premature stop codon downstream of the DDE Tnp4 domain and several substitutions were detected. The DDE Tnp4 domain, derived from transposases, is generally associated with DNA cleavage and modification functions and may play a role in genome regulation during stress responses (Hickman et al., 2010). Truncation beyond this domain could disrupt the catalytic or structural support regions of this protein, potentially compromising DNA-related processes required for drought adaptation. For the HAD-like hydrolase (AT1G14310), several non-synonymous substitutions were located within the core of the HAD domain, including Asn119, Trp136, and Ser-143. HAD domains are critical for phosphatase activity and are implicated in stress-responsive metabolic regulation. *AtHAD1*, a related *Arabidopsis* HAD-like phosphatase, has been shown to repress ABA responses (Lee et al., 2022). The observed mutations may modulate phosphatase activity or substrate specificity, thereby influencing drought-related signaling. In AT1G14315, a frameshift mutation was identified within the F-box domain, which is essential for forming Skp1-Cullin-F-box (SCF) E3 ubiquitin ligase complexes, controlling the degradation of regulatory proteins involved in hormonal and stress pathways. Mutations that disrupt this domain could impair substrate recruitment or complex stability, affecting hormone signaling such as ABA, which plays a critical role in drought responses (Lechner et al., 2006). Altogether, these domain-disrupting polymorphisms, including premature stops and frameshifts, suggest significant loss-of-function or altered functionality in different *A. thaliana* ecotypes.

**Fig. 6.**
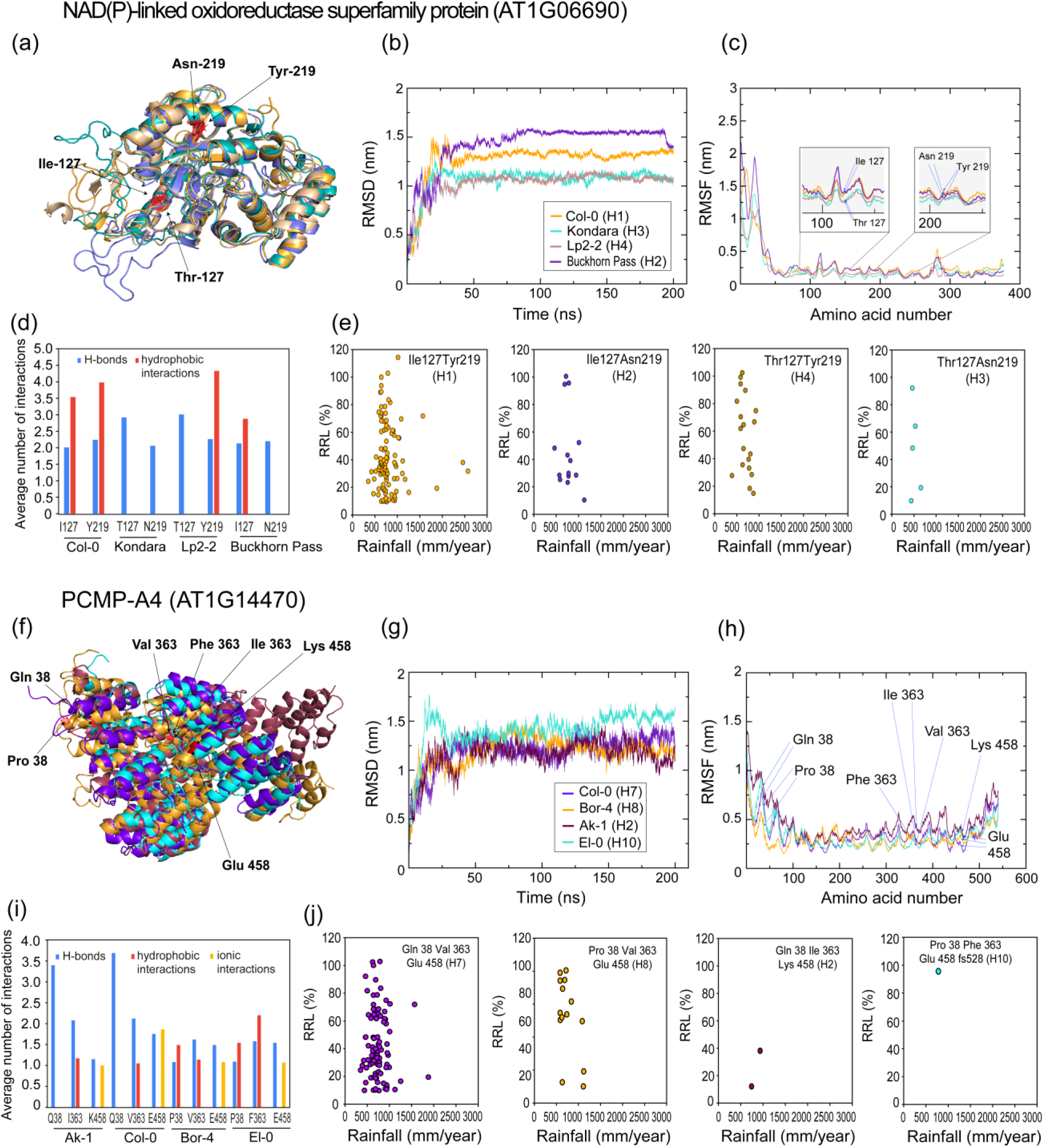
Effect of amino acid substitutions on predicted protein molecular dynamics. The moderate-impact amino acid substitutions in two top-ranking GWAS-delineated genes, NAD(P)-linked oxidoreductase uperfamily protein (AT1G06690) and pentatricopeptide repeat (PPR)-containing protein (PCMP-A4; AT1G14470), were identified by the POLYMORPH 1000 genome variants web tool http://tools.1001genomes.org/polymorph/), and amino acid sequence haplotypes (denoted by H1, H2, etc) were constructed (Table S4). AlphaFold (https://alphafold.ebi.ac.uk/)-predicted three-dimensional structures of epresentative haplotypes with the most amino acid substitutions are presented after molecular dynamics (MD) imulation as structural superpositions (**a, f**). Root mean square deviation (RMSD) plots over the simulation ime show how much the protein’s backbone structure deviates from its initial conformation, reflecting the overall structural stability and changes during the MD simulation (**b, g**). Backbone Root Mean Square Fluctuation (RMSF) profiles plotted across residue numbers highlight the flexibility and dynamic behaviour of ndividual residues during the MD simulation **(c, h).** Note that the color key for the curves is the same between **b**) and (**c**), and between (**g**) and (**h**). Predicted intramolecular interactions (hydrogen bonds, hydrophobic nteractions, and ionic interactions) between the polymorphic amino acids with their neighbouring residues are hown as bar graphs (**d, i**). The average number of interactions was calculated as the total interaction count divided by the number of frames in which the interaction was observed. Climate association of protein haplotypes is shown as correlation plots of the drought-tolerance phenotype (relative root length; RRL%) versus average annual rainfall in mm (**e, j**).

#### 3.4.3. Amino acid substitutions underlie the differential dynamics of polymorphic proteins

We performed MD simulations on representative protein haplotypes (Table S4) to better understand the effect of the amino acid polymorphisms on protein structures. For amino acid substitutions causing a change in polarity or charge of the side chains, MD simulations revealed transient variations in local protein conformations along with differential hydrogen bonding, hydrophobic, and ionic interactions between residues. In AT1G06690, the amino acid position 127 showed a change from Ile (nonpolar) to Thr (polar), and 219 from Asn (polar, aliphatic) to Tyr (polar, aromatic) (Fig. 6a; Table S4). The NADP^+^ cofactor-binding site was located between residues 263-273, spatially distant from the polymorphism positions 127 and 219. The root mean square deviation (RMSD) plot showed that the protein of haplotype H2 (Buckhorn Pass) exhibited the highest deviation over time, indicating greater conformational changes and the least structural stability. Proteins of haplotypes H1 (Col-0) and H3 (Kondara) showed moderate fluctuations, with the Col-0-type protein stabilizing at slightly higher RMSD values than the Kondara-type. The H4 (Lp2-2)-type protein maintained the lowest RMSD values throughout the simulation, suggesting higher structural stability (Fig. 6b). The root mean square fluctuation (RMSF) plot indicated that all ecotypes display higher flexibility at the N-terminal region due to the presence of non-regular secondary structures. The Buckhorn Pass-type protein exhibited the highest fluctuations at multiple regions, consistent with its unstable RMSD profile. In contrast, the Lp2-2-type maintained relatively low fluctuation across the protein. The Col-0 and Kondara-type proteins showed moderate fluctuations, with the Kondara-type being slightly more flexible in certain segments (Fig. 6c). These results suggest that even minor variations at positions 127 and 219 influence the global and local dynamics of AT1G06690. The Thr127 and Asn219 combination appears to increase protein flexibility as observed in Kondara, while Thr127 and Tyr219 in Lp2-2 stabilize the protein structure. The Ile127 and Asn219 combination in Buckhorn Pass destabilizes the conformation, whereas the Ile127 and Tyr219 combination in Col-0 maintains moderate stability (Fig. 6c). Notably, Kondara and Lp2-2, both carrying Thr at position 127, exhibit the lowest RMSD values, suggesting that the presence of Thr at this site contributes to enhanced structural stability. For position 219, the hydrogen bonding pattern is conserved across all haplotypes. In contrast, at position 127, Kondara and Lp2-2-type proteins consistently contain more hydrogen bonds due to the presence of Thr, whereas Col-0 and Buckhorn Pass-types have fewer due to Ile (Fig. 6d). Interestingly, the Thr127 and Asn219 combination was associated with low rainfall regions (Fig. 6e).

In AT1G14470, the GWAS and LAS-associated position Gln38Pro represents an open-chain to a ringed amino acid substitution. Other substitutions include Ile363Val, a conservative polymorphism, Ile363Phe, a shift from aliphatic to aromatic amino acid, and Glu458Lys, a change from a negative to a positively charged residue (Table S4). MD simulations revealed that substitutions at residue 38 have a greater influence on the structural dynamics of AT1G14470 than the other substitutions, consistent with the significant association of residue 38 with the drought tolerance phenotype (Table 3). In the RMSD trajectories, the H10 (El-0)-type protein with Pro38, Phe363, and Glu458 showed the highest backbone deviations, followed by the H8 (Bor-4)-type protein with Pro38, Val363 and Glu458, while the H2 (Ak-1)-type protein with Gln38, Ile363, and Lys458 and H7(Col-0) with Gln38, Val363, and Glu458 maintained comparatively lower RMSD, indicating slightly greater structural stability (Fig. 6). RMSF profiles, however, indicated that the H2 (Ak-1) and H7 (Col-0)-type proteins displayed the highest N-terminal flexibility around Gln38, followed by the Pro38-containing haplotypes. In Ak-1, the RMSD remained low, indicating global stability, while the high RMSF reflects localized flexibility in specific regions (Martínez 2015). The intramolecular interaction analysis further revealed that Gln38 in H2 and H7 engaged in multiple and persistent hydrogen bonds compared to the fewer hydrogen bonds observed in H8 (Bor-4) and H10 (El-0). In contrast, residues 363 and 458 exhibited fewer and predominantly transient hydrogen bonds that were similar across all haplotypes, underscoring the greater contribution of residue 38 to the structural dynamics.

For AT1G14310, the LAS-associated (GLM) position Leu20Phe represents a shift from aliphatic to aromatic amino acids (Table 3, Table S4). MD simulations for AT1G14310 showed that proteins of the haplotype H6 (Col-0; Leu20) exhibited lower backbone deviations compared to H2 (Gr-1; Phe20), which stabilized at a slightly higher RMSD. RMSF analysis revealed comparable flexibility profiles, with the most pronounced fluctuations occurring at the N-terminal region (Fig. S1). Hydrogen bond analysis demonstrated that Leu20 formed multiple persistent interactions with neighboring residues, whereas Phe20 did not display stable hydrogen bonds (Fig. S1). These findings suggest that the Leu20Phe substitution weakens local interactions, causing small differences in stability between the haplotypes.

AT5G35695 shows multiple substitutions affecting polarity and charge, such as Gly80Arg (nonpolar to positive) and Asp184Lys or Asp184Glu (negative to positive/longer chain), which were associated with the root length under drought in the LAS (Table 3). These variations led to predicted changes in protein conformation (Fig. S1), which may affect their function. In AT5G35695, MD simulations showed that proteins of the H10 haplotype (HKT2-4) with Gly80 and Glu184 stabilized at a lower RMSD compared to the H21 haplotype (Ll-0) with Arg80 and Lys184, indicating reduced global deviations. RMSF analysis revealed consistently lower fluctuations for H10, with the most pronounced differences observed at the N-terminus and at the substitution sites. These results suggest that the Arg80Gly and Lys184Glu substitutions stabilize local structure and conformational dynamics of the AT5G35695 protein, which can explain their significant association with the root length phenotypes under drought in the LAS (Table 3).

Despite having multiple substitutions associated with tolerance, the AlphaFold predicted model of AT4G26990 revealed much disorder in the apoprotein without defined secondary structures. This protein may stabilize after its interaction with RNAs or other RNA-binding proteins during stress granule assembly.

### 3.5. T-DNA insertion mutants of candidate genes show altered drought tolerance

We screened the available T-DNA insertion mutants of genes associated with the topmost significant SNPs delineated by the GWAS (MLM *P* <10^−3^; Table 1) with the LP and RP primers (Table S3). Altogether, we obtained homozygous T-DNA insertion lines for 16 genes. These lines were tested for seedling-level drought tolerance under the same phenotyping conditions as the GWAS. The knockout lines of *PCMP-A4* (SALK_129884C), *ATRBP45C* (SALK_088676C), *INCENP* (SALK_071684C), AT4G26990 (SALK_200826C), and *SKS6* (SALK_036979C), having the T-DNA inserted at exons (Fig. S2), showed a significant (Tukey’s test, *P*-value < 0.05) 20.1 ± 3.7%, 30 ± 7.1%, 38.6 ± 3%, 53.2 ± 7.5%, and 35.7 ± 1.1% reduction in RRL, respectively, compared to the WT (Fig. 7). The mutant of *ANAC094* (WiscDsLoxHs069_08D), with the T-DNA localized at an intron, showed a 38.9 ± 3.7% reduction in RRL. Knockdown lines of *ACD55.5* with T-DNA insertions in the promoter (SAIL_583_D08) exhibited a drastic 57.3 ± 2.1% and 36.6 ± 1.6% reduction in RRL corresponding to a 0.29‒0.47-fold gene expression compared to the WT (Figs. S2, S3). Knockdown lines of AT1G06690 (SALK_127525C; 5’ UTR), with 0.4‒0.6-fold gene expression compared to the WT, displayed a drastic 62.8 ± 3.8% and 60.7 ± 2.3% reduction in RRL. The mutants of *MSH6* (SALK_089638C), *GRF7* (SALK_084141C), and AT1G14315 (SAIL_837_C08) had T-DNA inserted at the exons. Among them, *GRF7* (SALK_084141C; exon) and AT1G14315 (SAIL_837_C08; exon) showed 45.9 ± 4.2% and 36.7 ± 5% increase in RRL compared to WT, indicating their negative role in drought tolerance. The mutants of AT1G14310 (SALK_129900C; 5’-UTR) and *NRPD4* (SALK_020157C; promoter) showed no significant reduction in gene expression compared to the WT (Figs. S2, S3). *NSE4A* (SALK_070555C; promoter) knockdown lines, with 0.4‒0.6-fold gene expression compared to the WT, displayed drought tolerance similar to the WT (Fig. 7, Fig. S2). Independent knockdown lines, *tad3*(1) and *tad3*(2), having the T-DNA inserted at the 5’-UTR (SALK_009025C), exhibited 25.9 ± 8% and 21.9 ± 6% reduction in RRL under drought, corresponding to a 0.6‒0.8-fold reduction in *TAD3* expression levels compared to the WT. Knockdown lines of *BAG1* with promoter insertion (SALK_017591C) showed 23.7 ± 8.4% and 27.3 ± 10.3% decrease in RRL, corresponding to 0.1‒0.4-fold gene expression compared to the WT (Fig. 7; Figs. S2, S3). However, the RRL reduction of *tad3* and *bag1* knockdown lines did not pass the significance criteria (Tukey’s test, *P* < 0.05). The reverse genetic analysis with T-DNA insertion mutants indicated a direct association of the genes AT1G06690, *ACD55.5*, *INCENP*, *SKS6*, *ANAC094*, *ATRBP45C*, AT4G26990, and *PCMP-A4* with seedling drought tolerance in *A. thaliana*.

**Fig. 7.**
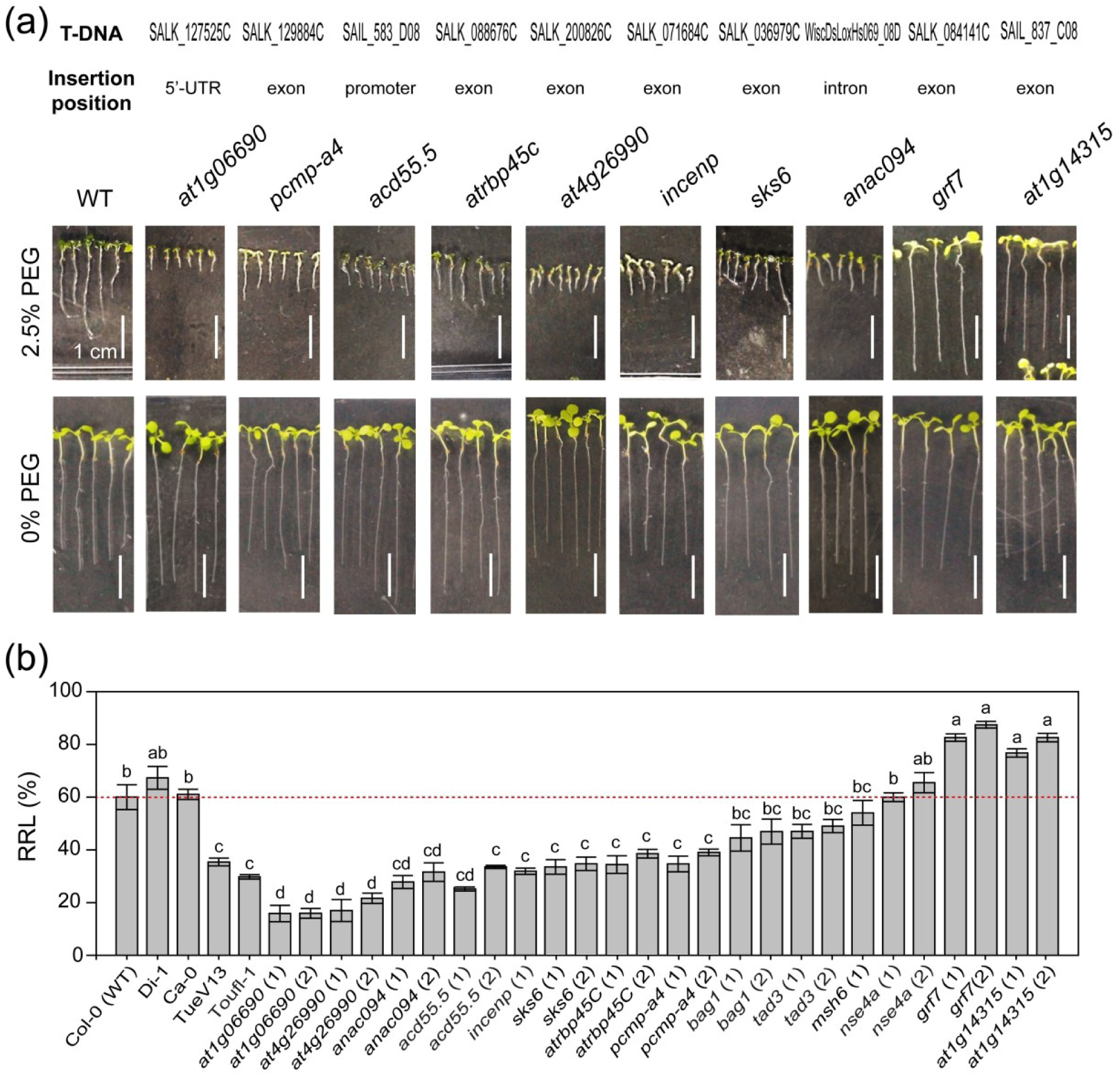
Drought tolerance of T-DNA insertion mutants of GWAS-delineated genes. **(a)** Hydroponically grown plants were placed on solidified agar plates and photographed. Representative images of T-DNA insertion mutants and their wild-type (WT) background (Col-0; NS60000) grown under drought (¼-Hoagland’s solution with 2.5% PEG) and non-stressful conditions (0% PEG) (see Methods) are shown. (**b**) Data for the wild-type and two homozygous T-DNA insertion lines for each gene, if isolated (see Supporting Information Fig. S2), are shown as average values of the relative root length (RRL), i.e., root length in 2.5% PEG/0% PEG, with the standard error of the longest eight of 20 seedlings. Two tolerant (Di-1 and Ca-0) and two sensitive (TueV13 and Toufl-1) ecotypes used in the GWAS (see Fig. 1; Supporting Information Table S1) were used as positive and negative controls. Asterisks indicate significant differences in RRL between the genotypes (Tukey’s test, *P* < 0.05). Vertical white bars indicate 1 cm. The red dotted line indicates the average RRL of WT. The experiment was repeated three times with similar results.

To further validate the phenotypes obtained in hydroponics, we cultured the homozygous T-DNA insertion mutants employed in the study aseptically in solidified sucrose-containing growth media under high salt (NaCl) (Fig. S4). Under 100 mM NaCl, mutants of AT4G26990, *INCENP*, *SKS6*, *PCMP-A4*, *ACD55.5* and *ANAC094* showed a significant (Tukey’s test, *P* < 0.05) reduction in RRL by 22.5 ± 2.5%, 28.4 ± 4.1%, 28 ± 2.3%, 43.9 ± 2.2%, 35.6 ± 7.3% and 43.4 ± 0.7%, respectively. On the other hand, mutants of AT1G14315 and *GRF7* showed a significant (Tukey’s test, *P* < 0.05) increase in RRL by 26.5 ± 4.7% and 45.6 ± 5.1%, respectively, from the WT. In the presence of a higher dose of 150 mM NaCl, the knock-out lines of *PCMP-A4* and knock-down lines of *ACD55.5* and *INCENP* maintained a significant (Tukey’s test, *P* < 0.05) reduction in RRL by 37.4 ± 4.2%, 38.7 ± 3.6% and 35.2 ± 4.5%, respectively, compared to the WT. It was observed that most of the mutants showing sensitivity to PEG in hydroponics also displayed sensitivity to NaCl in solid media, indicating similar osmotic stress conditions imposed by both the stressors. On the other hand, mutants of AT1G14315 and *GRF7*, two negative regulators of drought tolerance, also demonstrated insensitivity to salt stress, further strengthening their roles in drought sensitivity.

To identify the potential involvement of the GWAS-delineated genes for drought resistance in ABA signaling, we cultured their mutants in ABA-containing solid media along with the WT. Under 2.5 µM ABA, mutants of AT4G26990, AT1G06690, *ANAC094*, and *PCMP-A4* showed significant (Tukey’s test *P* < 0.05) reduction in RRL by 71.1 ± 2%, 70.1 ± 8.2%, 44 ± 9.6%, and 30.2 ± 2.6%, respectively, compared to the WT. In the presence of a higher concentration of 5 µM ABA, mutants of *ACD55.5*, *ATRBP45C*, AT4G26990, and *PCMP-A4* showed a significant (Tukey’s test *P* < 0.05) reduction in RRL by 70.1 ± 1.8%, 58.6 ± 5.1%, 54.1 ± 4.2%, and 35.8 ± 13.5%, respectively, compared to the WT. Altered ABA sensitivity of the mutants indicates their roles in the perception, transduction, or downstream response of ABA signaling, needing further experimental validation.

### 3.6. Expression levels of AT1G06690 contribute to differential ROS accumulation in roots

The reported involvement of the paralogs of AT1G06690 in the production of the cellular antioxidant PLP (Herrero et al., 2011), along with our observed drastically sensitive root length phenotype of the knockdown line of AT1G06690 (Fig. 7) prompted us to quantify the H_2_O_2_ levels in the seedling root tissues of 24 randomly chosen ecotypes, among those used in the ELP analysis of AT1G06690. A significant difference (Student’s *t*-test; *P* < 0.05) in average H_2_O_2_ levels was observed between ecotypes with contrasting SNP alleles at *Chr1:2052275* on the promoter of AT1G06690, having a significant association with the drought tolerance phenotype in the LAS (Fig. 4; Table 3). Ecotypes with the tolerant allele, C, showed an average H_2_O_2_ concentration of 40.5 µmoles g^−1^ FW, while those with the sensitive allele, G, showed a higher average of 69.6 µmoles g^−1^ FW. We also observed a significant difference in the root H_2_O_2_ levels between the WT (136.8 µmoles g^−1^ FW) and the knockdown line of AT1G06690, SALK_127525C (170.9 µmoles g^−1^ FW). In both comparisons, a higher expression of AT1G06690 coincided with lower H_2_O_2_ levels, and vice versa (Fig. 8). The same trend was observed in an independent repeat of the entire experiment. These results indicated the contribution of AT1G06690 to differential ROS accumulation in roots, explaining the natural variation in root growth under drought.

**Fig. 8.**
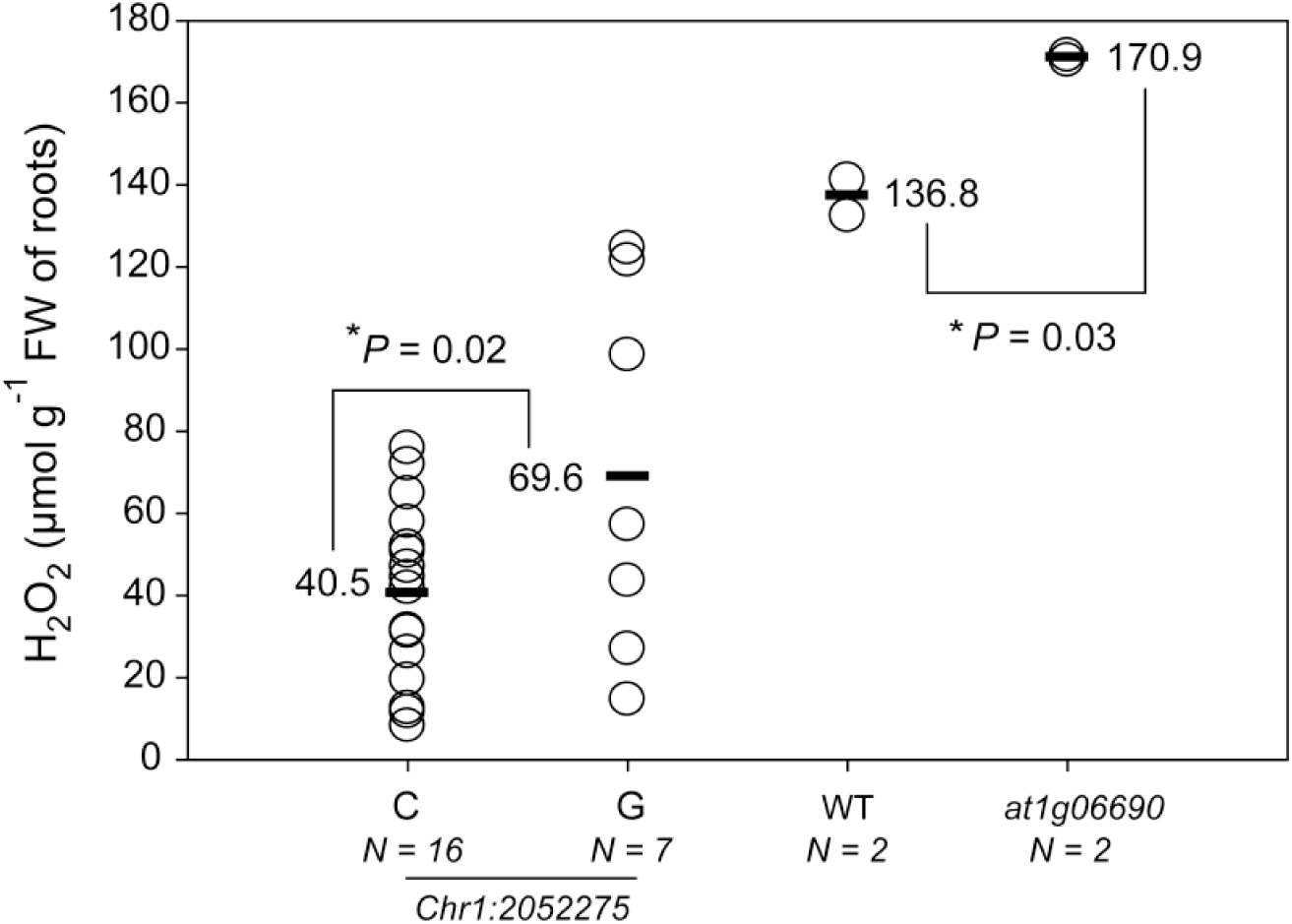
Root hydrogen peroxide content affected by the AT1G06690 locus. *Arabidopsis thaliana* plants of different ecotypes were grown hydroponically in control ¼-Hoagland’s media for 10 days and transferred to media with 2.5% PEG for five days. Twenty-four ecotypes containing the single-nucleotide polymorphism (SNP) alleles C or G at *Chr1:2052275* on the promoter of AT1G06690, the top-ranking gene in the GWAS (see Fig. 4), along with two independent lines of the T-DNA insertion mutant *at1g06690* (SALK_127525C) and its wild-type (WT) background (Col-0; NS60000), were used in the experiment. H_2_O_2_ levels were measured spectrophotometrically (see Methods) and expressed as µmoles g^−1^ fresh weight (FW) of roots. The average from three technical replicates is presented for each data point. Asterisks indicate significant differences between pairwise comparisons as shown in the figure (Student’s *t*-test, *P* < 0.05, *N* = number of biological replicates).

## 4. Discussion

The current GWA study unraveled genome-wide genetic loci contributing to drought tolerance by exploiting the natural variation in seedling root growth of *A. thaliana* under PEG-induced drought stress in the laboratory. Our hydroponic phenotyping platform used a low ionic strength growth medium, viz. ¼ Hoagland’s solution amplifying the stress effects of PEG at a very low concentration, viz. 2.5% w/v (Marik et al., 2024), in *A. thaliana* seedlings, capturing the wide variation in RRL among ecotypes at the same time (Fig. 1). Previous studies indicated that drought tolerance alleles of *A. thaliana* originated in the Mediterranean basin and Scandinavian coasts with very low rainfall (Exposito-Alonso et al., 2018; Exposito-Alonso et al., 2019). In our study, the ecotype Kondara from an area with a very low annual rainfall of < 500 mm showed high RRL under drought. A previous GWAS also identified longer root cells, including those at the meristem and elongation zone, in Kondara (Meijón et al., 2014). Eil-0 and Lp2-2, from moderately low rainfall (500-600 mm/year) areas, exhibited tolerance under our conditions, corroborating a high ABA accumulation in these ecotypes under drought (Kalladan et al., 2017). Wl-0, moderately tolerant under hydroponic conditions in our study (RRL 52.1%), demonstrated deeper roots in soil in another GWAS (Deja-Muylle et al., 2022). Ecotypes with moderate to high tolerance in our study, Ws-2 (40.85%), Est-1 (48.28%), NFA-10 (53.35%), Bor-4 (74.59%), and Lz-0 (71.96%), exhibited larger leaves under drought conditions (Clauw et al., 2016). The drought-sensitive ecotypes in our study primarily came from heavy rainfall areas (Fig. 1c, Table S1). Sensitive ecotypes from our study, Se-0 (26%), Uk-1 (17.4%), and Ak-1 (11.95%), also exhibited sensitive root cellular characteristics in a previous report (Meijón et al., 2014). RRS-7, showing root growth sensitivity in our study, exhibited non-functional stomata when exposed to varying concentrations of ABA under moderate to low vapor pressure deficits (Aliniaeifard et al., 2014). Despite coming from an area with only 352 mm of annual rainfall, Kz-9 (24.3%), one of the sensitive ecotypes in our study, accumulated medium amounts of ABA (Kalladan et al., 2017). On the other hand, ecotypes exhibiting low ABA accumulation, such as Ob-1, Sha, and Sorbo, demonstrated medium root lengths of 45.21%, 47.15%, and 48%, respectively, in our GWAS. Thus, a comparative analysis of different phenotypes across GWA studies suggested that our hydroponic phenotyping platform had effectively captured the natural variation of drought stress tolerance in *A. thaliana*, also indicating a connection between shoot and root traits for drought tolerance.

The top 10 most-significant SNPs detected by our GWAS could explain 47% of the observed variation in root length (Fig. 2b). The ridge regression curve reached a saturation, with 200 SNPs explaining 80% of the variance. The high contribution of the top 10 SNPs was also apparent from suggestive deviations from the expected *P*-values in the Q-Q plot (Fig. 2a). The genes associated with the top-ranking SNPs (*P* < 10^−3^) were mostly associated with stress tolerance, as indicated by the functional enrichment analysis (Table 2). Public gene expression data also indicated responsiveness of many of these genes to drought, osmotic stress, high salt, as well as the drought hormone, ABA, indicating that the GWAS had unraveled loci for drought tolerance (Table 1). We explored the ELPs and protein-level variation of the genes linked to the 10 most-significant SNPs to gather genetic evidence and identify the causative polymorphisms for drought tolerance. Concurrently, we conducted a network analysis of the genes associated with the top-ranking SNPs, revealing distinct processes of cell cycle regulation, DNA repair, tRNA modification, organellar translation, protein folding, and quality control responsible for root growth maintenance of *A. thaliana* under drought. Notably, some of the genes identified in our GWAS or their orthologs reportedly function in salt tolerance. Salt stress is a combination of an early osmotic and late ionic stress component, while drought presents solely an osmotic stress condition (Munns et al., 2008). The drought tolerance phenotypes of most of the knockout/knockdown mutants could be replicated under high salt stress in MS media (Fig. S4). ABA sensitivity of a mutant often reflects the involvement of the corresponding gene in ABA signaling (Ghassemian et al. 2000). Furthermore, desiccation tolerance is lost and re-established within a short developmental window in germinating seeds, relying on ABA signaling (Maia et al. 2014). Genetic studies show that loss of ABA sensitivity can pinpoint key regulators of the ABA signaling pathway, implying direct gene involvement in signal transduction. Conversely, ABA-sensitive mutants may reveal genes acting downstream or in parallel within the pathway, sometimes at the transcriptional or translational regulation levels (Fujii & Zhu 2009). In our study, mutants of AT1G06690, *PCMP-A4*, *ACD55.5*, *ATRBP45C*, AT4G26990, and *ANAC094* exhibited ABA hypersensitivity, indicating them to be putative negative regulators of the ABA signaling pathway (Yoshida et al. 2006; Umezawa et al. 2010). This notion is supported by the reported role of *CrNAC036*, an ortholog of *ANAC094*, in suppression of ABA biosynthesis by down-regulation of *9-cis-epoxy carotenoid dioxygenase 5* (*NCED5*) in *Citrus reticulata* (Zhu et al. 2020). However, additional research is required on the involvement of the GWAS-delineated genes in ABA signaling.

Detection of genes related to oxidative stress in the current drought tolerance GWAS points to the differential abilities of ROS generation/ scavenging between ecotypes in root tissues, as evidenced by our biochemical tests (Fig. 8), underlying their differential drought tolerance. The most significant SNP of the GWAS was on an exon of AT1G06690, a gene reportedly involved in osmotic stress resistance through the regulation of the PLP salvage pathway (Yu et al., 2020). In plants, the PLP salvage pathway, converting inactive forms of vitamin B_6_, pyridoxamine (PM), pyridoxine (PN), and pyridoxal (PL), into active PLP through the action of kinases, oxidases, and reductases, is a key mechanism for protection against drought and other abiotic stressors (Mooney et al., 2009). Mutations in pyridoxine/pyridoxamine phosphate oxidase (PPOX), involved in the conversion of pyridoxine 5′-phosphate (PNP) and/or pyridoxamine 5’-phosphate (PMP) into PLP, showed reduced root growth in Arabidopsis (Sang et al., 2011). Mutants of *pyridoxal 5’-phosphate synthase subunit 1* (*PDX1*), encoding part of an enzyme for *de novo* biosynthesis of PLP, also displayed markedly impaired root growth in Arabidopsis (Wagner et al., 2006). *Salt-overly sensitive 4* (*SOS4*), encoding a PL kinase, significantly contributed to salt tolerance, with its mutant showing salt sensitivity (Wang et al., 2004). The mutants of a paralog of AT1G06690, viz., *pyridoxal reductase 1* (*PLR1*), showed reduced root growth under drought (osmotic) and salt stresses, concomitant with higher levels of PN but lower levels of PM, PL, and the phosphorylated vitamers, pyridoxamine phosphate (PMP), pyridoxine phosphate (PNP), and pyridoxal phosphate (PLP), than the WT (Herrero et al., 2011). On the other hand, a higher drought tolerance of *pdx1.3* and *sos4* mutants compared to the WT plants was observed under soil culture conditions, attributable not to the PLP content, but to their smaller and more compact rosette morphology, which likely reduced their overall water loss (González et al., 2007). PLP functions as a biological antioxidant, and tobacco plants exhibited elevated levels of PLP under drought (Huang et al., 2013). *PDX1.1* concentrations increased in the root stele of ammonium-stressed Arabidopsis, as the production of non-phosphorylated vitamin B6 protected roots from oxidative stress (Liu et al., 2022). Corroborating these studies, our results suggest that the PLP salvage pathway is crucial for safeguarding roots against drought-induced excessive ROS, which justifies its association with the most significant SNP in our root-length GWAS. The ELPs in AT1G06690 caused by promoter SNPs, leading to alteration of *cis*-elements, were associated with drought tolerance (Fig. 4, Table 3). The NASC-procured knockout mutant seeds of AT1G06690, with the T-DNA inserted in an exon (SALK_058109C), did not germinate. Another T-DNA insertion event at the 5’-UTR (SALK_127525C), leading to gene knockdown, exhibited drought sensitivity in our study (Fig. 7; Fig. S2), as well as higher ROS levels in roots than the WT (Fig. 8). Other antioxidant genes identified by the GWAS included *LSU2* and *LSU4*, first reported in the context of sulfur deficiency, which participate in hormone signaling and stress responses in plants (Canales et al., 2023). LSUs are responsive to phytohormones like ABA, auxin, and ethylene and regulate growth under stress. The *lsu* knockout lines are reportedly sensitive to salt stress due to altered sulfur metabolism, production of sulfur-containing amino acids, and the function of sulfur-dependent antioxidant glutathione (Andrés-Barrao et al., 2021). Iron-dependent superoxide dismutase 2 physically interacts with LSUs to enable the ROS (H_2_O_2_) production in chloroplasts. Our transcriptome data indicated a suppression of *LSU2*, possibly to limit ROS-induced damage under drought (Table 1). Another GWAS-identified candidate, *SBH2,* encodes a hydrolase predominantly found in roots, which prevents excessive buildup of sphingoid long-chain bases (LCBs), assisting in the regulation of ROS, sphingolipid signaling under stress, and maintaining membrane integrity (Nakagawa et al., 2011). Sphingoid base hydroxylation by enzymes such as SBH2 is important for the regulation of LCBs in sphingolipids (Chen et al., 2008). Sphingolipids, structural components of the cell membrane and endomembranes, also participate in signaling, playing a crucial role in ABA-mediated plant stomatal responses to drought (Ali et al., 2018). *SKS6*, a multicopper oxidase-like protein involved in oxidative stress response and growth processes, including cotyledon vascular patterning, cell expansion, directional cell growth, and cell wall remodeling (Sproule et al., 2008; Pavoković et al., 2023). Previous studies showed that *SKS6* is upregulated under salt stress (Pavoković et al., 2023) but downregulated under drought stress (Brilhaus et al., 2015). *SKS6* is expressed in guard cells, root cortex, and apical meristem and is induced in the root tips and epidermis in response to exogenous ABA (Jacobs & Szymanska, 2005). The *SKS6* knock-out exhibited reduced root length (Fig. 7), indicating the vital role of this gene in drought tolerance. Another upstream regulator of *SKS6*, an *ANAC094* TF (as indicated by public DAPseq data), was also detected by our GWAS (Table 1). This gene was induced under drought in our transcriptome analysis, and its knockout mutant exhibited drought sensitivity (Fig. 7). These results highlight the essential role of the *ANAC094*-*SKS6* module in root growth maintenance under drought, which is in line with previous reports indicating that ANAC094 regulates root development (Yi et al., 2010; Nie et al., 2011).

Two RNA-binding proteins, *RBP45C* and AT4G26990, associated with the top-ranking SNPs delineated by the GWAS (Tables 1, 2), were involved in the enriched biological process of stress granule assembly. Stress granules are non-membranous condensates in the cytoplasm composed of proteins and RNA. The translation of non-essential proteins is temporarily suppressed under stress, and untranslated mRNAs are protected from degradation through incorporation into stress granules (Marondedze et al., 2020). AT4G26990 was upregulated by drought under our experimental conditions (Table 1), and polymorphisms in this gene translated into a large number of amino acid substitutions having a significant association with drought tolerance (Table 3). These may influence the interaction of AT4G26990 with target mRNAs or other proteins for stress granule assembly. *RBP45C* is typically induced under hypoxia (Schippers et al., 2024). The knock-out mutant of *RBP45C* and AT4G26990 displayed significant drought sensitivity (Fig. 7). These results suggest that differential abilities of ecotypes in the protection of essential RNAs through stress granule assembly play a major role in the natural variation of drought tolerance in *A. thaliana*.

*PCMP-A4*, delineated by our GWAS, encodes a protein of the PPR superfamily involved in post-translational RNA processing within organelles and plays vital roles in drought tolerance (Su et al., 2019; Wang et al., 2021; Luo et al., 2022; Wang & Tan, 2025). *Brassica rapa* orthologs of this gene were predicted to have evolved under drought through sequence analysis of ancestral and descendant natural populations (Franks et al., 2016). Loss of function of PPR proteins leads to dysfunctional mitochondria and chloroplasts, leading to growth defects (Wang & Tan, 2025). The knockout lines of *PCMP-A4* showed drought sensitivity in our study (Fig. 7). In addition, genetic evidence included sensitive haplotypes leading to truncated PCMP-A4 proteins, causing possible loss of function (Fig. 5; Table S4). *TAD3*, encoding an essential tRNA base editing enzyme, was detected by our GWAS with five significant SNPs (<10^−3^) directly localized on its exons and introns (Table 1). Two of these SNPs, *Chr5:8449400* and *Chr5:8450773*, led to Gln85His and Leu336Trp substitutions, respectively, in the TAD3 protein. The TAD3 enzyme deaminates adenine to inosine in the anticodon loop of cytoplasmic tRNAs to facilitate base pairing with codons at the wobble position. Previous reports indicate that TAD3 plays a role in ABA signaling, seed dormancy, and germination (Vu et al., 2015). TAD3 is also crucial for telomere length maintenance in *A. thaliana*, preventing chromosome ends from DNA damage (Bose et al., 2020). Knockout mutants of *TAD3* were seedling lethal, being arrested at the embryo stage, whereas knockdown lines showed growth retardation (Zhou et al., 2014). Moreover, *TAD3* exhibited epialleles and was found methylated in Nok-1, in contrast to Col-0 (Agorio et al., 2017). Under our experimental conditions, knockdown lines of *TAD3* exhibited a slight ∼20-25% reduction (*P* = 0.07) in RRL under drought (Fig. 7). Different RNA modifications, including tRNA base editing, have been implicated in abiotic stress tolerance (Cai et al., 2025). Specifically, tRNA editing at the wobble positions increases under salt stress (Janssen et al., 2022). TAD3 was also involved in an interaction network with other proteins involved in nucleobase modifications (Fig. 3c). These included *GSDA*, encoding a guanosine deaminase, whose drought-responsive rice ortholog *OsGSDA* was found to regulate gene expression through epigenetic modifications (Gotarkar et al., 2021). Another gene in the network, *PYRD*, encoding a pyrimidine deaminase, is reportedly induced in *DREB2G*-overexpressing and suppressed in *DREB2G* knockout plants. Furthermore, a dehydration-responsive element (DRE) was identified in the promoter region of the *PYRD* gene, suggesting its involvement in drought response (Namba et al., 2024). These observations support the roles of *TAD3* and its networking genes in drought tolerance.

*EMB2788*, involved in ribosome biogenesis by catalyzing the maturation of 5.8S rRNA from the tricistronic precursor rRNA transcript, comprising 18S rRNA, 5.8S rRNA, and 28S rRNA, was identified by our GWAS. *EMB2788* was also induced under drought, as identified in our transcriptome analysis and also found in a previously reported ABA-response proteome analysis (Zhang et al., 2019). Ribosome diversity is enhanced during drought stress, as plants need to adjust their protein production machinery by decreasing overall protein synthesis and encouraging the translation of stress-responsive mRNAs. This phenomenon has been noted in rice along with varied expression of ribosome-binding proteins in tomato (Martinez et al., 2020). In network ‘e’, *EMB2788* interacted with other genes associated with ribosome biogenesis, which were notably affected during drought. In drought-sensitive ecotypes of chickpea, ribosomal genes were identified as downregulated and hypomethylated, resulting in decreased ribosome biogenesis (Yadav et al., 2024). Heat shock proteins (HSPs) function as chaperones, preventing protein misfolding under different stresses, providing resistance to plants. In our research, certain HSPs were identified as potential candidates for drought tolerance by the GWAS, specifically *ATHSBP*, *HSP20-like chaperones superfamily protein* (ACD55.5; AT5G02480), and AT5G02502 (Jungkunz et al., 2011; Guan et al., 2013; Landi et al., 2017). *ATHSBP/HSP21* is a heat shock protein that inhibits the aggregation of denatured proteins under stress. Expression of a miRNA gene, *MIR156*, under the *HSP21* promoter resulted in enhanced growth and was associated with heat stress memory (Stief et al., 2014). *BAG1*, identified by our GWAS, localizes to the cytoplasm and interacts with a chaperone, heat shock cognate protein 70 (Hsc70), for proteasomal degradation of unimported plastid proteins (Lee et al., 2016), playing a role in cell protection during stress and inhibiting apoptosis (Doukhanina et al., 2006). In line with the genes regulating protein folding, our GWAS detected *CCT6-2*, encoding a subunit of the CCT chaperonin complex involved in folding of important cytoskeletal proteins actin and tubulin, which may have implications in cell growth and division leading to differential root growth regulation under drought (Table 1; Fig. 3). The network analysis indicated that CCT6-2 interacted with TAP46, acting as a positive regulator of TOR and ABA signaling (Fig. 3g). TAP46 is a protein phosphatase2A (PP2A)-associated protein activated by TOR kinase. TAP46 interacts with the ABA signaling TF, ABA insensitive 5 (ABI5), stabilizing it and preventing its negative regulation by PP2A or other phosphatases (Punzo et al., 2018; Hu et al., 2013). TOR signaling plays a role in drought response, as the ectopic expression of TOR in rice improved water usage efficiency and yield. Moreover, TOR overexpressors exhibited extended primary root growth when subjected to potassium chloride stress (Fu et al., 2020). A GWAS-identified gene, *raf43*, encoding a MAPKKK, lies upstream in drought signaling. *Raf43* was reportedly upregulated under drought (Hahn et al., 2013). The *raf43-1* T-DNA insertion mutants exhibited reduced expression of *RD17* and *DREB2A* genes, highlighting their involvement in drought tolerance (Virk et al., 2015).

Another aspect of drought stress signaling delineated by our GWAS is ER–PM tethering, represented by the candidate gene *VAP27-1*. Protein misfolding under stress induces ER stress, which is mitigated by an unfolded protein response (UPR). The ER and PM develop contact sites through tethering for the exchange of materials. This ER-PM connectivity has been recently demonstrated to play an active role in UPR, as the *vap27-1/3/4* triple mutant presented defects in ER-TM tethering and UPR signaling (Man et al., 2024). In the backdrop of these findings, identification of the key gene involved in ER-PM tethering, *VAP27-1*, by our GWAS indicates the importance of ER stress and UPR signaling for seedling drought tolerance in *A. thaliana*.

The GWAS-identified gene *APC5* interacted with other *APC* paralogs in the network ‘f’ (Fig. 3) to constitute the APC/C, a ubiquitin ligase complex marking specific cell cycle-regulating proteins (cyclins) for ubiquitin-mediated degradation to enable mitotic exit, keeping a check on endoreplication (Carneiro et al., 2021). It has been observed that the mitotic exit of proliferating cells triggered by stress is influenced by the modulation of APC/C (Claeys et al., 2012). APC/C acts as a positive regulator of GA signaling and is inhibited by drought and ABA-induced SNF1-related protein kinase 2 (SnRK2), highlighting its role in growth modulation under drought (Yang et al., 2019). Drought-induced lowering of gibberellic acid (GA) in rice inhibits plant growth, enhancing the levels of the negative GA signaling regulator SLENDER RICE 1, a rice DELLA protein inhibiting the breakdown of the ABA receptor pyrabactin resistance 1-like (PYL) through competitive binding with the APC/C activator, tillering, and dwarf 1. This led to enhanced ABA signaling and drought tolerance (Liao et al., 2023). Hence, the GWAS identification of *APC5*, encoding a subunit of the APC/C, points to the role of differential regulation of the cell cycle in maintaining Arabidopsis root growth under drought. Another GWAS-identified candidate, *INCENP*, encodes a centromere protein that facilitates the attachment of kinetochores to the microtubules (Bray et al., 2024). In our research, this gene was identified as a positive regulator of drought tolerance, with its knock-out showing drought sensitivity (Fig. 7).

DNA repair was among the enriched GO biological processes and KEGG pathways encompassing six GWAS-detected genes (Table 2, Fig. 3). This process also emerged as one of the major networks (Fig. 3). DNA damage due to excessive ROS is a hallmark of drought stress (Nisa et al., 2019). Hence, genes involved in DNA replication and their associated proteins, like helicases and topoisomerases involved in damage repair, are paramount in root growth maintenance under drought. Out of the GWAS-identified genes, AT1G14460 encodes a DNA polymerase involved in DNA repair (Liu et al., 2011). *RAD23-3* is a component of nucleotide excision repair and damage tolerance complex (Kunz et al., 2005; Mahapatra et al., 2020; Ena et al., 2015). *NSE4A* is a nucleus-localized protein engaged in DNA repair in a complex with *SMC5* and *NSE3* (Díaz et al., 2019). The roots of the *nse4a-2* mutant revealed an increased number of dead cells in the meristematic zone, thereby reducing the root length (Diaz et al., 2019). In our study, T-DNA insertion events in the promoter of *NSE4A*, leading to a 0.4‒0.6-fold decrease of gene expression (Fig. S1), did not show a significant reduction in RRL (Fig. 7). *MSH6* corrects mismatches in DNA and is upregulated in response to UV-B-induced DNA damage (Lario et al., 2011). The *MSH6* promoter region is differentially methylated in the progenies of salt-stressed plants, indicating roles in salt/osmotic stress tolerance and memory (Bilichak et al., 2012). A single homozygous knockout line of *MSH6* isolated by us also had no significant difference in drought tolerance from the wild type, possibly due to genetic redundancy. Rice orthologs of *DNA topoisomerase 4 subunit B* (AT4G11670), detected by our GWAS, were highly induced under salt and drought and conferred drought tolerance when heterologously expressed in *E. coli* (Li et al., 2018). *DUT1* encodes another DNA-repairing enzyme with dUTPase activity, which, upon silencing, led to increased DNA damage (Siaud et al., 2010). *MCM7*, a major hub gene identified by the network analysis, is a component of the DNA helicase machinery, reportedly downregulated under drought for preservation of resources, thereby reducing cell division (Brilhaus et al., 2015). However, we could not isolate homozygous T-DNA insertion lines of *MCM7* and *DUT1*.

AT5G35695, encoding a harbinger transposase-derived 1 (HARBI1)-like endonuclease, showed truncated proteins in drought-sensitive ecotypes (Table S4). This gene is a domesticated version of transposases, which play important roles in plant growth and cadmium stress responses by regulating transport processes, antioxidant defense, and phytohormone signaling, through interaction with TFs and epigenetically through histone modification (Huo et al., 2021; Jiang et al., 2023). Detection of AT5G35695 in the current GWAS, along with the protein polymorphisms associated with the phenotype, suggested its role in drought tolerance, which could not be verified due to the lack of a homozygous T-DNA insertion line for this gene in our reverse genetic screen. Future research is warranted to reveal the mechanism of action of AT5G35695 in drought signaling.

Our GWAS identified some of the negative regulators of drought tolerance, validated by reverse genetics, as far as homozygous T-DNA insertion lines could be isolated for these genes (Table 1, Fig. 7). The GWAS-identified candidate gene *GRF7* is a known negative regulator of drought tolerance, which represses ABA and drought-responsive genes, e.g., *DREB2A*. *GRF7*-deficient plants had higher survival rates under salt and drought stresses (Kim et al., 2012). In line with the previous reports, the knockout line of *GRF7* showed greater drought tolerance than the wild type in our study (Fig. 7). AT1G14310, encoding a haloacid dehalogenase-like hydrolase family protein of unknown function, was reportedly downregulated under salt stress (Chaves et al., 2009). We also noticed a downregulation of this gene under drought in Col-0 by our transcriptome analysis (Table 1). Overexpression of *OsHAD3* in rice reduced drought tolerance, indicating its negative role in drought (Zan et al., 2023). AT1G14315, encoding an F-box domain-containing protein, was downregulated under drought stress in *Chrysopogon zizanioides* (George et al., 2017). In our study, the knock-out of this gene showed higher RRL than the WT, indicating it as a negative regulator of drought tolerance (Fig. 7). Interestingly, the ecotypes from low rainfall areas contained frameshift and stop codon gains (haplotypes H3 and H7), leading to truncated proteins. Although the F-box domain was preserved at the extreme N-terminal end in each of these polymorphic forms, protein truncations and frameshifts led to altered predicted folding, leading to possible implications in their functions (Fig. 5, Table S4). Out of the ecotypes with truncated protein, three (Kondara, Sorbo, and Stepn-1) showed moderate to high drought tolerance, while four (Yeg-1, Ms-0, Sij-1, and Sij-2) were found sensitive. This might be due to epistatic effects of other loci, as often observed in the case of climate-associated alleles (Aggarwal et al., 2010). Two Spanish ecotypes from moderate rainfall regions, Ll-0 and Bla-1, were also found to contain a haplotype (H10) with a frameshift leading to a truncated protein, which might contribute to their moderate to high drought tolerance in this study (Table S4). Other GWAS-identified drought-suppressed genes in Col-0 included *LBD41*, *TPPI*, *LSU2*, and *UGT78D2*. *LBD41*, a TF belonging to the lateral organ boundary domain (LBD) family, is a known drought-responsive gene in Arabidopsis (Bray et al., 2002). It is a key regulator of hypoxia-induced genes, and its activity is modulated by oxidative stress common in drought (Schippers et al., 2024). An ortholog of *LBD4*, maize-specific *LBD33*, is a negative regulator of drought tolerance (Xiong et al., 2025). Again, *LBD41* was found to be downregulated during the formation of storage roots (Xu et al., 2016). The downregulation of *LBD41* in our transcriptome indicates its possible negative role in root growth under drought in *A. thaliana*. The drought-suppressed gene *TPPI* encodes trehalose-6-phosphate phosphatase, which converts trehalose-6-P into trehalose, aiding in the osmotic adjustment and removal of ROS (Lin et al., 2023). The expression of rice *TPPI* in maize conferred drought resistance (Nuccio et al., 2015). Arabidopsis has 11 TPP members that exhibit a varied spatio-temporal distribution. *TPP* members are targeted by DREB1A to regulate stomatal movement and drought tolerance (Lin et al., 2019; Lin et al., 2020). *TPPI* also plays a role in carbohydrate distribution between plant organs, promoting growth (Chai et al., 2025). Our GWAS detection of *TPPI* indicates a differential carbohydrate distribution dynamics in *A. thaliana* ecotypes, possibly contributing to their differential growth under drought. The activation of trehalose-6-phosphate (T6P)-SnRK signaling pathway and suppression of class I trehalose-phosphate synthases are hallmarks of drought responses. A downregulation of *TPP* orthologs of rose and poplar was also observed under drought (Jia et al., 2021; Fox et al., 2023). Through SnRK1, TOR, and auxin signaling, T6P also controls root branching and lateral root (LR) development, where *TPPI* was found to be induced in the LR primordia (Morales-Herrera et al., 2023). A potential reason behind the suppression of *TPPI* and *LBD41* might be the inhibition of LR emergence during drought. *UGT78D2*, encoding a flavonoid 3-O-glucosyltransferase in Arabidopsis, is responsible for the glycosylation of flavonoids and anthocyanins, whose accumulation is crucial for cold acclimation through antioxidant defense (Schulz et al., 2016; Li et al., 2020), also regulating the polar transport of auxin (Yin et al., 2013). Although the GWAS detection of *UGT78D2* may indicate the role of flavonoid metabolism in the differential drought tolerance of *A. thaliana* ecotypes, the reason for the drought suppression of *UGT78D2* in the tolerant ecotype Col-0 (Table 1) needs future exploration.

## 5. Conclusions and scope for future research

The genome-wide association study (GWAS) effectively unraveled genetic loci contributing to drought tolerance by exploiting the natural variation in seedling root growth of *Arabidopsis thaliana* under PEG-induced drought stress in the laboratory. Through network and functional enrichment analyses, the study demarcated key biological processes such as DNA repair, tRNA editing, protein quality control, and cell cycle regulation, which function synergistically to maintain root growth under drought. The identified candidate genes were further validated through the identification of expression level polymorphisms, amino acid substitutions, and reverse genetic evaluation using T-DNA insertion mutants, confirming their direct association with seedling drought tolerance and revealing roles in critical mechanisms like stress granule assembly and the PLP salvage pathway. These findings provide a genome-wide understanding of molecular signaling networks responsible for root growth maintenance under drought. Experimental validation of the predicted polymorphic protein structures and genetic complementation analysis of transgenic plants introducing different tolerant and sensitive alleles identified in this study will help to reveal the exact causative variations contributing to differential drought tolerance among the *A. thaliana* ecotypes. On the other hand, drought tolerance phenotypes of the knockout and knockdown lines used in the current study remain to be tested in soil-grown mature plants. The insights gained into drought tolerance genes and molecular signaling networks in *A. thaliana* can be extended to other crop species, crucial for developing stress-resilient crops using contemporary breeding methods and emerging biotechnologies like gene editing.

## CRediT authorship contribution statement

DM: Investigation (all wet experiments), Visualization, Writing – Original Draft Preparation. SVT: Investigation (protein modeling and molecular dynamics simulations), Data Curation, Formal analysis, Visualization, Writing – Original Draft Preparation. RK: Investigation (climate association, network, ridge regression, and promoter *cis*-element analysis), Software, Visualization. SD: Software, Supervision, Methodology, Validation, Data Curation, Formal analysis, Resources (dry lab). AS: Conceptualization, Data Curation, Formal analysis, Visualization, Funding acquisition, Methodology, Resources (wet lab), Project Administration, Writing – Original Draft Preparation, Writing – Review & Editing.

## Acknowledgements

We are grateful to Prof. Hiroyuki Koyama and Dr. Yuriko Kobayashi, Gifu University, Japan, for gifting the photographic mounts used in phenotyping and providing genotype information.

## Data availability

All data are presented within the paper. The transcriptome raw data are submitted to the NCBI Sequence Read Archive with an accession number PRJNA1086208.

## References

Aggarwal S., Negi S., Jha P., Singh P.K., Stobdan T., Pasha M.A.Q., … Mukhopadhyay A. (2010) EGLN1 involvement in high-altitude adaptation revealed through genetic analysis of extreme constitution types defined in Ayurveda. Proceedings of the National Academy of Sciences 107, 18961–18966.

Agho C., Avni A., Bacu A., Bakery A., Balazadeh S., Baloch F. S., … & Fragkostefanakis S. (2024). Integrative approaches to enhance reproductive resilience of crops for climate-proof agriculture. Plant stress, 100704.

Agorio A., Durand S., Fiume E., Brousse C., Gy I., Simon M., … Bouché N. (2017) An Arabidopsis Natural Epiallele Maintained by a Feed-Forward Silencing Loop between Histone and DNA. PLOS Genetics 13, e1006551.

Agrahari R.K., Enomoto T., Ito H., Nakano Y., Yanase E., Watanabe T., … Kobayashi Y. (2021) Expression GWAS of PGIP1 Identifies STOP1-Dependent and STOP1-Independent Regulation of PGIP1 in Aluminum Stress Signaling in Arabidopsis. Frontiers in Plant Science 12.

Ahluwalia O., Singh P.C. & Bhatia R. (2021) A review on drought stress in plants: Implications, mitigation and the role of plant growth promoting rhizobacteria. Resources, Environment and Sustainability 5, 100032.

Ahn C.S., Han J.-A., Lee H.-S., Lee S. & Pai H.-S. (2011) The PP2A Regulatory Subunit Tap46, a Component of the TOR Signaling Pathway, Modulates Growth and Metabolism in Plants. The Plant Cell 23, 185–209.

Ahn H.-K., Yoon J.-T., Choi I., Kim S., Lee H.-S. & Pai H.-S. (2019) Functional characterization of chaperonin containing T-complex polypeptide-1 and its conserved and novel substrates in Arabidopsis. Journal of Experimental Botany 70, 2741–2757.

Ali U., Li H., Wang X. & Guo L. (2018) Emerging Roles of Sphingolipid Signaling in Plant Response to Biotic and Abiotic Stresses. Molecular Plant 11, 1328–1343

Aliniaeifard S. & van Meeteren U. (2014) Natural variation in stomatal response to closing stimuli among *Arabidopsis thaliana* accessions after exposure to low VPD as a tool to recognize the mechanism of disturbed stomatal functioning. Journal of Experimental Botany 65, 6529–6542.

Alonso-Blanco C., Andrade J., Becker C., Bemm F., Bergelson J., Borgwardt K.M., … Zhou X. (2016) 1,135 Genomes Reveal the Global Pattern of Polymorphism in *Arabidopsis thaliana*. Cell 166, 481–491.

Altschul, S. F., Gish, W., Miller, W., Myers, E. W., & Lipman, D. J. (1990). Basic local alignment search tool. Journal of Molecular Biology, 215(3), 403–410.

Andrés-Barrao C., Alzubaidy H., Jalal R., Mariappan K.G., de Zélicourt A., Bokhari A., … Hirt H. (2021) Coordinated bacterial and plant sulfur metabolism in *Enterobacter* sp. SA187–induced plant salt stress tolerance. Proceedings of the National Academy of Sciences 118.

Atwell S., Huang Y.S., Vilhjálmsson B.J., Willems G., Horton M., Li Y., … Nordborg M. (2010) Genome-wide association study of 107 phenotypes in *Arabidopsis thaliana* inbred lines. Nature 465, 627–631.

Berman H.M. (2000) The Protein Data Bank. Nucleic Acids Research 28, 235–242.

Bilichak A., Ilnystkyy Y., Hollunder J. & Kovalchuk I. (2012) The Progeny of *Arabidopsis thaliana* Plants Exposed to Salt Exhibit Changes in DNA Methylation, Histone Modifications and Gene Expression. PLoS ONE 7, e30515.

Blum, M., Andreeva, A., Florentino, L. C., Chuguransky, S. R., Grego, T., Hobbs, E., Pinto, B. L., Orr, A., Paysan-Lafosse, T., Ponamareva, I., Salazar, G. A., Bordin, N., Bork, P., Bridge, A., Colwell, L., Gough, J., Haft, D. H., Letunic, I., Llinares-López, F., … Bateman, A. (2025). InterPro: the protein sequence classification resource in 2025. Nucleic Acids Research, 53(D1), D444–D456.

Bose S., Suescún A.V., Song J., Castillo-González C., Aklilu B.B., Branham E., … Shippen D.E. (2020) tRNA ADENOSINE DEAMINASE 3 is required for telomere maintenance in *Arabidopsis thaliana*. Plant Cell Reports 39, 1669–1685.

Bradbury P.J., Zhang Z., Kroon D.E., Casstevens T.M., Ramdoss Y. & Buckler E.S. (2007) TASSEL: software for association mapping of complex traits in diverse samples. Bioinformatics 23, 2633–2635.

Bray E.A. (2002) Classification of Genes Differentially Expressed during Water-deficit Stress in *Arabidopsis thaliana*: an Analysis using Microarray and Differential Expression Data. Annals of Botany 89, 803–811.

Bray S.M., Hämälä T., Zhou M., Busoms S., Fischer S., Desjardins S.D., … Yant L. (2024) Kinetochore and ionomic adaptation to whole-genome duplication in Cochlearia shows evolutionary convergence in three autopolyploids. Cell Reports 44, 114576. 10.1016/j.celrep.2024.114576

Brilhaus D., Bräutigam A., Mettler-Altmann T., Winter K. & Weber A.P.M. (2015) Reversible Burst of Transcriptional Changes during Induction of Crassulacean Acid Metabolism in Talinum triangulare. Plant Physiology 170, 102–122.

Cai J., Shen L., Kang H. & Xu T. (2025) RNA modifications in plant adaptation to abiotic stresses. Plant Communications 6, 101229.

Canales J., Arenas-M A., Medina J. & Vidal E.A. (2023) A Revised View of the *LSU* Gene Family: New Functions in Plant Stress Responses and Phytohormone Signaling. International Journal of Molecular Sciences 24, 2819.

Cao J., Schneeberger K., Ossowski S., Günther T., Bender S., Fitz J., … Weigel D. (2011) Whole-genome sequencing of multiple *Arabidopsis thaliana* populations. Nature Genetics 43, 956–963.

Carneiro A.K., Montessoro P. da F., Fusaro A.F., Araújo B.G. & Hemerly A.S. (2021) Plant CDKs—Driving the Cell Cycle through Climate Change. Plants 10, 1804. 10.3390/plants10091804

Chai L., Wang H., Yu H., Li H., Yi D., Ikram S., … Li Q. (2025) Trehalose-6-Phosphate Phosphatase *SlTPP1* Adjusts Diurnal Carbohydrate Partitioning in Tomato. Plant, Cell & Environment. 10.1111/pce.15599

Chaves M.M., Flexas J. & Pinheiro C. (2008) Photosynthesis under drought and salt stress: regulation mechanisms from whole plant to cell. Annals of Botany 103, 551–560. 10.1093/aob/mcn125

Chen K., Gao J., Sun S., Zhang Z., Yu B., Li J., … Zhao Y. (2020) BONZAI Proteins Control Global Osmotic Stress Responses in Plants. Current Biology 30, 4815–4825.e4.

Chen Y., Chen L., Lun A.T.L., Baldoni P.L. & Smyth G.K. (2025) edgeR v4: powerful differential analysis of sequencing data with expanded functionality and improved support for small counts and larger datasets. Nucleic Acids Res. 53. 10.1093/nar/gkaf018

Chen M., Markham J.E., Dietrich C.R., Jaworski J.G. & Cahoon E.B. (2008) Sphingolipid Long-Chain Base Hydroxylation Is Important for Growth and Regulation of Sphingolipid Content and Composition in Arabidopsis . The Plant Cell 20, 1862–1878.

Claeys H., Skirycz A., Maleux K. & Inzé D. (2012) DELLA Signaling Mediates Stress-Induced Cell Differentiation in Arabidopsis Leaves through Modulation of Anaphase-Promoting Complex/Cyclosome Activity. Plant Physiology 159, 739–747.10.1104/pp.112.195032

Clauw P., Coppens F., Korte A., Herman D., Slabbinck B., Dhondt S., … Inzé D. (2016) Leaf Growth Response to Mild Drought: Natural Variation in Arabidopsis Sheds Light on Trait Architecture. The Plant Cell 28, 2417–2434.

Cooper M. & Messina C.D. (2022) Breeding crops for drought-affected environments and improved climate resilience. The Plant Cell 35, 162–186.

D’Andrea, L. (2003). TPR proteins: the versatile helix. Trends in Biochemical Sciences, 28(12), 655–662. 10.1016/j.tibs.2003.10.007

Deja-Muylle A., Opdenacker D., Parizot B., Motte H., Lobet G., Storme V., … Beeckman T. (2022) Genetic Variability of *Arabidopsis thaliana* Mature Root System Architecture and Genome-Wide Association Study. Frontiers in Plant Science 12.

Dias-Fields L. & Adamala K.P. (2022) Engineering Ribosomes to Alleviate Abiotic Stress in Plants: A Perspective. Plants 11, 2097.

Díaz M., Pečinková P., Nowicka A., Baroux C., Sakamoto T., Gandha P.Y., … Pecinka A. (2019) The SMC5/6 Complex Subunit NSE4A Is Involved in DNA Damage Repair and Seed Development. The Plant Cell 31, 1579–1597.

Dossa K., Zhou R., Li D., Liu A., Qin L., Mmadi M.A., … You J. (2021) A novel motif in the 5’-UTR of an orphan gene ‘Big Root Biomass’ modulates root biomass in sesame. Plant Biotechnology Journal 19, 1065–1079.

Doukhanina E.V., Chen S., van der Zalm E., Godzik A., Reed J. & Dickman M.B. (2006) Identification and Functional Characterization of the BAG Protein Family in *Arabidopsis thaliana*. Journal of Biological Chemistry 281, 18793–18801. 10.1074/jbc.M511794200.

Ena S., Sutkovic J.. & Ragab Abdel Gawwad M. (2015) Interactome Analysis and Docking Sites Prediction of Radiation Sensitive 23 (RAD 23) Proteins in *Arabidopsis thaliana*. Current Proteomics 12, 28–44. 10.2174/1570164612666150225234240.

Eugeni Piller L., Abraham M., Dormann P., Kessler F. & Besagni C. (2012) Plastid lipid droplets at the crossroads of prenylquinone metabolism. Journal of Experimental Botany 63, 1609–1618.

European Commission Joint Research Centre. (2024). Persistent droughts, critical water shortages, and crops threatened in 2024. Retrieved from https://joint-research-centre.ec.europa.eu/jrc-news-and-updates/persistent-droughts-critical-water-shortages-and-crops-threatened-2024-07-31_en

Exposito-Alonso M., Burbano H.A., Bossdorf O., Nielsen R. & Weigel D. (2019) Natural selection on the *Arabidopsis thaliana* genome in present and future climates. Nature 573, 126–129.

Exposito-Alonso M., Vasseur F., Ding W., Wang G., Burbano H.A. & Weigel D. (2018) Genomic basis and evolutionary potential for extreme drought adaptation in *Arabidopsis thaliana*. Nature Ecology & Evolution 2, 352–358.

Fox H., Ben-Dor S., Doron-Faigenboim A., Goldsmith M., Klein T. & David-Schwartz R. (2023) Carbohydrate dynamics in Populus trees under drought: An expression atlas of genes related to sensing, translocation, and metabolism across organs. Physiologia Plantarum 175. 10.1111/ppl.14001.

Franks S.J., Kane N.C., O’Hara N.B., Tittes S. & Rest J.S. (2016) Rapid genome-wide evolution in *Brassica rapa* populations following drought revealed by sequencing of ancestral and descendant gene pools. Molecular Ecology 25, 3622–3631.

Friedman, J., Hastie, T., & Tibshirani, R. (2010). Regularization paths for generalized linear models via coordinate descent. J. Stat. Softw. 33:1–22.

Fu L., Wang P. & Xiong Y. (2020) Target of Rapamycin Signaling in Plant Stress Responses. Plant Physiology 182, 1613–1623. 10.1104/pp.19.01214.

Fujii H, Zhu J-K (2009) Arabidopsis mutant deficient in 3 abscisic acid-activated protein kinases reveals critical roles in growth, reproduction, and stress. Proc Natl Acad Sci USA 106: 8380–8385.

Gowri Shankar B.A., Sarani R., Michael D., Mridula P., Vasuki Ranjani C., Sowmiya G., … Seka K. (2007) Ion pairs in non-redundant protein structures. Journal of Biosciences 32, 693–704.

George S., Manoharan D., Li J., Britton M. & Parida A. (2017) Drought and salt stress in Chrysopogon zizanioides leads to common and specific transcriptomic responses and may affect essential oil composition and benzylisoquinoline alkaloids metabolism. Current Plant Biology 11–12, 12–22. 10.1016/j.cpb.2017.12.001.

Ghassemian M, Nambara E, Cutler S, Kawaide H, Kamiya Y, McCourt P (2000) Regulation of Abscisic Acid Signaling by the Ethylene Response Pathway in Arabidopsis. Plant Cell 12: 1117–1126.

González E., Danehower D. & Daub M.E. (2007) Vitamer Levels, Stress Response, Enzyme Activity, and Gene Regulation of Arabidopsis Lines Mutant in the Pyridoxine/Pyridoxamine 5′-Phosphate Oxidase (*PDX3*) and the Pyridoxal Kinase (*SOS4*) Genes Involved in the Vitamin B_6_ Salvage Pathway. Plant Physiology 145, 985–996.

Gotarkar D., Longkumer T., Yamamoto N., Nanda A.K., Iglesias T., Li L., … Kohli A. (2021) A drought-responsive rice amidohydrolase is the elusive plant guanine deaminase with the potential to modulate the epigenome. Physiologia Plantarum 172, 1853–1866. 10.1111/ppl.13392

Guan Q., Wen C., Zeng H. & Zhu J. (2013) A KH Domain-Containing Putative RNA-Binding Protein Is Critical for Heat Stress-Responsive Gene Regulation and Thermotolerance in Arabidopsis. Molecular Plant 6, 386–395.

Hahn A., Kilian J., Mohrholz A., Ladwig F., Peschke F., Dautel R., … Wanke D. (2013) Plant Core Environmental Stress Response Genes Are Systemically Coordinated during Abiotic Stresses. International Journal of Molecular Sciences 14, 7617–7641.

He J. & Gai J. (2023) Genome-Wide Association Studies (GWAS). Methods in Molecular Biology, 123–146.

He X.-J., Hsu Y.-F., Pontes O., Zhu J., Lu J., Bressan R.A., … Zhu J.-K. (2009) NRPD4, a protein related to the RPB4 subunit of RNA polymerase II, is a component of RNA polymerases IV and V and is required for RNA-directed DNA methylation. Genes & Development 23, 318–330. 10.1101/gad.1765209.

Hemati A., Moghiseh E., Amirifar A., Mofidi-Chelan M. & Asgari Lajayer B. (2022) Physiological Effects of Drought Stress in Plants. Plant Stress Mitigators, 113–124.

Herrero S., González E., Gillikin J.W., Vélëz H. & Daub M.E. (2011) Identification and characterization of a pyridoxal reductase involved in the vitamin B6 salvage pathway in Arabidopsis. Plant Molecular Biology 76, 157–169.

Hickman, A. B., Chandler, M., & Dyda, F. (2010). Integrating prokaryotes and eukaryotes: DNA transposases in light of structure. Critical Reviews in Biochemistry and Molecular Biology, 45(1), 50–69. 10.3109/10409230903505596.

Hoagland DR, Arnon DI (1938) The water culture method for growing plants without soil. Calif Agric Exp Stn 347, 32.

Hooshyaripor F., Sardari J., Dehghani M. & Noori R. (2022) A new concept of drought feeling against the meteorological drought. Scientific Reports 12.

Horton M.W., Hancock A.M., Huang Y.S., Toomajian C., Atwell S., Auton A., … Bergelson J. (2012) Genome-wide patterns of genetic variation in worldwide *Arabidopsis thaliana* accessions from the RegMap panel. Nature Genetics 44, 212–216. https://agriwelfare.gov.in/Documents/AR_English_2023_24.pdf

Hu R., Zhu Y., Shen G. & Zhang H. (2013) TAP46 Plays a Positive Role in the ABSCISIC ACID INSENSITIVE5-Regulated Gene Expression in Arabidopsis. Plant Physiology 164, 721–734. 10.1104/pp.113.233684.

Huang S., Zhang J., Wang L. & Huang L. (2013) Effect of abiotic stress on the abundance of different vitamin B6 vitamers in tobacco plants. Plant Physiology and Biochemistry 66, 63–67. 10.1016/j.plaphy.2013.02.010.

Huo L., Guo Z., Wang P., Sun X., Xu K. & Ma F. (2021) MdHARBI1, a MdATG8i-interacting protein, plays a positive role in plant thermotolerance. Plant Science 306, 110850.

Jacobs J. & Roe J.L. (2005) SKS6, a multicopper oxidase-like gene, participates in cotyledon vascular patterning during *Arabidopsis thaliana* development. Planta 222, 652–666.

Janssen K.A., Xie Y., Kramer M.C., Gregory B.D. & Garcia B.A. (2022) Data-Independent Acquisition for the Detection of Mononucleoside RNA Modifications by Mass Spectrometry. Journal of the American Society for Mass Spectrometry 33, 885–893.

Jia X., Feng H., Bu Y., Ji N., Lyu Y. & Zhao S. (2021) Comparative Transcriptome and Weighted Gene Co-expression Network Analysis Identify Key Transcription Factors of Rosa chinensis ‘Old Blush’ After Exposure to a Gradual Drought Stress Followed by Recovery. Frontiers in Genetics 12. 10.3389/fgene.2021.690264.

Jiang N., Shi Y., Li M., Du Z., Chen J., Jiang W., … Huang J. (2023) Expression of OsHARBI1-1 enhances the tolerance of *Arabidopsis thaliana* to cadmium. BMC Plant Biology 23.

Jumper, J., Evans, R., Pritzel, A., Green, T., Figurnov, M., Ronneberger, O., Tunyasuvunakool, K., Bates, R., Žídek, A., Potapenko, A., Bridgland, A., Meyer, C., Kohl, S. A. A., Ballard, A. J., Cowie, A., Romera-Paredes, B., Nikolov, S., Jain, R., Adler, J., … Hassabis, D. (2021). Highly accurate protein structure prediction with AlphaFold. Nature, 596(7873), 583–589.

Jungkunz I., Link K., Vogel F., Voll L.M., Sonnewald S. & Sonnewald U. (2011) AtHsp70-15-deficient Arabidopsis plants are characterized by reduced growth, a constitutive cytosolic protein response and enhanced resistance to TuMV. The Plant Journal 66, 983–995.

Kalladan R., Lasky J.R., Chang T.Z., Sharma S., Juenger T.E. & Verslues P.E. (2017) Natural variation identifies genes affecting drought-induced abscisic acid accumulation in *Arabidopsis thaliana*. Proceedings of the National Academy of Sciences 114, 11536–11541.

Khakwani A.A., Dennett M.D., & Munir M. (2011) Early growth response of six wheat varieties under artificial osmotic stress condition. Pakistan Journal of Agricultural Sciences 48(2), 119–123.

Kim J.-S., Mizoi J., Kidokoro S., Maruyama K., Nakajima J., Nakashima K., … Yamaguchi-Shinozaki K. (2012) Arabidopsis GROWTH-REGULATING FACTOR7 Functions as a Transcriptional Repressor of Abscisic Acid– and Osmotic Stress–Responsive Genes, Including DREB2A. The Plant Cell 24, 3393–3405.

Kobayashi Y., Sadhukhan A., Tazib T., Nakano Y., Kusunoki K., Kamara M., … Koyama H. (2016) Joint genetic and network analyses identify loci associated with root growth under NaCl stress in *Arabidopsis thaliana*. Plant, Cell & Environment 39, 918–934.

Kodama Y., Suetsugu N. & Wada M. (2011) Novel protein-protein interaction family proteins involved in chloroplast movement response. Plant Signaling & Behavior 6, 483–490.

Kõressaar T., Lepamets M., Kaplinski L., Raime K., Andreson R. & Remm M. (2018) Primer3_masker: integrating masking of template sequence with primer design software. Bioinformatics 34, 1937–1938.

Kotera E., Tasaka M. & Shikanai T. (2005) A pentatricopeptide repeat protein is essential for RNA editing in chloroplasts. Nature 433, 326–330.

Kunz B.A., Anderson H.J., Osmond M.J. & Vonarx E.J. (2005) Components of nucleotide excision repair and DNA damage tolerance in *Arabidopsis thaliana*. Environmental and Molecular Mutagenesis 45, 115–127. 10.1002/em.20094

Landi S. & Esposito S. (2017) Nitrate Uptake Affects Cell Wall Synthesis and Modeling. Frontiers in Plant Science 8. 10.3389/fpls.2017.01376

Lario L.D., Ramirez-Parra E., Gutierrez C., Casati P. & Spampinato C.P. (2011) Regulation of plant MSH2 and MSH6 genes in the UV-B-induced DNA damage response. Journal of Experimental Botany 62, 2925–2937. 10.1093/jxb/err001

Lee, S., Choi, E., Kim, T., Hwang, J., & Lee, J.-H. (2022). AtHAD1, A haloacid dehalogenase-like phosphatase, is involved in repressing the ABA response. Biochemical and Biophysical Research Communications, 587, 119–125. 10.1016/j.bbrc.2021.11.095

Lee D.W., Kim S.J., Oh Y.J., Choi B., Lee J. & Hwang I. (2016) Arabidopsis BAG1 Functions as a Cofactor in Hsc70-Mediated Proteasomal Degradation of Unimported Plastid Proteins. Molecular Plant 9, 1428–1431.

Lechner E., Achard P., Vansiri A., Potuschak T. & Genschik P. (2006) F-box proteins everywhere. Current Opinion in Plant Biology 9, 631–638.

Legeay M., Doncheva N.T., Morris J.H. & Jensen L.J. (2020) Visualize omics data on networks with Omics Visualizer, a Cytoscape App. F1000Research 9, 157.

Li H., Yang X., Lu M., Chen J. & Shi T. (2020) Gene expression and evolution of Family-1 UDP-glycosyltransferases—insights from an aquatic flowering plant (sacred lotus). Aquatic Botany 166, 103270. 10.1016/j.aquabot.2020.103270

Li L.-H., Lv M.-M., Li X., Ye T.-Z., He X., Rong S.-H., … Xu Z.-J. (2018) The Rice OsDUF810 Family: OsDUF810.7 May be Involved in the Tolerance to Salt and Drought. Molecular Biology 52, 489–496.

Liao Z., Zhang Y., Yu Q., Fang W., Chen M., Li T., … Luo L. (2023) Coordination of growth and drought responses by GA-ABA signaling in rice. New Phytologist 240, 1149–1161. 10.1111/nph.19209

Lin Q., Wang J., Gong J., Zhang Z., Wang S., Sun J., … Qi S. (2023) The *Arabidopsis thaliana* trehalose-6-phosphate phosphatase gene *AtTPPI* improves chilling tolerance through accumulating soluble sugar and JA. Environmental and Experimental Botany 205, 105117. 10.1016/j.envexpbot.2022.105117

Lin Q., Wang S., Dao Y., Wang J. & Wang K. (2020) *Arabidopsis thaliana* trehalose-6-phosphate phosphatase gene *TPPI* enhances drought tolerance by regulating stomatal apertures. Journal of Experimental Botany 71, 4285–4297. 10.1093/jxb/eraa173

Lin Q., Yang J., Wang Q., Zhu H., Chen Z., Dao Y. & Wang K. (2019) Overexpression of the trehalose-6-phosphate phosphatase family gene *AtTPPF* improves the drought tolerance of *Arabidopsis thaliana*. BMC Plant Biology 19. 10.1186/s12870-019-1986-5.

Liu S.-L., Baute G.J. & Adams K.L. (2011) Organ and Cell Type–Specific Complementary Expression Patterns and Regulatory Neofunctionalization between Duplicated Genes in *Arabidopsis thaliana*. Genome Biology and Evolution 3, 1419–1436. 10.1093/gbe/evr114.

Liu, X., Wang, D.-R., Chen, G.-L., Wang, X., Hao, S.-Y., Qu, M.-S., Liu, J.-Y., Wang, X.-F., & You, C.-X. (2024). MdTPR16, an apple tetratricopeptide repeat (TPR)-like superfamily gene, positively regulates drought stress in apple. Plant Physiology and Biochemistry, 210, 108572. 10.1016/j.plaphy.2024.108572

Liu Y., Maniero R.A., Giehl R.F.H., Melzer M., Steensma P., Krouk G., … von Wirén N. (2022) PDX1.1-dependent biosynthesis of vitamin B6 protects roots from ammonium-induced oxidative stress. Molecular Plant 15, 820–839. 10.1016/j.molp.2022.01.012

Livak K.J. & Schmittgen T.D. (2001) Analysis of Relative Gene Expression Data Using Real-Time Quantitative PCR and the 2^−ΔΔCT^ Method. Methods 25, 402–408.

Lu J. & Holmgren A. (2014) The Thioredoxin Superfamily in Oxidative Protein Folding. Antioxidants & Redox Signaling 21, 457–470.

Lundquist P.K., Poliakov A., Bhuiyan N.H., Zybailov B., Sun Q. & van Wijk K.J. (2012) The Functional Network of the Arabidopsis Plastoglobule Proteome Based on Quantitative Proteomics and Genome-Wide Coexpression Analysis. Plant Physiology 158, 1172–1192.

Luo Z., Xiong J., Xia H., Wang L., Hou G., Li Z., … Luo L. (2022) Pentatricopeptide Repeat Gene-Mediated Mitochondrial RNA Editing Impacts on Rice Drought Tolerance. Frontiers in Plant Science 13. 10.3389/fpls.2022.926285.

McDonald I.K., Thornton J.M. (1994) Satisfying hydrogen bonding potential in proteins. Journal of Molecular Biology 238, 777–793.

Mahapatra K. & Roy S. (2020) An insight into the mechanism of DNA damage response in plants-role of SUPPRESSOR OF GAMMA RESPONSE 1: An overview. Mutation Research/Fundamental and Molecular Mechanisms of Mutagenesis 819–820, 111689. 10.1016/j.mrfmmm.2020.111689

Maia J., Dekkers B.J.W., Dolle M.J., Ligterink W. & Hilhorst H.W.M. (2014) Abscisic acid (ABA) sensitivity regulates desiccation tolerance in germinated Arabidopsis seeds. New Phytologist 203, 81–93.

Man Y., Zhang Y., Chen L., Zhou J., Bu Y., Zhang X., … Lin J. (2024) The VAMP-associated protein VAP27-1 plays a crucial role in plant resistance to ER stress by modulating ER–PM contact architecture in Arabidopsis. Plant Communications 5, 100929.

Marik D. & Sadhukhan A. (2025) Unearthing the hidden organ: vital role of the root in drought tolerance of plants. Authorea. 10.22541/au.174348393.34907001/v1

Marik D., Sharma P., Chauhan N.S., Jangir N., Shekhawat R.S., Verma D., … Sadhukhan A. (2024) *Peribacillus frigoritolerans* T7-IITJ, a potential biofertilizer, induces plant growth-promoting genes of *Arabidopsis thaliana*. Journal of Applied Microbiology 135.

Marondedze C., Thomas L., Lilley K.S. & Gehring C. (2020) Drought Stress Causes Specific Changes to the Spliceosome and Stress Granule Components. Frontiers in Molecular Biosciences 6.

Martínez L. (2015) Automatic Identification of Mobile and Rigid Substructures in Molecular Dynamics Simulations and Fractional Structural Fluctuation Analysis. PLOS ONE 10, e0119264.

Martinez-Seidel F., Beine-Golovchuk O., Hsieh Y.-C. & Kopka J. (2020) Systematic Review of Plant Ribosome Heterogeneity and Specialization. Frontiers in Plant Science 11. 10.3389/fpls.2020.00948

Meijón M., Satbhai S.B., Tsuchimatsu T. & Busch W. (2013) Genome-wide association study using cellular traits identifies a new regulator of root development in Arabidopsis. Nature Genetics 46, 77–81.

Mergner J., Frejno M., List M., Papacek M., Chen X., Chaudhary A., … Kuster B. (2020) Mass-spectrometry-based draft of the Arabidopsis proteome. Nature 579, 409–414.

Mittler R. & Blumwald E. (2015) The Roles of ROS and ABA in Systemic Acquired Acclimation. The Plant Cell 27, 64–70.

Mooney S., Leuendorf J.-E., Hendrickson C. & Hellmann H. (2009) Vitamin B6: A Long Known Compound of Surprising Complexity. Molecules 14, 329–351. 10.3390/molecules14010329

Morales-Herrera S., Jourquin J., Coppé F., Lopez-Galvis L., De Smet T., Safi A., … Beeckman T. (2023) Trehalose-6-phosphate signaling regulates lateral root formation in *Arabidopsis thaliana*. Proceedings of the National Academy of Sciences 120. 10.1073/pnas.2302996120

Mukherjee A., Dwivedi S., Bhagavatula L. & Datta S. (2023) Integration of light and ABA signaling pathways to combat drought stress in plants. Plant Cell Reports 42, 829–841.

Munns R. & Tester M. (2008) Mechanisms of Salinity Tolerance. Annual Review of Plant Biology 59, 651–681. 10.1146/annurev.arplant.59.032607.092911

Nakagawa N., Kato M., Takahashi Y., Shimazaki K., Tamura K., Tokuji Y., … Imai H. (2011) Degradation of long-chain base 1-phosphate (LCBP) in Arabidopsis: functional characterization of LCBP phosphatase involved in the dehydration stress response. Journal of Plant Research 125, 439–449. 10.1007/s10265-011-0451-9

Namba J., Harada M., Shibata R., Toda Y., Maruta T., Ishikawa T., … Ogawa T. (2024) AtDREB2G is involved in the regulation of riboflavin biosynthesis in response to low-temperature stress and abscisic acid treatment in *Arabidopsis thaliana*. Plant Science 347, 112196. 10.1016/j.plantsci.2024.112196

Nie J., Stewart R., Zhang H., Thomson J.A., Ruan F., Cui X. & Wei H. (2011) TF-Cluster: A pipeline for identifying functionally coordinated transcription factors via network decomposition of the shared coexpression connectivity matrix (SCCM). BMC Systems Biology 5.

Nisa M.-U., Huang Y., Benhamed M. & Raynaud C. (2019) The Plant DNA Damage Response: Signaling Pathways Leading to Growth Inhibition and Putative Role in Response to Stress Conditions. Frontiers in Plant Science 10.

Nuccio M.L., Wu J., Mowers R., Zhou H.-P., Meghji M., Primavesi L.F., … Lagrimini L.M. (2015) Expression of trehalose-6-phosphate phosphatase in maize ears improves yield in well-watered and drought conditions. Nature Biotechnology 33, 862–869. 10.1038/nbt.3277

Ono K., Fong D., Gao C., Churas C., Pillich R., Lenkiewicz J., … Chen J. (2025) Cytoscape Web: bringing network biology to the browser. Nucleic Acids Research. gkaf365, 10.1093/nar/gkaf365

Pavoković D., Horvatić A., Tomljanović I., Balen B. & Krsnik-Rasol M. (2023) Sugar beet cells cellular and extracellular events taking place in response to drought and salinity. Acta botanica Croatica 82, 128–141. 10.37427/botcro-2023-008

Pixley K.V., Cairns J.E., Lopez-Ridaura S., Ojiewo C.O., Dawud M.A., Drabo I., … Zepeda-Villarreal E.A. (2023) Redesigning crop varieties to win the race between climate change and food security. Molecular Plant 16, 1590–1611.

Ponder, J. W., & Case, D. A. (2003). Force Fields for Protein Simulations. In Protein Simulations (pp. 27–85). Elsevier.

Punzo P., Ruggiero A., Possenti M., Nurcato R., Costa A., Morelli G., … Batelli G. (2018) The PP2A-interactor TIP41 modulates ABA responses in *Arabidopsis thaliana*. The Plant Journal 94, 991–1009. 10.1111/tpj.13913

Rajeswar S. & S N. (2021) PEG-induced Drought Stress in Plants: A Review. Research Journal of Pharmacy and Technology, 6173–6178.

Ruiz-Aguilar B., Torres-Serrallonga N.B., Ortega-Amaro M.A., Duque-Ortiz A., Ovando-Vázquez C. & Jiménez-Bremont J.F. (2024) Transcriptome Analysis Reveals Genes Responsive to Three Low-Temperature Treatments in *Arabidopsis thaliana*. Plants 13, 3127. 10.3390/plants13223127

Sadhukhan A., Kobayashi Y., Kobayashi Y., Tokizawa M., Yamamoto Y.Y., Iuchi S., … Sahoo L. (2014) VuDREB2A, a novel DREB2-type transcription factor in the drought-tolerant legume cowpea, mediates DRE-dependent expression of stress-responsive genes and confers enhanced drought resistance in transgenic Arabidopsis. Planta 240, 645–664.

Sadhukhan A., Kobayashi Y., Nakano Y., Iuchi S., Kobayashi M., Sahoo L. & Koyama H. (2017) Genome-wide Association Study Reveals that the Aquaporin NIP1;1 Contributes to Variation in Hydrogen Peroxide Sensitivity in *Arabidopsis thaliana*. Molecular Plant 10, 1082–1094.

Sakhare A.S., Kumar S., Ellur R.K., Prahalada G.D., Kota S., Kumar R.R., … Chinnusamy V. (2024) Genome-wide association study on root traits under non-stress and osmotic stress conditions to improve drought tolerance in rice (*Oryza sativa* Lin.). Acta Physiologiae Plantarum 47.

Sampaio M., Neves J., Cardoso T., Pissarra J., Pereira S. & Pereira C. (2022) Coping with Abiotic Stress in Plants— An Endomembrane Trafficking Perspective. Plants 11, 338.

Sang Y., Locy R.D., Goertzen L.R., Rashotte A.M., Si Y., Kang K. & Singh N.K. (2011) Expression, in vivo localization and phylogenetic analysis of a pyridoxine 5′-phosphate oxidase in *Arabidopsis thaliana*. Plant Physiology and Biochemistry 49, 88–95. 10.1016/j.plaphy.2010.10.003

Schippers J.H.M., von Bongartz K., Laritzki L., Frohn S., Frings S., Renziehausen T., … Schmidt-Schippers R.R. (2024) ERFVII-controlled hypoxia responses are in part facilitated by MEDIATOR SUBUNIT 25 in *Arabidopsis thaliana*. The Plant Journal 120, 748–768. 10.1111/tpj.17018

Schmid M.W., Heichinger C., Coman Schmid D., Guthörl D., Gagliardini V., Bruggmann R., … Grossniklaus U. (2018) Contribution of epigenetic variation to adaptation in Arabidopsis. Nature Communications 9.

Schulz E., Tohge T., Zuther E., Fernie A.R. & Hincha D.K. (2016) Flavonoids are determinants of freezing tolerance and cold acclimation in *Arabidopsis thaliana*. Scientific Reports 6.

Science Advances. (2024). Unprecedented fall drought leads to Dust Bowl-like crop losses in 2024. Science, 10(4), eado6864. https://www.science.org/doi/10.1126/sciadv.ado6864

Sagisaka S. (1976) The Occurrence of Peroxide in a Perennial Plant, Populus gelrica. Plant Physiology 57, 308–309. 10.1104/pp.57.2.308

Shannon P., Markiel A., Ozier O., Baliga N.S., Wang J.T., Ramage D., … Ideker T. (2003) Cytoscape: A Software Environment for Integrated Models of Biomolecular Interaction Networks. Genome Research 13, 2498–2504.

Siaud N., Dubois E., Massot S., Richaud A., Dray E., Collier J. & Doutriaux M.-P. (2010) The SOS screen in Arabidopsis: A search for functions involved in DNA metabolism. DNA Repair 9, 567–578. 10.1016/j.dnarep.2010.02.009

Şimşek Ö., Isak M.A., Dönmez D., Dalda Şekerci A., İzgü T. & Kaçar Y.A. (2024) Advanced Biotechnological Interventions in Mitigating Drought Stress in Plants. Plants 13, 717.

Small I.D. & Peeters N. (2000) The PPR motif – a TPR-related motif prevalent in plant organellar proteins. Trends in Biochemical Sciences 25, 45–47. doi: 10.1016/s0968-0004(99)01520-0

Sproule, Ann K. (2008) Microtubule involvement in the plant low temperature response. Diss. University of Saskatchewan.

Stief A., Altmann S., Hoffmann K., Pant B.D., Scheible W.-R. & Bäurle I. (2014) Arabidopsis miR156 Regulates Tolerance to Recurring Environmental Stress through SPL Transcription Factors. The Plant Cell 26, 1792–1807. 10.1105/tpc.114.123851

Su H.-G., Li B., Song X.-Y., Ma J., Chen J., Zhou Y.-B., … Ma Y.-Z. (2019) Genome-Wide Analysis of the DYW Subgroup PPR Gene Family and Identification of GmPPR4 Responses to Drought Stress. International Journal of Molecular Sciences 20, 5667. 10.3389/fpls.2022.926285

Szklarczyk D., Kirsch R., Koutrouli M., Nastou K., Mehryary F., Hachilif R., … von Mering C. (2022) The STRING database in 2023: protein–protein association networks and functional enrichment analyses for any sequenced genome of interest. Nucleic Acids Research 51, D638–D646.

Uffelmann E., Huang Q.Q., Munung N.S., de Vries J., Okada Y., Martin A.R., … Posthuma D. (2021) Genome-wide association studies. Nature Reviews Methods Primers 1: 59. 10.1038/s43586-021-00056-9

Umezawa T, Nakashima K, Miyakawa T, Kuromori T, Tanokura M, Shinozaki K, Yamaguchi-Shinozaki K (2010) Molecular Basis of the Core Regulatory Network in ABA Responses: Sensing, Signaling and Transport. Plant and Cell Physiology 51: 1821–1839

UniProt Consortium. (2025). UniProt: the Universal Protein Knowledgebase in 2025. Nucleic Acids Research, 53(D1), D609–D617.

Virk N., Li D., Tian L., Huang L., Hong Y., Li X., … Song F. (2015) Arabidopsis Raf-Like Mitogen-Activated Protein Kinase Kinase Kinase Gene Raf43 Is Required for Tolerance to Multiple Abiotic Stresses. PLOS ONE 10, e0133975.

Voothuluru P., Wu Y. & Sharp R.E. (2024) Not so hidden anymore: Advances and challenges in understanding root growth under water deficits. The Plant Cell 36, 1377–1409.

Vu W.T., Chang P.L., Moriuchi K.S. & Friesen M.L. (2015) Genetic variation of transgenerational plasticity of offspring germination in response to salinity stress and the seed transcriptome of *Medicago truncatula*. BMC Evolutionary Biology 15.

Wagner S., Bernhardt A., Leuendorf J.E., Drewke C., Lytovchenko A., Mujahed N., … Hellmann H. (2006) Analysis of the Arabidopsis *rsr4-1/pdx1-3* Mutant Reveals the Critical Function of the PDX1 Protein Family in Metabolism, Development, and Vitamin B6 Biosynthesis. The Plant Cell 18, 1722–1735. 10.1105/tpc.105.036269.

Wang D., Liu X., He G., Wang K., Li Y., Guan H., … Li Y. (2025) GWAS and transcriptome analyses unravel *ZmGRAS15* regulates drought tolerance and root elongation in maize. BMC Genomics 26.

Wang H., Liu D., Liu C. & Zhang A. (2004) The pyridoxal kinase gene *TaPdxK* from wheat complements vitamin B6 synthesis-defective *Escherichia coli*. Journal of Plant Physiology 161, 1053–1060. 10.1016/j.jplph.2004.05.012.

Wang X., An Y., Xu P. & Xiao J. (2021) Functioning of PPR Proteins in Organelle RNA Metabolism and Chloroplast Biogenesis. Frontiers in Plant Science 12.

Wang Y. & Tan B.-C. (2025) Pentatricopeptide repeat proteins in plants: Cellular functions, action mechanisms, and potential applications. Plant Communications 6, 101203.

Wang Z., Wang M., Yang C., Zhao L., Qin G., Peng L., … Zhao C. (2021) SWO1 modulates cell wall integrity under salt stress by interacting with importin ɑ in Arabidopsis. Stress Biology 1.

Xiong J., Mi X., Du L. & Wang X. (2025) The LBD Transcription Factor ZmLBD33 Confers Drought Tolerance in Transgenic Arabidopsis. Plants 14, 1305. 10.3390/plants14091305.

Xu L., Wang J., Lei M., Li L., Fu Y., Wang Z., … Li Z. (2016) Transcriptome Analysis of Storage Roots and Fibrous Roots of the Traditional Medicinal Herb *Callerya speciosa* (Champ.) ScHot. PLOS ONE 11, e0160338. 10.1371/journal.pone.0160338.

Yadav S., Yadava Y.K., Meena S., Kalwan G., Bharadwaj C., Paul V., … Jain P.K. (2024) Novel insights into drought-induced regulation of ribosomal genes through DNA methylation in chickpea. International Journal of Biological Macromolecules 266, 131380. 10.1016/j.ijbiomac.2024.131380.

Yamamoto Y.Y., Yoshioka Y., Hyakumachi M., Maruyama K., Yamaguchi-Shinozaki K., Tokizawa M. & Koyama H. (2011) Prediction of transcriptional regulatory elements for plant hormone responses based on microarray data. BMC Plant Biology 11, 39.

Yan Y., Gan J., Tao Y., Okita T.W. & Tian L. (2022) RNA-Binding Proteins: The Key Modulator in Stress Granule Formation and Abiotic Stress Response. Frontiers in Plant Science 13.

Yang G., Yu Z., Gao L. & Zheng C. (2019) SnRK2s at the Crossroads of Growth and Stress Responses. Trends in Plant Science 24, 672–676. 10.1016/j.tplants.2019.05.010.

Yi K., Menand B., Bell E. & Dolan L. (2010) A basic helix-loop-helix transcription factor controls cell growth and size in root hairs. Nature Genetics 42, 264–267.

Yin R., Han K., Heller W., Albert A., Dobrev P.I., Zažímalová E. & Schäffner A.R. (2013) Kaempferol 3-O-rhamnoside-7-O-rhamnoside is an endogenous flavonol inhibitor of polar auxin transport in Arabidopsis shoots. New Phytologist 201, 466–475. 10.1111/nph.12558.

Yoshida T, Nishimura N, Kitahata N, Kuromori T, Ito T, Asami T, Shinozaki K, Hirayama T (2005) ABA-Hypersensitive Germination3Encodes a Protein Phosphatase 2C (AtPP2CA) That Strongly Regulates Abscisic Acid Signaling during Germination among Arabidopsis Protein Phosphatase 2Cs. Plant Physiology 140: 115–126.

Ytterberg A.J., Peltier J.-B. & van Wijk K.J. (2006) Protein Profiling of Plastoglobules in Chloroplasts and Chromoplasts. A Surprising Site for Differential Accumulation of Metabolic Enzymes. Plant Physiology 140, 984–997.

Yu J., Sun H., Zhang J., Hou Y., Zhang T., Kang J., … Long R. (2020) Analysis of Aldo–Keto Reductase Gene Family and Their Responses to Salt, Drought, and Abscisic Acid Stresses in *Medicago truncatula*. International Journal of Molecular Sciences 21, 754.

Zan, X., Zhou, Z., Wan, J., Chen, H., Zhu, J., Xu, H., Zhang, J., Li, X., Gao, X., Chen, R., Huang, Z., Xu, Z., & Li, L. (2023). Overexpression of OsHAD3, a Member of HAD Superfamily, Decreases Drought Tolerance of Rice. Rice, 16(1). 10.1186/s12284-023-00647-y.

Zepner L., Karrasch P., Wiemann F. & Bernard L. (2020) ClimateCharts.net – an interactive climate analysis web platform. International Journal of Digital Earth 14, 338–356.

Zhang H., Liu P., Guo T., Zhao H., Bensaddek D., Aebersold R. & Xiong L. (2019) Arabidopsis Proteome and the Mass Spectral Assay Library. bioRxiv preprint doi: 10.1101/665547.

Zhao L., Bai T., Wei H., Gardea-Torresdey J.L., Keller A. & White J.C. (2022) Nanobiotechnology-based strategies for enhanced crop stress resilience. Nature Food 3, 829–836.

Zhou W., Karcher D. & Bock R. (2013) Importance of adenosine-to-inosine editing adjacent to the anticodon in an Arabidopsis alanine tRNA under environmental stress. Nucleic Acids Research 41, 3362–3372.

Zhou W., Karcher D. & Bock R. (2014) Identification of Enzymes for Adenosine-to-Inosine Editing and Discovery of Cytidine-to-Uridine Editing in Nucleus-Encoded Transfer RNAs of Arabidopsis. Plant Physiology 166, 1985–1997.

Zhu F., Luo T., Liu C., Wang Y., Zheng L., Xiao X., … Cheng Y. (2020) A NAC transcription factor and its interaction protein hinder abscisic acid biosynthesis by synergistically repressing NCED5 in Citrus reticulata. Journal of Experimental Botany 71, 3613–3625.

