## Supplementary data for "Genome-wide association study, network analysis, and reverse genetics pinpoint novel genes associated with seedling root growth variation of *Arabidopsis thaliana* under drought"

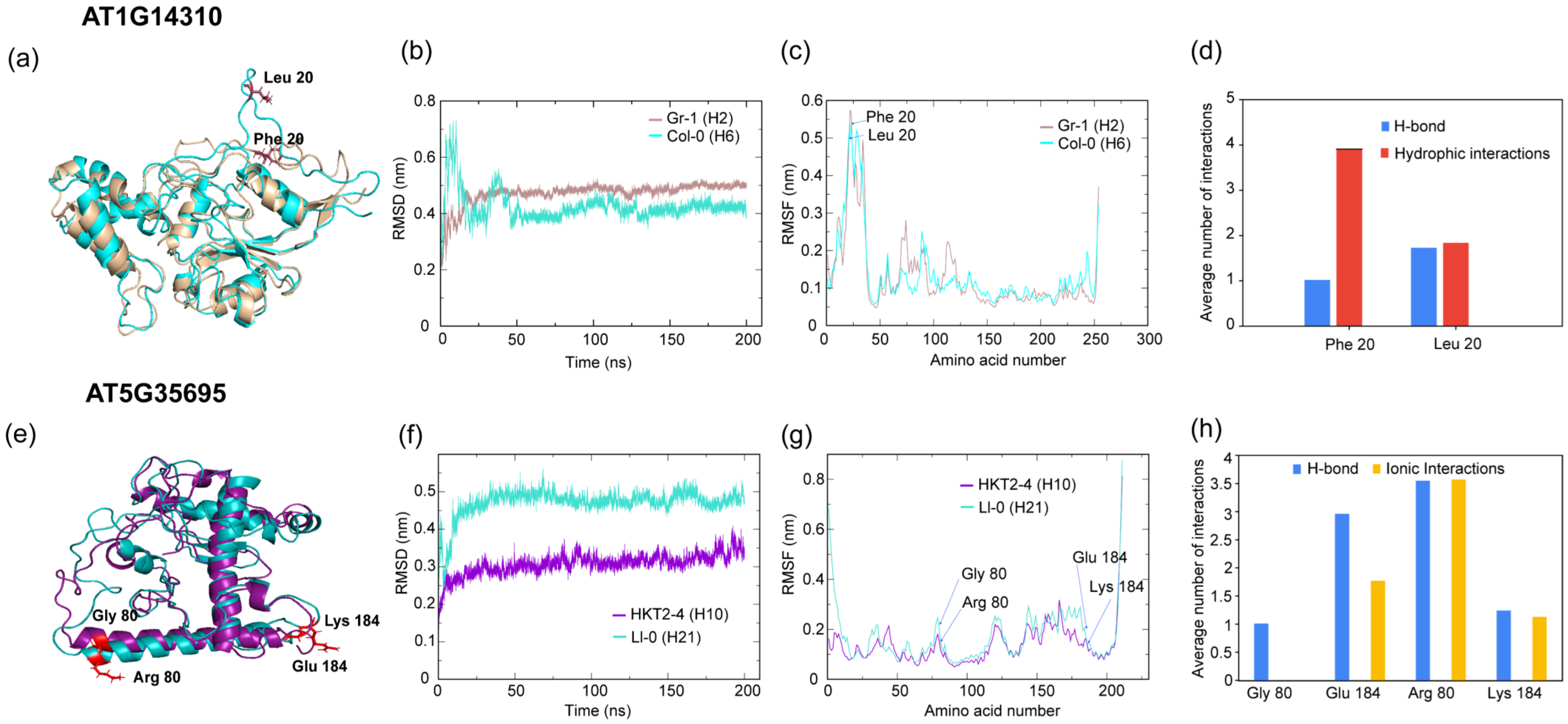


**Fig. S1. Effect of amino acid substitutions on predicted protein molecular dynamics.** The moderate-impact amino acid substitutions in two top-ranking GWAS-delineated genes AT1G14310 and AT5G35695 were identified by the POLYMORPH 1000 genome variants web tool (http://tools.1001genomes.org/polymorph/), and amino acid sequence haplotypes (denoted by H1, H2, etc) were constructed (Table 3, S4). (**a, e**) AlphaFold (https://alphafold.ebi.ac.uk/)-predicted three-dimensional structures of representative haplotypes with the most amino acid substitutions are presented after molecular dynamics (MD) simulation as structural superpositions**.** Backbone Root Mean Square Deviation (RMSD) plots over the simulation time show how much the protein's backbone structure deviates from its initial conformation, reflecting the overall structural stability and changes during the MD simulation (**b, f**). Backbone Root Mean Square Fluctuation (RMSF) profiles plotted across residue numbers highlight the flexibility and dynamic behavior of individual residues during the MD simulation **(c, g).** Predicted intramolecular interactions (hydrogen bonds, hydrophobic interactions, and ionic interactions) between the polymorphic amino acids with their neighbouring residues are shown as bar graphs (**d, h**). The average number of interactions was calculated as the total interaction count divided by the number of frames in which the interaction was observed.

### Fig. S2 Validation of homozygosity of the TDNA insertion mutants used in the study (Contd..)

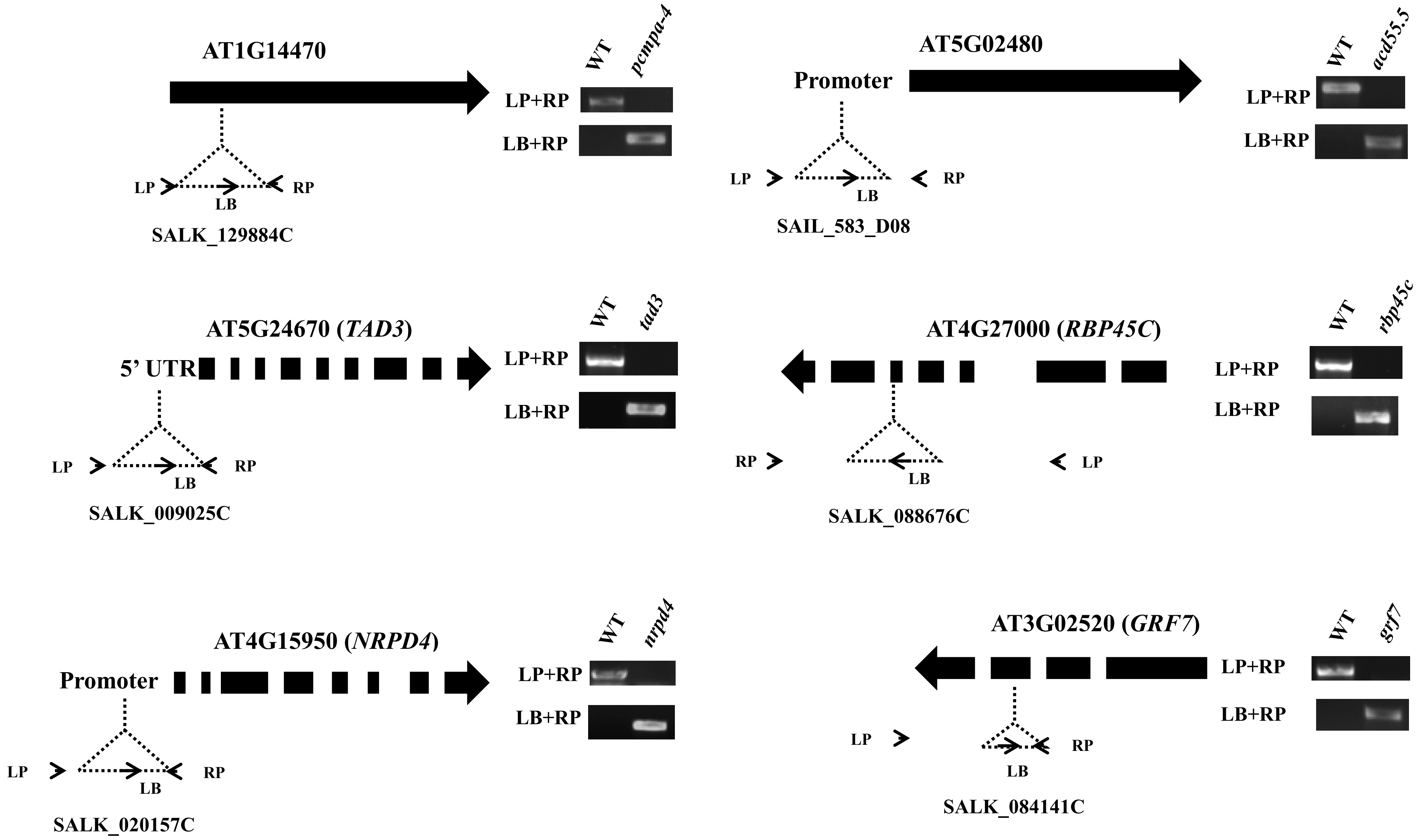


Continued…

**Fig. S2 Validation of homozygosity of the TDNA insertion mutants used in the study (Contd..)**

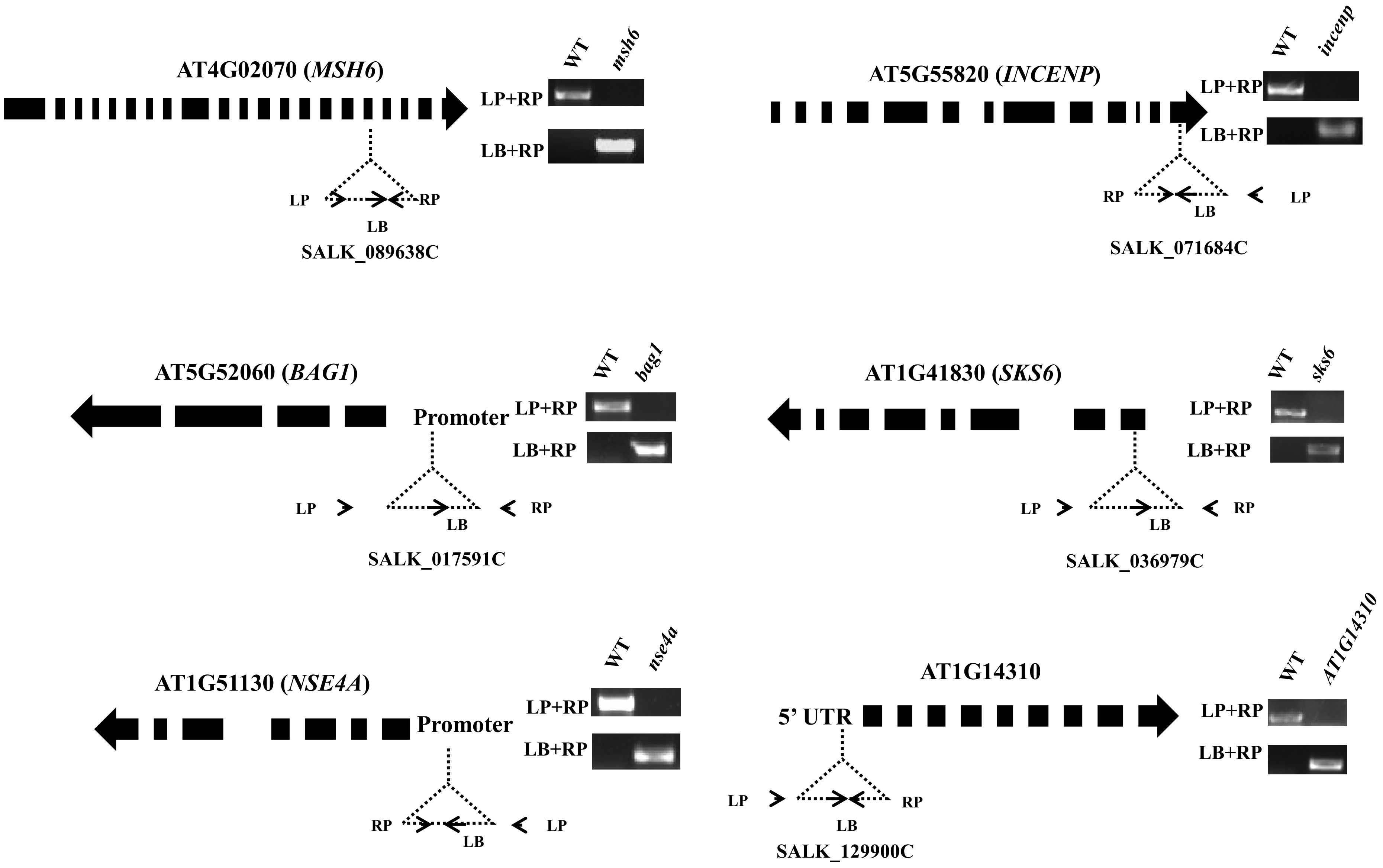


**Continued…**

### Fig. S2 Validation of homozygosity of the TDNA insertion mutants used in the study

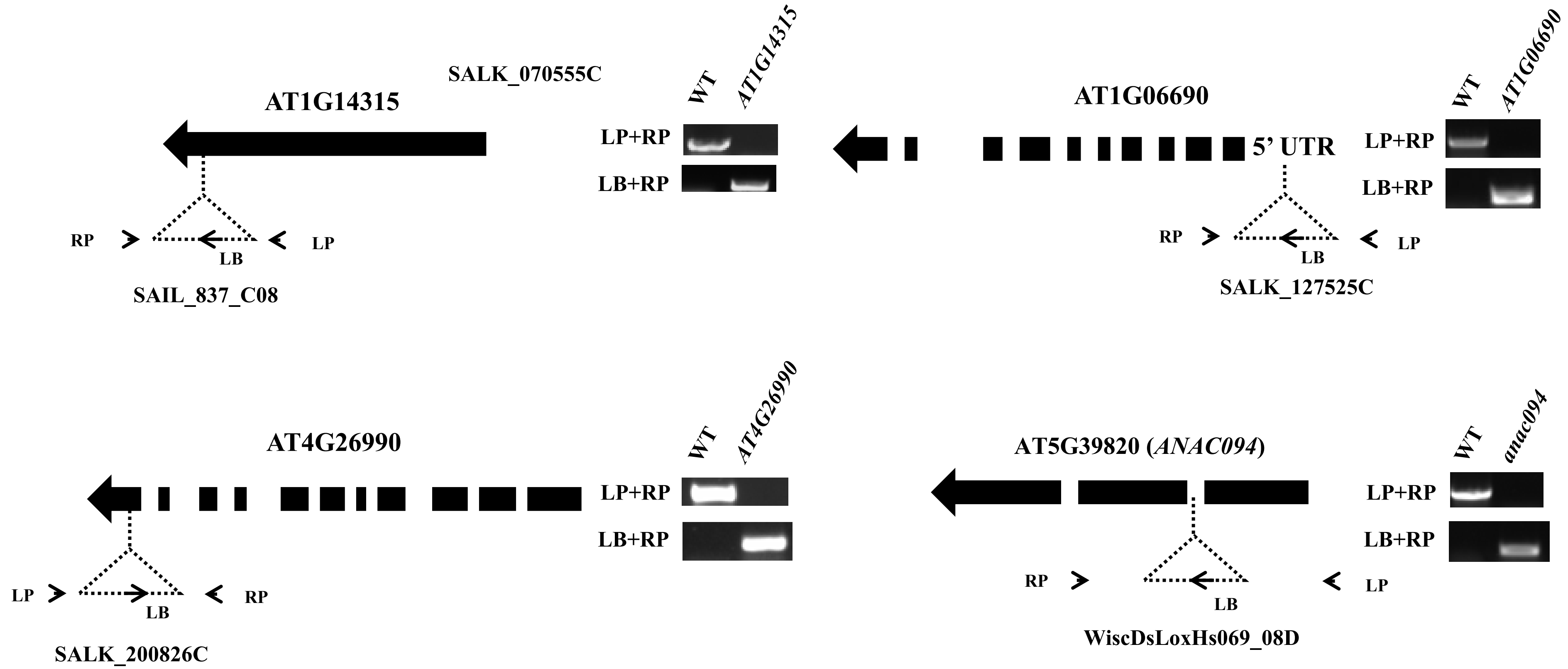


**Fig. S2 Validation of homozygosity of the TDNA insertion mutants used in the study**. TDNA insertion mutants used in the study, viz., *pcmpa-4* (SALK_129884C), *acd55.5* (SAIL_583_D08), *tad3* (SALK_009025C), *rbp45c* (SALK_088676C), *nrpd4* (SALK_020157C), *grf7* (SALK_084141C), *msh6* (SALK_089638C), *incenp* (SALK_071684C), *bag1* (SALK_017591C), *sks6* (SALK_036979C), *nse4a* (SALK_070555C), AT1G14310 (SALK_129900C), AT1G14315 (SAIL_837_C08), AT1G06690 (SALK_127525C), AT4G26990 (SALK_200826C), *anac094* (WiscDsLoxHs069_08D) were tested for homozygosity by PCR-based method. The primer list is available in Table S3. Position of the T-DNA inserted is represented by dotted triangles where T-DNA left border (LB), forward genomic primer (LP), and reverse genomic primer (RP) are labelled along with gel images of amplification. Gene coding sequences are denoted by thick black arrows in the 5' to 3' direction, where intermediate gaps indicate introns.

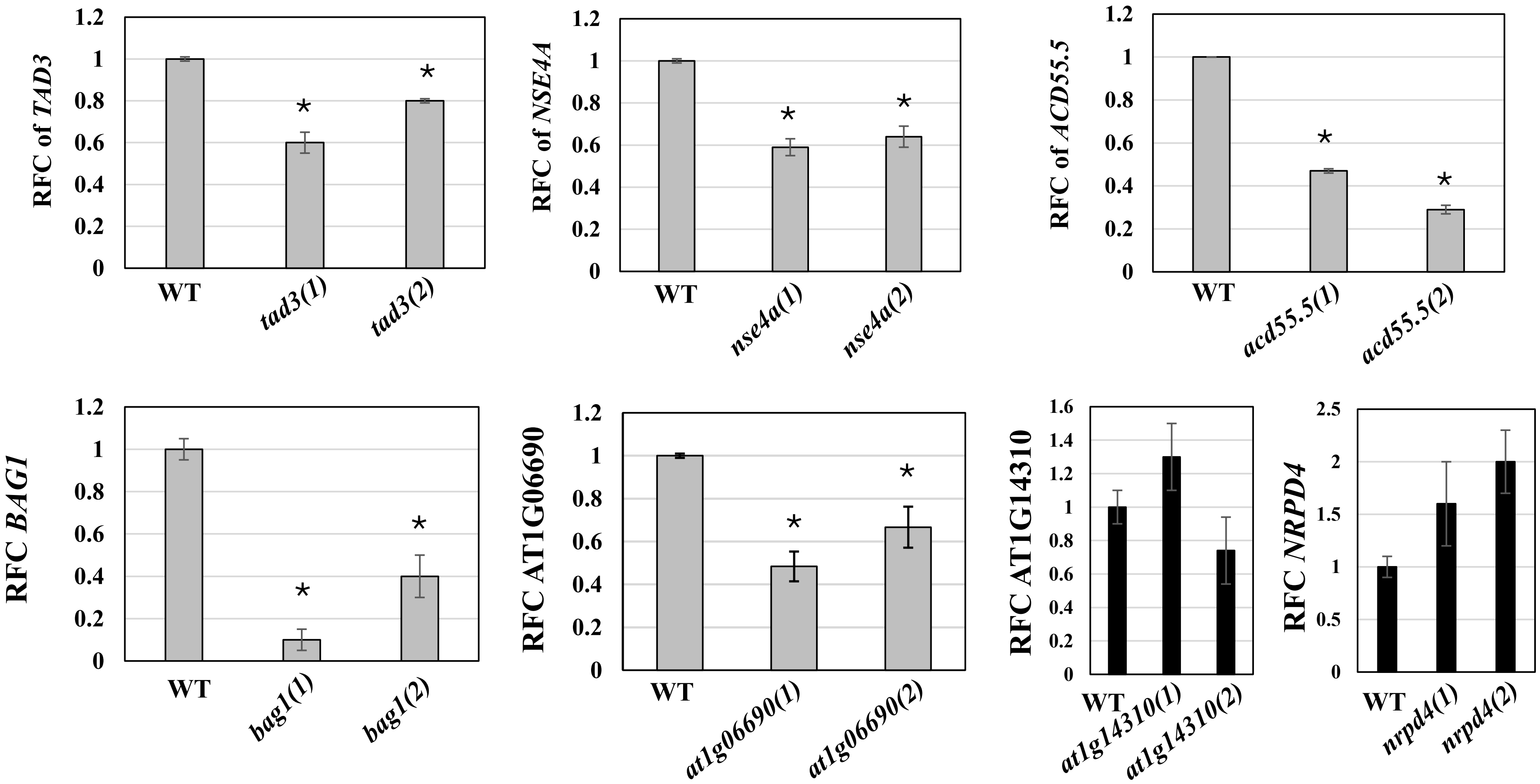


**Fig. S3 Gene expression levels of mutants with T-DNA insertion in the promoter and 5' UTR**. T-DNA insertion events in the promoter or 5' UTR were tested for expression level reduction. The confirmed homozygous mutants: *at1g14310* (SALK_129900C), *nrpd4* (SALK_020157C), *tad3* (SALK_009025C), *nse4a* (SALK_070555C), *acd55.5* (SAIL_583_D08), *bag1* (SALK_017591C), and *at1g06690* (SALK_127525C) were grown in ¼-Hoagland's media for 10 days from germination, followed by 5 days of drought stress in ¼-Hoagland's media supplemented with 2.5% PEG. The seedlings were harvested, followed by RNA isolation and qPCR analysis. Fold change in expression levels of two independent knockdown lines with respect to the wild type (WT) is represented. *AtUBQ1* was used as an internal control for normalization of expression levels, and fold change of expression was calculated according to the $2^{-\Delta\Delta Ct}$ method. Asterisks indicate significant differences between WT and knockdown lines (Student's t-test, $N = 3$ biological replicates).

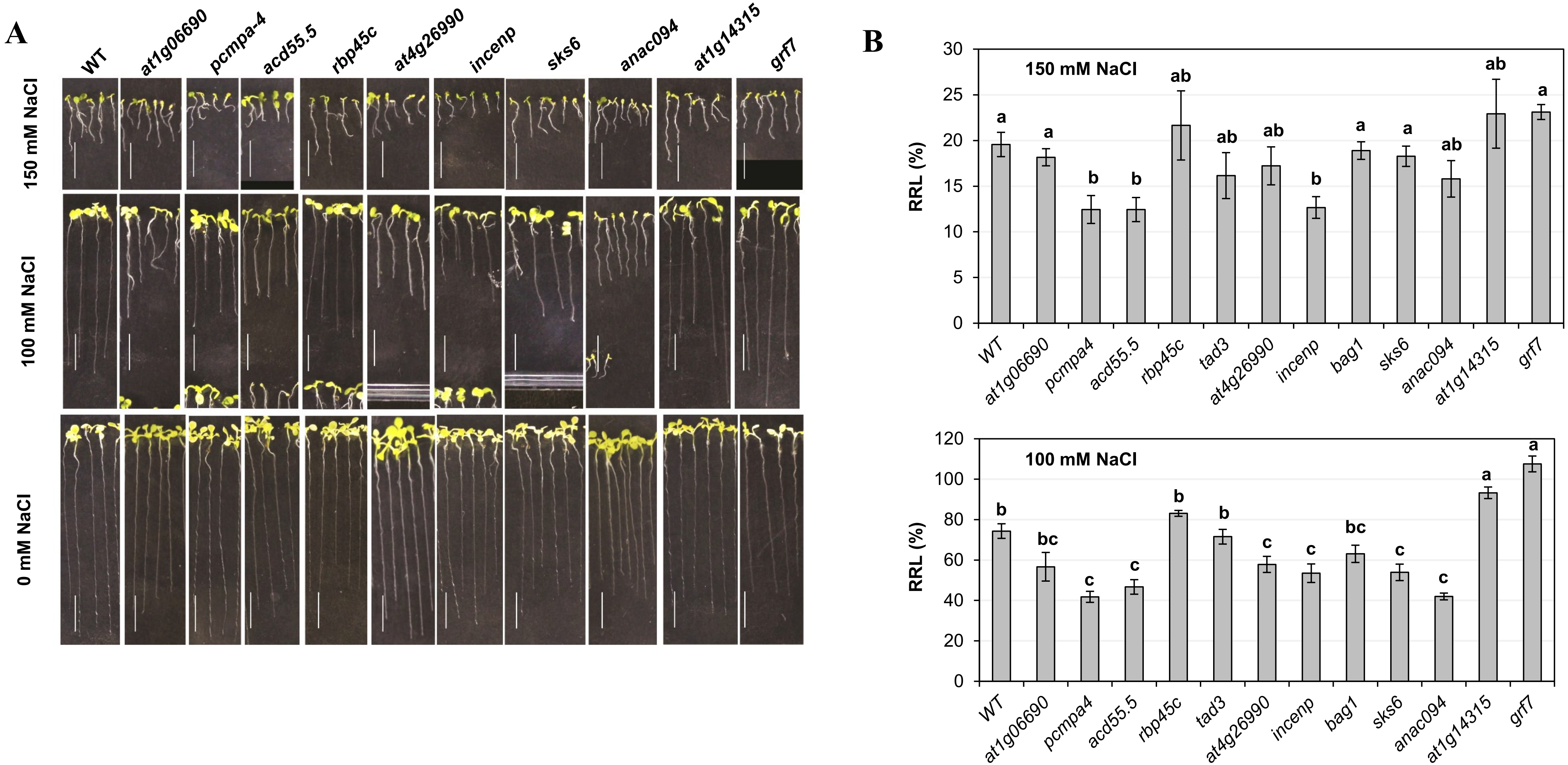


**Fig. S4 Evaluation of salt tolerance of T-DNA insertion mutants used in the study. (A)**Wild-type (Col-0) and T-DNA mutants: *at1g06690*, *pcmpa-4*, *acd55.5, rbp45c*, *at4g26990*, *tad3*, *incenp*, *bag1*, *sks6*, *anac094*, *at1g14315*, *grf7* (see Fig. 7) were grown in ½ MS media with 3% sucrose alone (0 mM) or supplemented with 100 mM or 150 mM NaCl for 14 days and photographed. Scale bar = 1 cm. **(B)** Relative root length (RRL) was calculated by dividing the average root length of stressed plants by the control (0 mM) plants and expressed as a percentage. The average RRL of the top 8 out of 20 plants is shown with standard error. Significant differences ($P < 0.05$; Tukey's test) are shown by different lowercase letters above the bars.

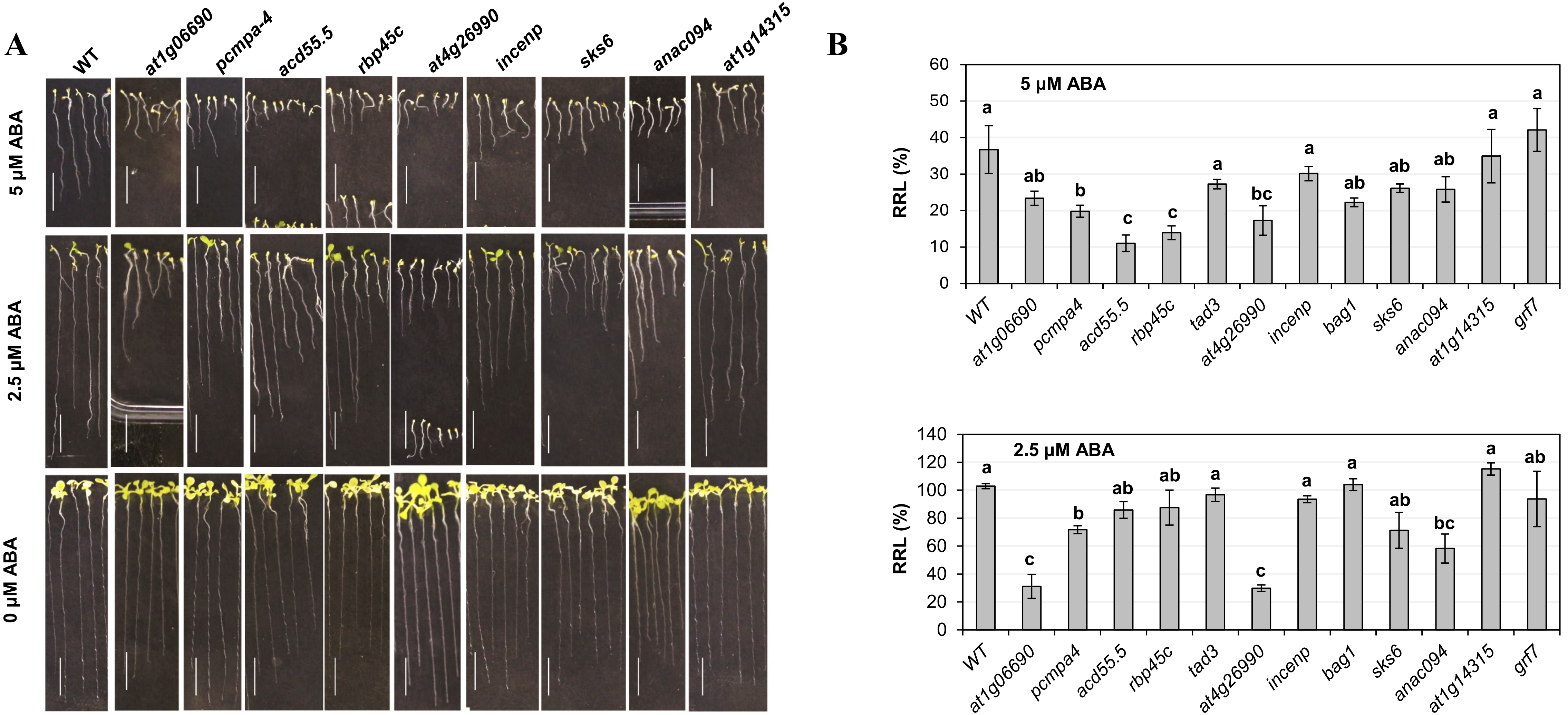


**Fig. S5 Evaluation of ABA sensitivity of T-DNA insertion mutants used in the study. (A)**Wild-type (Col-0) and T-DNA mutants: *at1g06690*, *pcmpa-4*, *acd55.5, rbp45c*, *at4g26990*, *tad3*, *incenp*, *bag1*, *sks6*, *anac094*, *at1g14315*, *grf7* (see Fig. 7) were grown in ½ MS media with 3% sucrose alone (0 mM) or supplemented with 2.5 µM or 5 µM abscisic acid (ABA) for 14 days and photographed. Scale bar = 1 cm. **(B)** Relative root length (RRL) was calculated by dividing the average root length of ABA-treated plants by the control (0 mM) plants and expressed as a percentage. The average RRL of the top 8 out of 20 plants is shown with standard error. Significant differences ($P < 0.05$; Tukey's test) are shown by different lowercase letters above the bars.

### Table S1. List of ecotypes used in the study

| Ecotype | Drought tolerance (RRL; %) | Latitude | Longitude | Geographical location | Annual precipitation (1901-2022) in mm | Average annual temperature (°C) |
|---|---|---|---|---|---|---|
| Rsch-0 | 3.38 | 45.5333 | 4.85 | GRJX+2X5 Saint-Romain-en-Gal, France | 837 | 11 |
| Pla-3 | 7.3 | 41.5 | 2.25 | G722+22X La Conreria, Spain | 727 | 13 |
| Tu-0 | 8.99 | 45 | 7.5 | 2G22+225 Gerbole, Metropolitan City of Turin, Italy | 839 | 11.4 |
| Ts-1 | 9.64 | 41.7194 | 2.93056 | PW9J+X2R Tossa de Mar, Spain | 767 | 14.2 |
| Yeg-1 | 9.66 | 39.8692 | 45.3622 | V9C6+222 Yeghegis, Armenia | 439 | 7.7 |
| Bsch-0 | 9.87 | 50.0167 | 8.6667 | 2M99+XXX Dreieich, Germany | 643 | 9.5 |
| Ra-0 | 9.98 | 46 | 3.3 | 2822+222 Saint-Clément-de-Régnat, France | 733 | 10.5 |
| Tu-SB30-3 | 10.04 | 48.53 | 9.06 | G3J6+222 Tübingen, Germany | 691 | 9.4 |
| Buckhorn Pass | 10.29 | 41.3599 | -122.755 | 966R+222 Callahan, California, USA | 1129 | 7.1 |
| Star-8 | 10.4 | 48.43 | 8.82 | CRJ9+2X5 Starzach, Germany | 940 | 8.1 |
| TueV-13 | 10.68 | 48.52 | 9.05 | G392+X2R Tübingen, Germany | 691 | 9.4 |
| HKT2-4 | 10.86 | 48.14 | 9.4 | 4CQ2+X2R Langenenslingen, Germany | 909 | 7.8 |
| Rennes-1 | 10.98 | 48.5 | -1.41 | GH2R+22R Saint-James, France | 815 | 11 |
| Ty-0 | 11.92 | 56.4278 | -5.23439 | CQJC+222 Taynuilt, United Kingdom | 2743 | 7.8 |
| Ak-1 | 11.96 | 48.0683 | 7.62551 | 3J9H+XXX Vogtsburg, Germany | 745 | 10.1 |
| Uod-7 | 12.06 | 48.3 | 14.45 | 8C2X+2X5 Sankt Georgen an der Gusen, Austria | 946 | 8.3 |
| Pr-0 | 12.81 | 50.1448 | 8.60706 | 4JR6+222 Frankfurt am Main, Germany | 643 | 9.5 |
| Pla-1 | 12.84 | 41.5 | 2.25 | G722+22X La Conreria, Spain | 727 | 13 |
| Pn-0 | 13.62 | 48.0653 | -2.96591 | 329J+X2R Pontivy, France | 940 | 11.1 |
| Bak-7 | 14.03 | 41.7942 | 43.4767 | QFQH+XXX Tsaghveri, Georgia | 705 | 7.4 |
| Sp-0 | 14.62 | 52.5339 | 13.181 | G5JJ+222 Berlin, Germany | 579 | 9.1 |
| Ciste-1 | 14.7 | 41.62 | 12.87 | JVCC+222 Cori, Province of Latina, Italy | 875 | 14.7 |
| Me-0 | 14.77 | 51.9183 | 10.1138 | W496+X2R Seesen, Germany | 870 | 8.1 |
| Pro-0 | 14.85 | 43.25 | -6 | 7222+22R San Martín, Proaza, Spain | 937 | 11.2 |
| Stepn-2 | 15.29 | 54.09 | 60.46 | 3FR5+2X5 Stepnoe, Chelyabinsk Oblast, Russia | 469 | 2.2 |
| Yo-0 | 15.48 | 37.45 | -119.35 | 8592FM22+22, Sierra, California | 736 | 9.4 |
| Alst-1 | 16.3 | 54.8 | -2.4333 | RH29+2X5 Alston, United Kingdom | 1257 | 6.7 |
| Nd-1 | 16.96 | 50 | 10 | 2222+222 Arnstein, Germany | 685 | 8.3 |
| Ag-0 | 17.01 | 45 | 1.3 | 2822+222 Archignac, France | 979 | 11.5 |
| Toufl-1 | 17.31 | 31.47 | -7.42 | FH9J+X2R Toufliht, Morocco | 423 | 14.1 |
| Uk-1 | 17.4 | 48.0333 | 7.7667 | 2QJC+222 Umkirch, Germany | 745 | 10.1 |
| Utrecht | 18.35 | 52.0918 | 5.1145 | 34R6+222 Utrecht, Netherlands | 792 | 9.7 |
| Rd-0 | 18.47 | 50.5 | 8.5 | GG22+22X Hüttenberg, Germany | 693 | 8.9 |
| Aa-0 | 18.72 | 50.9167 | 9.57073 | WH99+XXX Neuenstein, Germany | 709 | 8.2 |
| Br-0 | 18.84 | 49.2 | 16.6166 | 5JXC+X2R Brno, Czechia | 586 | 8.1 |
| Jm-1 | 19.22 | 49 | 15 | 2222+222 Rottal, Austria | 697 | 7 |
| Ms-0 | 19.23 | 55.7522 | 37.6322 | QJ2H+2XX Tverskoy District, Moscow, Russia | 659 | 5 |
| Pog-0 | 19.27 | 49.2655 | -123.206 | 7Q9R+X2R Vancouver, British Columbia, Canada | 1882 | 8.6 |
| Po-1 | 20.15 | 50.7167 | 7.1 | P39X+XXX Bonn, Germany | 770 | 9.9 |
| Wc-1 | 20.17 | 52.6 | 10.0667 | H3X9+XXX Celle, Germany | 681.8 | 8.9 |
| Kl-0 | 21.76 | 50.95 | 6.9666 | WXX9+XXX Cologne, Germany | 836.5 | 9.9 |
| Tha-1 | 21.86 | 52.08 | 4.3 | 38H2+X2R The Hague, Netherlands | 805.8 | 9.9 |
| Ema-1 | 21.92 | 51.3 | 0.5 | 8G22+225 Aylesford, United Kingdom | 687.5 | 10.3 |
| Nz-1 | 21.93 | -37.7871 | 175.283 | 676H+2X5 Hamilton, New Zealand | 1352 | 13.6 |
| Tu-Scha-9 | 22.01 | 48.53 | 9.05 | G3J2+222 Tübingen, Germany | 691.4 | 9.4 |
| Si-0 | 23.08 | 50.8738 | 8.02341 | V2CC+222 Siegen, Germany | 758.3 | 8.3 |
| Blh-1 | 23.56 | 48 | 19 | 2222+222 Bernecebaráti, Hungary | 711.7 | 8.8 |
| Kro-0 | 23.74 | 50.0742 | 8.96617 | 3X99+XXX Hainburg, Germany | 643.2 | 9.5 |
| Kz-9 | 24.3 | 49.5 | 73.1 | G32X+2XX Novo-Troitskoye, Kazakhstan | 352 | 3.1 |
| Nw-4 | 24.78 | 50.5 | 8.5 | GG22+22X Hüttenberg, Germany | 692.6 | 8.9 |
| Zdr-1 | 25.22 | 49.3853 | 16.2544 | 97Q2+X2X Vratislávka, Křižanov, Czechia | 587 | 7.7 |
| NFA-8 | 25.44 | 51.4083 | -0.6383 | C956+X2R Ascot, United Kingdom | 757 | 9.9 |
| Pf-0 | 25.54 | 48.5479 | 9.11033 | H426+222 Tübingen, Germany | 691.4 | 9.4 |
| Zu-1 | 25.77 | 47.3667 | 8.55 | 9HC2+222 Zürich, Switzerland | 1173 | 8.2 |
| Se-0 | 26 | 38.3333 | -3.53333 | 8FH9+XXX Santa Elena, Spain | 484 | 15.8 |
| Timpo-1 | 26.18 | 39.27 | 16.27 | 779C+X2R Cosenza, Province of Cosenza, Italy | 929 | 13.7 |
| Gr-1 | 26.36 | 47 | 15.5 | 2G22+225 Hausmannstätten, Austria | 998.6 | 7.7 |
| Baa-1 | 27.2 | 51.3333 | 6.1 | 83HX+XXX Baarlo, Netherlands | 756 | 10.2 |
| Cdm-0 | 27.59 | 39.73 | -5.74 | P7H5+XXX Casas de Miravete, Spain | 392 | 15.4 |
| Sij-2 | 27.78 | 41.45 | 70.05 | C3X2+X2R Nurekata, Uzbekistan | 777.5 | 4.8 |
| Omo-2-3 | 27.81 | 56.1509 | 15.7735 | 5Q2C+222 Möcklö, Sweden | 570.7 | 7.2 |
| Kl-5 | 28.31 | 50.95 | 6.9666 | WXX9+XXX Cologne, Germany | 836.5 | 9.9 |
| Fr-4 | 28.35 | 50.1102 | 8.6822 | 4M6J+222 Frankfurt am Main, Germany | 643.2 | 9.5 |
| Wt-5 | 28.37 | 52.3 | 9.3 | 8822+222 Apelern, Germany | 738 | 9 |
| Uod-1 | 28.4 | 48.3 | 14.45 | 8C2X+2X5 Sankt Georgen an der Gusen, Austria | 946.4 | 8.3 |

| | | | | | | |
|---|---|---|---|---|---|---|
| Bd-0 | 28.43 | 52.4584 | 13.287 | F76Q+2X5 Berlin, Germany | 572 | 9.4 |
| Fi-1 | 29.42 | 50.5 | 8.0167 | G22C+22R Dornburg, Germany | 758.3 | 8.3 |
| Chat-1 | 29.44 | 48.0717 | 1.33867 | 389R+X2R Châteaudun, France | 626.6 | 10.6 |
| Sij-1 | 29.84 | 41.45 | 70.05 | C3X2+X2R Nurekata, Uzbekistan | 777.5 | 4.8 |
| Lip-0 | 29.85 | 50 | 19.3 | 2822+222 Poręba Wielka, Poland | 747.6 | 8.3 |
| Voeran-1 | 29.91 | 46.36 | 11.23 | 966H+2X5 Tramin an der Weinstraße, Autonomous | 779 | 7.8 |
| Wei-0 | 29.99 | 47.25 | 8.26 | 7725+2XX Hitzkirch, Switzerland | 1113 | 9 |
| Rou-0 | 30.06 | 49.4424 | 1.09849 | C3QX+XXX Rouen, France | 686.8 | 10.6 |
| An-1 | 30.07 | 51.2167 | 4.4 | 6C92+X2R Antwerp, Belgium | 780.8 | 10.4 |
| Da-0 | 31.08 | 49.8724 | 8.65081 | VMC2+222 Darmstadt, Germany | 655.5 | 9.6 |
| Aitba-2 | 31.28 | 31.48 | -7.45 | FHH2+X2R أوكوك, Morocco | 423 | 14.1 |
| Ct-1 | 31.66 | 37.3 | 15 | 72X2+X2R Lentini, Free municipal consortium of | 522.5 | 17.5 |
| Ba-1 | 31.69 | 56.5459 | -4.79821 | H52X+2X5 Bridge of Orchy, United Kingdom | 2574 | 5.9 |
| Pa-1 | 31.71 | 38.07 | 13.22 | 3699+XXX Giardinello, Metropolitan City of Palermo, | 586.6 | 16.8 |
| Ga-2 | 32 | 50.3 | 8 | 8222+222 Schönborn, Germany | 642.6 | 9.1 |
| Do-0 | 32.34 | 50.7224 | 8.2372 | P69R+X2R Dillenburg, Germany | 758.3 | 8.3 |
| Pt-0 | 32.51 | 53.476 | 10.6065 | FJH6+X2R Büchen, Germany | 636 | 8.9 |
| Ga-0 | 32.69 | 50.3 | 8 | 8222+222 Schönborn, Germany | 642.6 | 9.1 |
| Vezzano-2 | 32.88 | 46.6297 | 10.817 | JRH9+XXX Schlanders, Autonomous Province of | 1140 | 0.6 |
| Rovero-1 | 33.02 | 46.2543 | 11.167 | 7529+2XX Roveré della Luna, Autonomous Province of | 778.6 | 7.8 |
| Pu2-23 | 33.02 | 49.42 | 16.36 | C9C6+222 Doubravník, Czechia | 587 | 7.7 |
| Petro-1 | 33.33 | 44.34 | 21.46 | 8FR6+222 Bistrica, Serbia | 721.6 | 11.7 |
| Pi-2 | 34.3 | 47.04 | 10.51 | 2GQ5+XXX Serfaus, Austria | 1027 | 3.7 |
| Ey1.5-2 | 34.52 | 48.43 | 8.77 | CQJC+222 Starzach, Germany | 940.2 | 8.1 |
| Bolin-1 | 34.9 | 44.46 | 25.74 | FP6R+222 Bolintin-Vale, Romania | 607.1 | 11.4 |
| Pla-0 | 34.98 | 41.5 | 2.25 | G722+22X La Conreria, Spain | 726.6 | 13 |
| Sei-0 | 35.13 | 46.5438 | 11.5614 | GHQ6+X2R Seis am Schlern, Autonomous Province of | 974 | 3.5 |
| Rsch-4 | 35.41 | 56.3 | 34 | 8222+222 Kosterevo, Tver Oblast, Russia | 631.4 | 4.1 |
| Gy-0 | 35.99 | 49 | 2 | 2222+222 Triel-sur-Seine, France | 660 | 10.7 |
| RRS-7 | 36.71 | 41.5609 | -86.4251 | HH6C+222 North Liberty, Indiana, USA | 955 | 9.8 |
| Ob-0 | 36.88 | 50.2 | 8.5833 | 5HXJ+X2R Oberursel, Germany | 643.2 | 9.5 |
| Bch-1 | 37.1 | 49.5166 | 9.3166 | G899+XXX Buchen, Germany | 702.9 | 8.8 |
| Or-0 | 37.34 | 50.3827 | 8.01161 | 92H5+XXX Diez, Germany | 642.6 | 9.1 |
| Gel-1 | 37.43 | 51.0167 | 5.86667 | 2V9C+X2R Sittard, Netherlands | 734.9 | 10.1 |
| Lo-1 | 37.88 | 47.6166 | 7.6666 | JMC9+2X5 Lörrach, Germany | 911 | 8.9 |
| Boot-1 | 38.01 | 54.4 | -3.2667 | CP2H+2X5 Holmrook, United Kingdom | 2460 | 8.2 |
| Sg-1 | 38.05 | 47.6667 | 9.5 | MG92+X2X Friedrichshafen, Germany | 936.7 | 8.5 |
| Slavi-1 | 38.4 | 41.43 | 23.65 | CMJ2+222 Paril, Bulgaria | 627.7 | 11 |
| En-1 | 38.58 | 50 | 8.5 | 2G22+225 Rüsselsheim, Germany | 643.2 | 9.5 |
| Hh-0 | 39 | 54.4175 | 9.88682 | CV9Q+XXX Holtsee, Germany | 817.8 | 8.6 |
| Ge-0 | 39.09 | 46.5 | 6.08 | Lausanne, a city in western Switzerland, located on the | 1121 | 6.7 |
| Sq-8 | 39.14 | 51.4083 | -0.6383 | C956+X2R Ascot, United Kingdom | 757 | 9.9 |
| KNO-18 | 39.41 | 41.2816 | -86.621 | 79JJ+222 Knox, Indiana, USA | 970 | 9.8 |
| Bla-1 | 39.41 | 41.6833 | 2.8 | MRJ2+222 Blanes, Spain | 767.3 | 14.2 |
| Ru3.1-31 | 39.65 | 48.56 | 9.16 | H565+2X5 Pliezhausen, Germany | 691.4 | 9.4 |
| Ge-1 | 40 | 46.5 | 6.08 | Lausanne, a city in western Switzerland, located on the | 1121 | 6.7 |
| Old-2 | 40.27 | 53.1667 | 8.2 | 559X+XXX Oldenburg, Germany | 753.6 | 9.3 |
| Ws-2 | 40.86 | 52.3 | 30 | 8222+222 Babichi, Belarus | 626.8 | 7.1 |
| Altenb-2 | 41.25 | 46.3716 | 11.2376 | 96CR+222 Kaltern an der Weinstraße, Autonomous | 779 | 7.8 |
| Zdr-6 | 41.5 | 49.3853 | 16.2544 | 97Q2+X2X Vratislávka, Křižanov, Czechia | 587 | 7.7 |
| CIBC-5 | 41.61 | 51.4083 | -0.6383 | C956+X2R Ascot, United Kingdom | 757 | 9.9 |
| Wa-1 | 41.89 | 52.3 | 21 | 8222+222 Warsaw, Poland | 527 | 8.1 |
| Wal-HasB-4 | 42.8 | 48.6 | 9.19 | H5XQ+XXX Schlaitdorf, Germany | 691.4 | 9.4 |
| Old-1 | 43 | 53.1667 | 8.2 | 559X+XXX Oldenburg, Germany | 753.6 | 9.3 |
| In-0 | 43.12 | 47.5 | 11.5 | GG22+22X Simmering, Austria | 948 | 7 |
| Nok-3 | 43.3 | 52.24 | 4.45 | 6CRX+2X5 Noordwijk, Netherlands | 805.8 | 9.9 |
| Sapporo-0 | 43.38 | 43.0553 | 141.346 | 3962+222 Sapporo, Hokkaido, Japan | 1256 | 6.9 |
| Ven-1 | 44.68 | 52.0333 | 5.55 | 2HJ2+222 Veenendaal, Netherlands | 777.3 | 9.5 |
| Ob-1 | 45.21 | 50.2 | 8.5833 | 5HXJ+X2R Oberursel, Germany | 643.2 | 9.5 |
| Wt-1 | 45.9 | 52.3 | 9.3 | 8822+222 Apelern, Germany | 738 | 9 |
| Ma-2 | 46.2 | 50.8167 | 8.7667 | RQ9C+X2R Marburg, Germany | 692.6 | 8.9 |
| Sha | 47.15 | 37.29 | 71.3 | 8J9H78R2+22, Sughd Region, north of the city of | 652 | 1.9 |
| Nw-0 | 47.35 | 50.5 | 8.5 | GG22+22X Hüttenberg, Germany | 692.6 | 8.9 |
| La-1 | 47.39 | 52.7333 | 15.2333 | P6HH+XXX Gorzów Wielkopolski, Poland | 547 | 9.1 |
| Np-0 | 48 | 52.6969 | 10.981 | MXXH+XXX Rohrberg, Germany | 639.5 | 8.9 |
| Sorbo | 48.07 | 38.35 | 68.48 | 8FXJ+X2R Lattaband, Tajikistan | 464 | 14.6 |
| Stepn-1 | 48.17 | 54.06 | 60.48 | 3F6J+222 Stepnoe, Chelyabinsk Oblast, Russia | 469 | 2.2 |
| Cnt-1 | 48.21 | 51.3 | 1.1 | 832X+2X5 Canterbury, United Kingdom | 735.4 | 10.3 |
| Est-1 | 48.29 | 58.3 | 25.3 | 8822+222 Supsi, Viljandi County, Estonia | 640.6 | 5.4 |
| Nd-0 | 51.94 | 50 | 10 | 2222+222 Arnstein, Germany | 685 | 8.3 |
| Wl-0 | 52.11 | 47.9299 | 10.8134 | WRJ6+222 Fuchstal, Germany | 1009 | 7.4 |
| Leb-3 | 53.01 | 51.65 | 80.82 | MR29+2X5 Lebyazh'e, Altai Krai, Russia | 332 | 2.6 |

| | | | | | | |
|---|---|---|---|---|---|---|
| NFA-10 | 53.35 | 51.4083 | -0.6383 | C956+X2R Ascot, United Kingdom | 757 | 9.9 |
| Bsch-2 | 53.91 | 50.0167 | 8.6667 | 2M99+XXX Dreieich, Germany | 643 | 9.5 |
| Cen-0 | 55.39 | 49 | 0.5 | 2G22+225 Saint-Aubin-du-Thenney, France | 714.5 | 10.5 |
| Tscha-1 | 55.7 | 47.0748 | 9.9042 | 3W92+X2R Tschagguns, Austria | 1040 | 6.6 |
| Ei-2 | 56.05 | 50.3 | 6.3 | 8822+222 Auw bei Prüm, Germany | 1026 | 8.2 |
| Pa-3 | 57.17 | 38.07 | 13.22 | 3699+XXX Giardinello, Metropolitan City of Palermo, | 586.6 | 16.8 |
| Apost-1 | 57.81 | 39.01 | 16.47 | 2F59+XXX Gimigliano, Province of Catanzaro, Italy | 929.4 | 13.7 |
| Mrk-0 | 58.37 | 49 | 9.3 | 2822+222 Großbottwar, Germany | 673.9 | 9.4 |
| Ciste-2 | 59.09 | 41.62 | 12.87 | JVCC+222 Cori, Province of Latina, Italy | 874.5 | 14.7 |
| Mz-0 | 60.42 | 50.3 | 8.3 | 8822+222 Bad Camberg, Germany | 643 | 9.1 |
| Bs-1 | 61.05 | 47.5 | 7.5 | GG22+22X Leymen, France | 911 | 8.9 |
| Mammo-2 | 61.38 | 38.38 | 16.22 | 96H9+XXX Mammola, Metropolitan City of Reggio | 895.4 | 15.7 |
| Bn-0 | 61.47 | 50.5 | 9.5 | GG22+22X Hosenfeld, Germany | 709 | 8.2 |
| Bor-1 | 61.75 | 49.4013 | 16.2326 | C62H+2X5 Strážek, Czechia | 587 | 7.7 |
| Ove-0 | 62.06 | 53.3422 | 8.42255 | 8CR9+2X5 Ovelgönne, Germany | 753.6 | 9.3 |
| Lag2-2 | 63.32 | 41.8296 | 46.2831 | R7HJ+X2R Lagodekhi, Georgia | 800 | 11.3 |
| Lc-0 | 63.36 | 57 | -4 | 2222+222 Kingussie, United Kingdom | 1150 | 5.8 |
| Borsk-2 | 64.22 | 53.04 | 51.75 | 2QQ2+X2X Novoborskii, Samara Oblast, Russia | 531 | 4.4 |
| Moran-1 | 64.36 | 39.83 | 16.17 | R5H9+XXX Morano Calabro, Province of Cosenza, Italy | 847.1 | 14.3 |
| Mh-0 | 64.54 | 50.95 | 20.5 | WGX2+X2X Kostomłoty Drugie, Poland | 653.3 | 7.6 |
| HR-5 | 64.71 | 51.4083 | -0.6383 | C956+X2R Ascot, United Kingdom | 757 | 9.9 |
| Ka-0 | 65.19 | 47 | 14 | 2222+222 Minigraben, Austria | 1609 | 4.1 |
| Ep-0 | 65.47 | 50.1721 | 8.38912 | 599Q+XXX Kelkheim, Germany | 642.6 | 9.1 |
| Mammo-1 | 66.61 | 38.36 | 16.23 | 966H+2X5 Mammola, Metropolitan City of Reggio | 895.4 | 15.7 |
| Go-0 | 67.16 | 51.5338 | 9.9355 | GWJQ+2X5 Göttingen, Germany | 782.9 | 8.5 |
| Jm-0 | 67.85 | 49 | 15 | 2222+222 Rottal, Austria | 697 | 7 |
| Kelsterbach-4 | 68.15 | 50.0667 | 8.5333 | 3G9J+X2R Kelsterbach, Germany | 643.2 | 9.5 |
| Is-0 | 68.39 | 50.5 | 7.5 | GG22+22X Rengsdorf, Germany | 757.9 | 8.9 |
| Db-0 | 68.53 | 50.3055 | 8.324 | 8869+2X5 Bad Camberg, Germany | 642.6 | 9.1 |
| Fei-0 | 69.58 | 40.92 | -8.54 | WF96+X2R Santa Maria da Feira, Portugal | 1088 | 15.3 |
| Co-3 | 70.08 | 40.12 | -8.25 | 4QC2+225 Lousã, Portugal | 1037.2 | 15 |
| No-0 | 70.28 | 51.0581 | 13.2995 | 3862+222 Nossen, Germany | 589.6 | 9 |
| Je-0 | 71.59 | 50.927 | 11.587 | WHJR+222 Jena, Germany | 643.7 | 7.8 |
| Kil-0 | 71.74 | 55.6395 | -5.66364 | J8QR+X2R Tarbert, United Kingdom | 1571 | 8.3 |
| Lecho-1 | 71.93 | 41.43 | 23.5 | CGJ2+225 Petrovo, Bulgaria | 628 | 11 |
| Lz-0 | 71.96 | 46 | 3.3 | 2822+222 Saint-Clément-de-Régnat, France | 733 | 10.5 |
| Col-0 | 72.13 | 38.3 | -92.3 | 8P22+222 St Elizabeth, Missouri, USA | 1016 | 13.4 |
| Hs-0 | 73.71 | 52.24 | 9.44 | 6CRQ+2X5 Bad Münder, Germany | 738 | 9 |
| Com-1 | 73.88 | 49.416 | 2.823 | CRC9+2X5 Compiègne, France | 639.4 | 10.7 |
| Dra-1 | 74.22 | 49.4167 | 16.2667 | C7CC+222 Drahonín, Doubravník, Czechia | 587 | 7.7 |
| Bor-4 | 74.6 | 49.4013 | 16.2326 | C62H+2X5 Strážek, Czechia | 587 | 7.7 |
| Lago-1 | 74.71 | 39.18 | 16.26 | 57J5+2X5 Malito, Province of Cosenza, Italy | 929 | 13.7 |
| Hi-0 | 77.74 | 52 | 5 | 2222+222 Benschop, Netherlands | 792 | 9.7 |
| Gu-0 | 78.4 | 50.3 | 8 | 8222+222 Schönborn, Germany | 643 | 9.1 |
| Ll-0 | 78.61 | 41.59 | 2.49 | HFRR+222 Sant Vicenç de Montalt, Spain | 727 | 13 |
| Nok-1 | 80.15 | 52.24 | 4.45 | 6CRX+2X5 Noordwijk, Netherlands | 806 | 9.9 |
| Hn-0 | 81.49 | 51.3472 | 8.28844 | 87XQ+XXX Meschede, Germany | 841 | 7.9 |
| Mer-6 | 81.57 | 38.92 | -6.34 | WMC5+2X5 Mérida, Spain | 503 | 16.3 |
| Nie1-2 | 83.36 | 48.52 | 8.8 | GR92+X2R Bondorf, Germany | 806 | 9 |
| Chi-1 | 84.09 | 53.7502 | 34.7361 | QP2R+22R Zhizdra, Kaluga Oblast, Russia | 623 | 5 |
| Di-1 | 85.92 | 47 | 5 | 2222+222 Corberon, France | 831 | 10.9 |
| Kr-0 | 86.24 | 51.3317 | 6.55934 | 8HH6+X2R Krefeld, Germany | 818 | 10.4 |
| Dra-2 | 87.49 | 49.4167 | 16.2667 | C7CC+222 Drahonín, Doubravník, Czechia | 587 | 7.7 |
| Co-2 | 88.15 | 40.12 | -8.25 | 4QC2+225 Lousã, Portugal | 1037 | 15 |
| Na-1 | 88.88 | 47.5 | 1.5 | GG22+22X Cour-Cheverny, France | 622 | 11.2 |
| Li-2:1 | 89.29 | 50.3833 | 8.0666 | 93H9+XXX Limburg, Germany | 643 | 9.1 |
| Gie-0 | 89.53 | 50.584 | 8.67825 | HMHJ+X2R Giessen, Germany | 693 | 8.9 |
| Li-6 | 90.6 | 50.3833 | 8.0666 | 93H9+XXX Limburg, Germany | 643 | 9.1 |
| Kondara | 92.05 | 38.48 | 68.49 | FFHR+X2R Hisor, Tajikistan | 464 | 14.6 |
| Ca-0 | 93.69 | 50.2981 | 8.26607 | 872C+222 Bad Camberg, Germany | 643 | 9.1 |
| Jl-3 | 94.15 | 49.2 | 16.6166 | 5JXC+X2R Brno, Czechia | 586 | 8.1 |
| Da(1)-12 | 94.62 | | | Unknown, Czechia | 691 | 8.1 |
| El-0 | 95.46 | 51.5105 | 9.68253 | GM5J+X2R Niemetal, Germany | 783 | 8.5 |
| Lp2-2 | 99.03 | 49.38 | 16.81 | 9RH6+X2R Lipovec, Czechia | 586 | 8.1 |
| Mnz-0 | 99.93 | 50.001 | 8.26664 | 272C+222 Mainz, Germany | 643 | 9.1 |
| Ha-0 | 100.37 | 52.3721 | 9.73569 | 9PCR+222 Hanover, Germany | 721 | 9 |
| Dr-0 | 102.3 | 51.051 | 13.7336 | 3P2H+2X5 Dresden, Germany | 624 | 9.2 |
| Kin-0 | 102.71 | 44.46 | -85.37 | FJ6J+222 Manton, Michigan, USA | 789 | 6.5 |
| Hl-3 | 117.86 | 52.1444 | 9.37827 | 49QH+XXX Hamelin, Germany | 738 | 9 |
| Eil-0 | 150.95 | 51.4599 | 12.6327 | FJ6H+2X5 Eilenburg, Germany | 564 | 8.8 |

### Table S2. Network interacting partners of genes identified by a GWAS of drought tolerance in *Arabidopsis thaliana*

| AGI Code | Gene Symbol | (-)log$_{10}$ GWAS *P*-value | Short description | Sub network | Average Shortest Path Length | Betweenness Centrality | Closeness Centrality | Clustering Coefficient | Degree | Eccentricity | Neighborhood Connectivity | Number Of Directed Edges | Number Of Undirected Edges | Omics Visualizer:: Connected rows | Partner Of MultiEdged Node Pairs | Radiality | Stress | Topological Coefficient | Enriched GO Biological Process |
|---|---|---|---|---|---|---|---|---|---|---|---|---|---|---|---|---|---|---|---|
| AT1G14460 | **AT1G14460** | 4.33 | AAA-type atpase family protein | (a) | 2.5 | 8.30E-05 | 0.4 | 0.8 | 4 | 3 | 11.8 | 0 | 4 | 5 | 0 | 1 | 2 | 0.7 | Cellular response to stress, DNA metabolic process, DNA repair, DNA replication, Nucleobase-containing compound metabolic process |
| AT1G01550 | *BPS1* | | Bypass 1 | (a) | 1.5 | 5.76E-03 | 0.7 | 0.7 | 22 | 3 | 26.2 | 0 | 22 | 0 | 0 | 1 | 138 | 0.6 | |
| AT2G46080 | *BPS2* | | Bypass 2 | (a) | 1.5 | 5.76E-03 | 0.7 | 0.7 | 22 | 3 | 26.2 | 0 | 22 | 0 | 0 | 1 | 138 | 0.6 | |
| AT4G01360 | *BPS3* | | Bypass 3 | (a) | 1.5 | 5.76E-03 | 0.7 | 0.7 | 22 | 3 | 26.2 | 0 | 22 | 1 | 0 | 1 | 138 | 0.6 | Cellular response to stress, Cell cycle, Cellular response to stress, DNA metabolic process, DNA repair, DNA replication, Nucleobase-containing |
| AT1G07270 | *CDC6B* | | Cell division control protein 6 homolog b | (a) | 1.5 | 6.98E-03 | 0.7 | 0.8 | 25 | 3 | 24.8 | 0 | 25 | 0 | 0 | 1 | 144 | 0.6 | |
| AT2G31270 | *CDT1A* | | Homolog of yeast cdt1 a | (a) | 1.4 | 9.56E-03 | 0.7 | 0.7 | 24 | 2 | 24.9 | 0 | 24 | 0 | 0 | 1 | 248 | 0.6 | |
| AT3G54710 | *CDT1B* | | Homolog of yeast cdt1 b homolog of yeast cdt1 b | (a) | 1.4 | 9.56E-03 | 0.7 | 0.7 | 24 | 2 | 24.9 | 0 | 24 | 0 | 0 | 1 | 248 | 0.6 | |
| AT4G16970 | *Dl4515c* | | Protein kinase superfamily protein | (a) | 1.9 | 3.32E-05 | 0.5 | 1 | 7 | 3 | 36 | 0 | 7 | 4 | 0 | 1 | 2 | 0.9 | Cellular response to stress, DNA metabolic process, DNA repair, Nucleobase-containing compound |
| AT3G46940 | ***DUT1*** | 3.43 | dUTP-pyrophosphatase-like 1 | (a) | 1.9 | 8.30E-05 | 0.5 | 0.9 | 5 | 2 | 23.6 | 0 | 5 | 4 | 0 | 1 | 2 | 0.6 | Cellular response to stress, DNA metabolic process, DNA repair, Nucleobase-containing compound |
| AT2G40550 | *ETG1* | | E2F target gene 1 | (a) | 1.9 | 3.32E-05 | 0.5 | 1 | 8 | 3 | 34.3 | 0 | 8 | 6 | 0 | 1 | 2 | 0.9 | Cell cycle, Cellular response to stress, DNA metabolic process, DNA repair, DNA replication, Nucleobase-containing compound metabolic process |
| AT3G02820 | *F13E7.24* | | Zinc knuckle (CCHC-type) family protein | (a) | 1.6 | 8.25E-04 | 0.6 | 0.9 | 18 | 3 | 26.4 | 0 | 18 | 5 | 0 | 1 | 32 | 0.6 | Cell cycle |
| AT3G42660 | *F4JF14_ARATH* | | Transducin family protein / WD-40 repeat family protein | (a) | 1.5 | 1.51E-03 | 0.6 | 0.9 | 20 | 3 | 26.3 | 0 | 20 | 0 | 0 | 1 | 50 | 0.6 | |
| AT3G61500 | *F4JFC6_ARATH* | | BPS1-like protein | (a) | 1.5 | 5.76E-03 | 0.7 | 0.7 | 22 | 3 | 26.2 | 0 | 22 | 0 | 0 | 1 | 138 | 0.6 | |
| AT3G12530 | *GINS2* | | GINS complex subunit 2 | (a) | 1.6 | 9.62E-04 | 0.6 | 0.9 | 19 | 3 | 26.4 | 0 | 19 | 5 | 0 | 1 | 36 | 0.6 | Cellular response to stress |
| AT5G17510 | *K3M16.80* | | Mediator of RNA polymerase II transcription subunit-like protein | (a) | 1.9 | 3.32E-05 | 0.5 | 1 | 7 | 3 | 36 | 0 | 7 | 0 | 0 | 1 | 2 | 0.9 | |
| AT1G44900 | *MCM2* | | Minichromosome maintenance 2 | (a) | 1.4 | 1.11E-02 | 0.7 | 0.7 | 24 | 2 | 25 | 0 | 24 | 0 | 0 | 1 | 240 | 0.6 | |
| AT5G46280 | *MCM3* | | Minichromosome maintenance 3 | (a) | 1.1 | 6.52E-02 | 0.9 | 0.5 | 38 | 2 | 20.7 | 0 | 38 | 7 | 0 | 1 | 904 | 0.5 | Cell cycle, Cellular response to stress, DNA metabolic process, DNA repair, DNA replication, Nucleobase-containing compound metabolic process |
| AT2G16440 | *MCM4* | | Minichromosome maintenance 4 | (a) | 1.1 | 7.28E-02 | 0.9 | 0.5 | 39 | 2 | 20.4 | 0 | 39 | 0 | 0 | 1 | 1028 | 0.5 | |
| AT2G07690 | *MCM5* | | Minichromosome maintenance 5 | (a) | 1.1 | 6.09E-02 | 0.9 | 0.5 | 38 | 2 | 20.8 | 0 | 38 | 0 | 0 | 1 | 952 | 0.5 | |
| AT5G44635 | *MCM6* | | Minichromosome maintenance 6 | (a) | 1.2 | 4.57E-02 | 0.9 | 0.5 | 35 | 2 | 20.8 | 0 | 35 | 0 | 0 | 1 | 792 | 0.5 | |
| AT4G02060 | ***MCM7*** | 3.36 | Minichromosome maintenance 7 | (a) | 1 | 9.09E-02 | 1 | 0.5 | 40 | 2 | 20 | 0 | 40 | 7 | 0 | 1 | 1098 | 0.5 | Cell cycle |
| AT3G09660 | *MCM8* | | Minichromosome maintenance 8 | (a) | 1.2 | 4.57E-02 | 0.9 | 0.5 | 35 | 2 | 20.8 | 0 | 35 | 0 | 0 | 1 | 792 | 0.5 | |
| AT2G14050 | *MCM9* | | Minichromosome maintenance 9 | (a) | 1.5 | 5.76E-03 | 0.7 | 0.7 | 22 | 3 | 26.2 | 0 | 22 | 5 | 0 | 1 | 138 | 0.6 | Cell cycle, Cellular response to stress, DNA metabolic process, DNA repair, Nucleobase-containing |
| AT2G20980 | *MCM10* | | Minichromosome maintenance 10 | (a) | 1.4 | 1.41E-02 | 0.7 | 0.7 | 27 | 3 | 23.9 | 0 | 27 | 3 | 0 | 1 | 238 | 0.6 | DNA metabolic process, DNA replication, Nucleobase-containing compound metabolic process |
| AT5G45430 | *MFC19.10* | | Protein kinase superfamily protein | (a) | 1.7 | 2.43E-04 | 0.6 | 0.9 | 14 | 3 | 29.6 | 0 | 14 | 0 | 0 | 1 | 10 | 0.7 | |
| AT4G13020 | *MHK* | | Protein kinase superfamily protein | (a) | 1.7 | 2.43E-04 | 0.6 | 0.9 | 14 | 3 | 29.6 | 0 | 14 | 0 | 0 | 1 | 10 | 0.7 | |
| AT4G02070 | ***MSH6*** | 3.36 | MUTS homolog 6 | (a) | 2.3 | 8.30E-05 | 0.4 | 0.7 | 3 | 3 | 17 | 0 | 3 | 5 | 0 | 1 | 2 | 0.6 | Cellular response to stress, DNA metabolic process, DNA repair, Nucleobase-containing compound |
| AT2G37560 | *ORC2* | | Origin recognition complex second largest subunit 2 | (a) | 1.5 | 8.42E-03 | 0.7 | 0.7 | 25 | 3 | 24.3 | 0 | 25 | 3 | 0 | 1 | 156 | 0.6 | DNA metabolic process, DNA replication, Nucleobase-containing compound metabolic process |
| AT5G16690 | *ORC3* | | Origin recognition complex subunit 3 | (a) | 1.5 | 5.02E-03 | 0.7 | 0.8 | 22 | 3 | 24.2 | 0 | 22 | 3 | 0 | 1 | 112 | 0.6 | DNA metabolic process, DNA replication, Nucleobase-containing compound metabolic process |
| AT2G01120 | *ORC4* | | Origin recognition complex subunit 4 | (a) | 1.5 | 5.44E-03 | 0.7 | 0.8 | 23 | 3 | 25.2 | 0 | 23 | 3 | 0 | 1 | 120 | 0.6 | DNA metabolic process, DNA replication, Nucleobase-containing compound metabolic process |
| AT4G29910 | *ORC5* | | Origin recognition complex protein 5 | (a) | 1.5 | 5.00E-03 | 0.7 | 0.8 | 23 | 3 | 25 | 0 | 23 | 0 | 0 | 1 | 112 | 0.6 | |
| AT1G26840 | *ORC6* | | Origin recognition complex protein 6 | (a) | 1.9 | 5.05E-05 | 0.5 | 1 | 8 | 3 | 27.5 | 0 | 8 | 3 | 0 | 1 | 2 | 0.7 | DNA metabolic process, DNA replication, Nucleobase-containing compound metabolic process |
| AT1G07370 | *PCNA* | | Proliferating cellular nuclear antigen 1 | (a) | 1.6 | 3.17E-02 | 0.6 | 0.5 | 15 | 2 | 23.3 | 0 | 15 | 8 | 0 | 1 | 714 | 0.6 | Cellular response to stress, DNA metabolic process, DNA repair, DNA replication, Nucleobase-containing compound metabolic process |
| AT2G29570 | *PCNA2* | | Proliferating cell nuclear antigen 2 | (a) | 1.7 | 2.81E-02 | 0.6 | 0.5 | 14 | 2 | 23.2 | 0 | 14 | 8 | 0 | 1 | 662 | 0.6 | Cellular response to stress, DNA metabolic process, DNA repair, DNA replication, Nucleobase-containing compound metabolic process |
| AT5G67100 | *POLA* | | DNA polymerase alpha catalytic subunit | (a) | 1.5 | 2.86E-02 | 0.7 | 0.7 | 22 | 2 | 24.1 | 0 | 22 | 5 | 0 | 1 | 422 | 0.6 | Cell cycle, DNA metabolic process, DNA replication, Nucleobase-containing compound metabolic process |
| AT1G80190 | *PSF1* | | Partner of SLD five 1 | (a) | 1.6 | 9.62E-04 | 0.6 | 0.9 | 19 | 3 | 26.4 | 0 | 19 | 4 | 0 | 1 | 36 | 0.6 | Cell cycle, DNA metabolic process, DNA replication, Nucleobase-containing compound metabolic process |
| AT5G27740 | *RFC3* | | Replication factor C subunit 3 | (a) | 1.7 | 1.16E-02 | 0.6 | 0.7 | 11 | 2 | 24.5 | 0 | 11 | 8 | 0 | 1 | 334 | 0.6 | |
| AT1G77470 | *RFC5* | | Replication factor C subunit 5 | (a) | 1.8 | 4.80E-03 | 0.5 | 0.8 | 7 | 2 | 23 | 0 | 7 | 8 | 0 | 1 | 162 | 0.5 | Cellular response to stress, DNA metabolic process, DNA repair, DNA replication, Nucleobase-containing compound metabolic process, Nucleotide excision |

| | | | | | | | | | | | | | | | | | | | |
|---|---|---|---|---|---|---|---|---|---|---|---|---|---|---|---|---|---|---|---|
| AT5G49010 | *SLD5* | | Synthetic lethality with DPB11-1 5 | (a) | 1.6 | 9.62E-04 | 0.6 | 0.9 | 19 | 3 | 26.4 | 0 | 19 | 5 | 0 | 1 | 36 | 0.6 | Cellular response to stress, DNA metabolic process, DNA repair, DNA replication, Nucleobase-containing compound metabolic process |
| AT4G19110 | *T18B16.80* | | Protein kinase superfamily protein | (a) | 1.7 | 2.43E-04 | 0.6 | 0.9 | 14 | 3 | 29.6 | 0 | 14 | 0 | 0 | 1 | 10 | 0.7 | |
| AT3G03460 | *T21P5.12* | | Mediator of RNA polymerase II transcription subunit-like protein | (a) | 1.9 | 3.32E-05 | 0.5 | 1 | 7 | 3 | 36 | 0 | 7 | 0 | 0 | 1 | 2 | 0.9 | |
| AT5G12130 | *TERC* | | Pigment defective 149 | (a) | 1.5 | 5.44E-03 | 0.7 | 0.8 | 23 | 3 | 25.2 | 0 | 23 | 0 | 0 | 1 | 120 | 0.6 | |
| AT1G19080 | *TTN10* | | Titan 10 | (a) | 1.6 | 8.25E-04 | 0.6 | 0.9 | 18 | 3 | 26.4 | 0 | 18 | 4 | 0 | 1 | 32 | 0.6 | Cell cycle, DNA metabolic process, DNA replication, Nucleobase-containing compound metabolic process |
| AT1G67620 | *F12B7.17* | | Lojap-related protein | (b) | 1.2 | 3.33E-01 | 0.9 | 0.6 | 5 | 2 | 4 | 0 | 5 | 0 | 0 | 1 | 14 | 0.76 | |
| AT5G39800 | ***ML41Z*** | 3.01 | Mitochondrial ribosomal protein L41Z | (b) | 1.3 | 0 | 0.8 | 1 | 4 | 2 | 5 | 0 | 4 | 0 | 0 | 0.9 | 0 | 0.8 | |
| AT5G55810 | ***NMNAT*** | 3.16 | Nicotinate/nicotinamide mononucleotide adenyltransferase | (b) | 2 | 0 | 0.5 | 0 | 1 | 3 | 5 | 0 | 1 | 1 | 0 | 0.8 | 0 | 0 | Nucleobase-containing compound metabolic process |
| AT4G15950 | ***NRPD4*** | 3.4 | Nuclear RNA polymerase D4 | (b) | 1.7 | 0 | 0.6 | 1 | 3 | 3 | 5 | 0 | 3 | 0 | 0 | 0.9 | 0 | 1 | |
| AT5G18790 | *F17K4.40* | | Ribosomal protein L33 family protein | (b) | 1.2 | 6.67E-02 | 0.9 | 0.8 | 5 | 2 | 4.4 | 0 | 5 | 0 | 0 | 1 | 6 | 0.7 | |
| ATCG00760 | *rpl36* | | Ribosomal protein L36 | (b) | 1.2 | 6.67E-02 | 0.9 | 0.8 | 5 | 2 | 4.4 | 0 | 5 | 0 | 0 | 1 | 6 | 0.7 | |
| AT5G53070 | *MNB8.13* | | Ribosomal protein L9/RNAse H1 | (b) | 1.2 | 6.67E-02 | 0.9 | 0.8 | 5 | 2 | 4.4 | 0 | 5 | 0 | 0 | 1 | 6 | 0.7 | |
| AT1G68725 | *AGP19* | | Arabinogalactan protein 19 | (c) | 1.9 | 0 | 0.5 | 0 | 1 | 2 | 7 | 0 | 1 | 0 | 0 | 0.9 | 0 | 0 | |
| AT5G28050 | *GSDA* | | Guanosine deaminase | (c) | 1.9 | 0 | 0.5 | 0 | 1 | 2 | 7 | 0 | 1 | 3 | 0 | 0.9 | 0 | 0 | Nucleobase-containing compound metabolic process, tRNA modification, tRNA wobble adenosine to inosine |
| AT4G20960 | *PYRD-2* | | Pyrimidine deaminase | (c) | 1.9 | 0 | 0.5 | 0 | 1 | 2 | 7 | 0 | 1 | 0 | 0 | 0.9 | 0 | 0 | |
| AT3G47390 | *PYRR* | | Photosensitive 1 | (c) | 1.9 | 0 | 0.5 | 0 | 1 | 2 | 7 | 0 | 1 | 1 | 0 | 0.9 | 0 | 0 | Nucleobase-containing compound metabolic process |
| AT3G05300 | *T12H1.27* | | Cytidine/deoxycytidylate deaminase family protein | (c) | 1.9 | 0 | 0.5 | 0 | 1 | 2 | 7 | 0 | 1 | 3 | 0 | 0.9 | 0 | 0 | Nucleobase-containing compound metabolic process, tRNA modification, tRNA wobble adenosine to inosine editing, Cell cycle, Cellular response to stress, DNA metabolic process, DNA repair, DNA replication, Nucleobase-containing compound metabolic process |
| AT1G48175 | *TAD2* | | tRNA arginine adenosine deaminase 2 | (c) | 1.9 | 0 | 0.5 | 0 | 1 | 2 | 7 | 0 | 1 | 3 | 0 | 0.9 | 0 | 0 | Nucleobase-containing compound metabolic process, tRNA wobble adenosine to inosine editing, tRNA |
| AT5G24670 | ***TAD3*** | 3.41 | tRNA arginine adenosine deaminase 3 | (c) | 1 | 1 | 1 | 0 | 7 | 1 | 1 | 0 | 7 | 2 | 0 | 1 | 42 | 0 | Nucleobase-containing compound metabolic process, tRNA modification |
| AT1G68720 | *TADA* | | tRNA arginine adenosine deaminase | (c) | 1.9 | 0 | 0.5 | 0 | 1 | 2 | 7 | 0 | 1 | 3 | 0 | 0.9 | 0 | 0 | Nucleobase-containing compound metabolic process, tRNA wobble adenosine to inosine editing, tRNA |
| AT5G49570 | *PNG1* | | Peptide-N-glycanase 1 | (d) | 1.5 | 0 | 0.7 | 0 | 1 | 2 | 2 | 0 | 1 | 0 | 0 | 0.8 | 0 | 0 | |
| AT3G02540 | ***RAD23-3*** | 3.41 | Radiation sensitive 23-3 | (d) | 1 | 1 | 1 | 0 | 2 | 1 | 1 | 0 | 2 | 5 | 0 | 1 | 2 | 0 | Cellular response to stress |
| AT5G63220 | *MDC12.19* | | Golgi-to-ER traffic-like protein | (d) | 1.5 | 0 | 0.7 | 0 | 1 | 2 | 2 | 0 | 1 | 0 | 0 | 0.8 | 0 | 0 | |
| AT4G27010 | **EMB2788** | 3.7 | Embryo defective 2788 | (e) | 1.5 | 0 | 0.7 | 0 | 1 | 2 | 2 | 0 | 1 | 1 | 0 | 0.8 | 0 | 0 | Nucleobase-containing compound metabolic process |
| AT1G14300 | **AT1G14300** | 3.3 | ARM repeat superfamily protein | (e) | 1.5 | 0 | 0.7 | 0 | 1 | 2 | 2 | 0 | 1 | 0 | 0 | 0.8 | 0 | 0 | |
| AT2G21440 | *F3K23.20* | | RNA-binding (RRM/RBD/RNP motifs) family protein | (e) | 1 | 1 | 1 | 0 | 2 | 1 | 1 | 0 | 2 | 0 | 0 | 1 | 2 | 0 | |
| AT2G39090 | *APC7* | | Anaphase-promoting complex 7 | (f) | 1.3 | 0 | 0.8 | 1 | 2 | 2 | 3 | 0 | 2 | 1 | 0 | 0.9 | 0 | 1 | Cell cycle |
| AT3G05870 | *APC11* | | Anaphase-promoting complex/cyclosome 11 | (f) | 1.3 | 0 | 0.8 | 1 | 2 | 2 | 3 | 0 | 2 | 1 | 0 | 0.9 | 0 | 1 | Cell cycle |
| AT5G51050 | *APC2* | | ATP/phosphate carrier 2 | (f) | 1 | 1.67E-01 | 1 | 0.7 | 3 | 1 | 2.3 | 0 | 3 | 2 | 0 | 1 | 2 | 0.8 | Cell cycle, Protein K11-linked ubiquitination |
| AT1G06590 | ***APC5*** | 3.14 | Anaphase-promoting complex subunit 5 | (f) | 1 | 1.67E-01 | 1 | 0.7 | 3 | 1 | 2.3 | 0 | 3 | 2 | 0 | 1 | 2 | 0.8 | Cell cycle, Protein K11-linked ubiquitination |
| AT5G53000 | *TAP46* | | 2A phosphatase associated protein of 46 kD | (g) | 1.5 | 0 | 0.7 | 0 | 1 | 2 | 2 | 0 | 1 | 0 | 0 | 0.8 | 0 | 0 | |
| AT3G02530 | ***CCT6-2*** | 3.61 | Chaperonin containing t-complex polypeptide-1 subunit 6-2 | (g) | 1 | 1 | 1 | 0 | 2 | 1 | 1 | 0 | 2 | 0 | 0 | 1 | 2 | 0 | |
| AT5G14240 | *F18O22.30* | | Thioredoxin superfamily protein | (g) | 1.5 | 0 | 0.7 | 0 | 1 | 2 | 2 | 0 | 1 | 0 | 0 | 0.8 | 0 | 0 | |
| AT5G21140 | *EMB1379* | | Embryo defective 1379 | (h) | 1 | 0 | 1 | 1 | 2 | 1 | 2 | 0 | 2 | 4 | 0 | 1 | 0 | 1 | Cellular response to stress, DNA metabolic process, DNA repair, Nucleobase-containing compound |
| AT3G15150 | *MMS21* | | High ploidy 2 | (h) | 1 | 0 | 1 | 1 | 2 | 1 | 2 | 0 | 2 | 5 | 0 | 1 | 0 | 1 | Cell cycle, Cellular response to stress, DNA metabolic process, DNA repair, Nucleobase-containing |
| AT1G51130 | ***NSE4A*** | 3.25 | Non-SMC element 4A | (h) | 1 | 0 | 1 | 1 | 2 | 1 | 2 | 0 | 2 | 4 | 0 | 1 | 0 | 1 | Cellular response to stress, DNA metabolic process, DNA repair, Nucleobase-containing compound |

**Table S3. Oligonucleotide sequences used in the study**

| AGI code | Primer name | Sequence (5'--> 3') |
|---|---|---|
| **Real time q-PCR** | | |
| AT3G52590 | *UBQ1* _RT_Fw | TCGTAAGTACAATCAGGATAAGATG |
| | *UBQ1* _RT_Rv | CACTGAAACAAGAAAAACAAACCCT |
| AT1G06690 | AT1G06690_RT_Fw | CCCATCCCAGGAGCCAAGAA |
| | AT1G06690_RT_Rv | GGGACCGTAGCTCACTCACT |
| AT1G14310 | AT1G14310_RT_Fw | GTGGGGAGAAGATGTGCAGAC |
| | AT1G14310_RT_Rv | TGATTGAGTAAGTGTGTACAGGGTC |
| AT1G14470 | AT1G14470_RT_Fw | TGAGCCTGACCGTGTGACTT |
| | AT1G14470_RT_Rv | AGCATAATGATCCGCCAGTGG |
| AT3G46930 | AT3G46930_RT_Fw | TCTTTTACCTAGCCAGATCTGTCCA |
| | AT3G46930_RT_Rv | GCTGCTGTGGTGGTGGAATG |
| AT5G35695 | AT5G35695_RT_Fw | AGCTGGATTAGTCTTAACATGTGCA |
| | AT5G35695_RT_Rv | CGTCACCTTCATTTCCCACTTCAT |
| AT4G26990 | AT4G26990_RT_Fw | AGCCTCCTTTCCCCACGAAT |
| | AT4G26990_RT_Rv | GGTTGGAGAGGCAAAGAATAAAGGT |
| AT3G02510 | AT3G02510_RT_Fw | CCGCCACACTCTTGCCATTG |
| | AT3G02510_RT_Rv | TCTCAAGTTCTTCAGTGACCAGTG |
| AT5G02480 | AT5G02480_RT_Fw | TGAGAGGAGGAGTGGAGGAAGA |
| | AT5G02480_RT_Rv | ACATGGTTGCATCTTTCATCCGA |
| AT4G15950 | AT4G15950_RT_Fw | CGGAATGATGAGCAGCAGACT |
| | AT4G15950_RT_Rv | TGTTGAGAAAGGCAAAGACGCA |
| AT5G52060 | AT5G52060_RT_Fw | AGCAAGAGATTGAAGAGGAGCCT |
| | AT5G52060_RT_Rv | CTCAATGGCGTCACGGGATG |
| AT1G51130 | AT1G51130_RT_Fw | CCGAGGGTTGGTTGTACAAGAA |
| | AT1G51130_RT_Rv | TCTCCGCTTACATCTCCTTCGA |
| AT5G24670 | AT5G24670_RT_Fw | AGGGGAAAAGAGTTTGAACCATCA |
| | AT5G24670_RT_Rv | ACAACAATGCATTAGACCGTGGT |
| **Homozygosity assessment** | | |
| SALK T-DNA left border primer | SALK-LB-1.3 | ATTTTGCCGATTTCGGAAC |
| SAIL T-DNA left border primer | SAIL-LB-1 | TAGCATCTGAATTTCATAACCAATCTCGATACAC |
| Wisc T-DNA left border primer | WiscDsLoxHs LB | TGATCCATGTAGATTTCCCGGACATGAAG |
| AT1G06690 | SALK_058109C_KO_LP | AAGCCGCTTCTCTGAAAAATC |
| | SALK_058109C_KO_RP | TCTTCTTGGCTTCTCCTCCTC |
| | SALK_127525C_KO_LP | GCATAGTTGTGAAGGCGCTAC |
| | SALK_127525C_KO_RP | ACCGTTGTCAAGACTGGTGTC |
| AT1G14470 | SALK_129884C_KO_LP | ATGAACAAAGCTCTTGGGTCC |
| | SALK_129884C_KO_RP | TGCACTCTCCACTGACTCATG |
| AT3G02520 | SALK_084141C_KO_LP | GTCTTAAACCAAAAGCGGACC |
| | SALK_084141C_KO_RP | AGGAGCTGTCATGATGAATGC |
| AT5G02480 | SAIL_583_D08_KO_LP | TCACCAACAATGTCTCCACATAG |
| | SAIL_583_D08_KO_RP | TCAATCCACCACAAGATCTCC |
| AT5G24670 | SALK_009025C_KO_LP | ACCTTAGCCACTAACAACCCC |
| | SALK_009025C_KO_RP | ATGGTCAGGTGACAATGAAGG |
| AT4G15950 | SALK_020157C_KO_LP | ACATGAGGATCATATGCCCTG |
| | SALK_020157C_KO_RP | CTTGAGAGAGGATTTGAGCCC |
| AT4G27000 | SALK_088676C_KO_LP | TCTCGAAGTTGGATCTTGCAG |
| | SALK_088676C_KO_RP | CCATTGTCAAACGGTTACAGC |

| | | |
|---|---|---|
| AT4G02070 | SALK_089638C_KO_LP | GTGTTAGCTTTGTGGGTCGTC |
| | SALK_089638C_KO_RP | TCCAAGCTCATCTAGCACCAC |
| AT4G11670 | SALK_043386C_KO_LP | TTGATGTATCCAAGGGCAGAG |
| | SALK_043386C_KO_RP | GAAGCTTGCCTGCATCTGTAC |
| AT5G55820 | SALK_071684C_KO_LP | ACGAAGATTGGTTTCGACATG |
| | SALK_071684C_KO_RP | TAAACAACGAGAAGCCAAAGG |
| AT5G52060 | SALK_017591C_KO_LP | TCCTTTCCAAGTCCCAAATTC |
| | SALK_017591C_KO_RP | ACTCCCACCACAACAAATACG |
| AT1G41830 | SALK_036979C_KO_LP | TGGTCTTAGTGAATCGGAGATG |
| | SALK_036979C_KO_RP | AACAGTCAATGCGGTTTTACG |
| AT1G51130 | SALK_070555C_KO_LP | GAAGGTGCACTAAGTTGTGGG |
| | SALK_070555C_KO_RP | GTCGAACAAGGAGAGACGTTG |
| AT1G14310 | SALK_129900C_KO_LP | CTGAAGCAAAGAAGAAACCCC |
| | SALK_129900C_KO_RP | GAGCTGGAGAGTGGTCTCATG |
| AT1G14315 | SAIL_837_C08_KO_LP | AGTTGTGACGGTCTTGTTTGC |
| | SAIL_837_C08_KO_RP | AAGATGTGCAGACATTTTCGG |
| AT4G26990 | SALK_200826C_KO_LP | TATTCTCTGGTCCACGACCAC |
| | SALK_200826C_KO_RP | TTACTGTCCTCCAGGTGATGG |
| AT5G39820 | WiscDsLoxHs069_08D_KO_LP | CGAAACAAAGGAAGAAAAACG |
| | WiscDsLoxHs069_08D_KO_RP | AAAGCAAGTTAGGGCAAATCC |

### Table S4. Amino acid haplotypes associated with drought tolerance in 142 *Arabidopsis thaliana* ecotypes

| Haplotype | Representative ecotype | Number of ecotypes | RRL | Rainfall (mm/year) | Amino acid haplotypes | |
|---|---|---|---|---|---|---|
| **AT1G06690.1** | | | | | **127** | **219** |
| H1 | Col-0 | 104 | 41.4 ± 2.38 | 807.9 ± 31.2 | Ile | Tyr |
| H2 | Buckhorn Pass | 14 | 46.1 ± 7.89 | 770.5 ± 46.6 | Ile | Asn |
| H3 | Kondara | 5 | 46.7 ± 14.9 | 512.4 ± 39.7 | Thr | Asn |
| H4 | Lp2-2 | 19 | 57.5 ± 6.4 | 693.5 ± 32.9 | Thr | Tyr |
| | | | | | **polar→ nonpolar** | **polar→ nonpolar** |

| Haplotype | Representative ecotype | Number of ecotypes | RRL | Rainfall (mm/year) | Amino acid haplotypes | | | |
|---|---|---|---|---|---|---|---|---|
| **AT1G14310.1** | | | | | **8** | **20** | **173** | **207** |
| H1 | Zu-1 | 2 | 18.4 ± 7.5 | 1041 ± 132 | Ser | Leu | Thr | Asp |
| H2 | Gr-1 | 38 | 36.3 ± 3.9 | 821.3 ± 32.7 | Ser | Phe | Asn | Asp |
| H3 | Kondara | 6 | 40.7 ± 12 | 575.9 ± 56.7 | Ser | Leu | Asn | Glu |
| H4 | Altenb-2 | 1 | 41.3 | 779 | * | | | |
| H5 | CIBC-5 | 31 | 42.1 ± 3.9 | 829.4 ± 72.2 | Ser | Phe | Thr | Asp |
| H6 | Col-0 | 64 | 51.14 ± 3.4 | 739.2 ± 34.2 | Ser | Leu | Asn | Asp |
| | | | | | **stop gain** | **aliphatic→aromatic** | | |

| Haplotype | Representative ecotype | Number of ecotypes | RRL | Rainfall (mm/year) | Amino acid haplotypes | | | | | |
|---|---|---|---|---|---|---|---|---|---|---|
| **AT1G14315.2** | | | | | **58** | **119** | **136** | **143** | **223** | **296** |
| H1 | Pog-0 | 1 | 19.3 | 1882 | Ser | Asn | Cys | Ser | Trp | Phe |
| H2 | Nok-3 | 17 | 28.96 ± 3.4 | 828.51 ± 108.8 | Leu | Asn | Trp | Ser | Trp | Val |
| H3 | Aitba-2 | 1 | 31.3 | 423 | Leu | Asn | * | | | |
| H4 | Or-0 | 2 | 33.6 ± 3.7 | 710.8 ± 68.2 | Leu | Asn | Trp | Phe | Trp | Val |
| H5 | Gr-1 | 35 | 37.04 ± 4.2 | 871.83 ± 60.2 | Leu | Asn | Cys | Ser | Trp | Val |
| H6 | Bs-1 | 4 | 37.28 ± 9.5 | 753.95 ± 79.4 | Leu | Asn | Cys | Ser | Trp | Phe |
| H7 | Kondara | 7 | 39.26 ± 10.3 | 578.57 ± 58.4 | Leu | Asn | Trp | Ser | * | |
| H8 | CIBC-5 | 32 | 44.22 ± 4.2 | 752.87 ± 25.6 | Ser | Asn | Cys | Ser | Trp | Val |
| H9 | Col-0 | 31 | 54.51 ± 4.7 | 731.41 ± 37.8 | Leu | Asn | Trp | Ser | Trp | Phe |
| H10 | Ll-0 | 2 | 59 ± 19.6 | 747.15 ± 20.2 | Leu | Ile | Gly | Leu | | |
| H11 | Bor-4 | 10 | 72.72 ± 7.6 | 689.75 ± 27.9 | Leu | Asn | Trp | Phe | Trp | Phe |
| | | | | | | **frameshift** | **polar→ nonpolar** | | **stop gain** | **aromatic→aliophatic** |

| Haplotype | Representative ecotype | Number of ecotypes | RRL | Rainfall (mm/year) | Amino acid haplotypes | | | | | | |
|---|---|---|---|---|---|---|---|---|---|---|---|
| **AT1G14470.1** | | | | | **38** | **227** | **343** | **363** | **458** | **528** | **529** |
| H1 | Uk-1 | 1 | 17.4 | 745 | Gln | Lys | Arg | Val | Lys | Pro | Leu |
| H2 | Ak-1 | 2 | 25 ± 13 | 840.85 ± 95.8 | Gln | Lys | Arg | Ile | Lys | Pro | Leu |
| H3 | Pu2-23 | 2 | 29.1 ± 3.9 | 587 | Gln | Lys | | | | | |
| H4 | An-1 | 1 | 30.1 | 780.8 | Pro | Lys | Arg | Phe | Glu | Pro | Leu |
| H5 | Sp-0 | 2 | 38.35 ± 23.7 | 666.3 ± 87.3 | Gln | Lys | Agr | | | | |
| H6 | Ag-0 | 23 | 39.15 ± 4.2 | 940.72 ± 107.8 | Gln | Lys | Arg | Ile | Glu | Pro | Leu |
| H7 | Col-0 | 95 | 41.5 ± 2.5 | 749.73 ± 22.5 | Gln | Lys | Arg | Val | Glu | Pro | Leu |
| H8 | Bor-4 | 14 | 72.91 ± 6.3 | 754.9 ± 54.5 | Pro | Lys | Arg | Val | Glu | Pro | Leu |
| H9 | Na-1 | 1 | 88.9 | 622 | Pro | Lys | Arg | Ile | Lys | Pro | Leu |
| H10 | El-0 | 1 | 95.5 | 783 | Pro | Lys | Arg | Phe | Glu | Pro | Thr |
| | | | | | **polar→ nonpolar** | **frameshift** | **frameshift** | **aliphati c→aro** [illegible] | **negative →** [illegible] | **frameshift** | |

| Haplotype | Representative ecotype | Number of ecotypes | RRL | Rainfall (mm/year) | Amino acid haplotypes |
|---|---|---|---|---|---|
| **AT1G14480.1** | | | | | **367** |
| H1 | Pu2-23 | 2 | 29.1 ± 3.9 | 587 ± 0 | * |

| | | | | | |
|---|---|---|---|---|---|
| H2 | col-0 | 140 | 44.75 ± 2.2 | 781.2 ± 24.6 | Trp |

| **AT3G02510.1** | | | | | **341** | **351** | **389** |
|---|---|---|---|---|---|---|---|
| H1 | Aitba-2 | 1 | 31.3 | 423 | Ala | Met | Lys |
| H2 | Col-0 | 93 | 41.8 ± 2.6 | 830.7 ± 34.1 | Ala | Lys | Lys |
| H3 | Ts-1 | 37 | 44.72 ± 4.3 | 691.5 ± 25.9 | Ala | Lys | Arg |
| H4 | Ema-1 | 10 | 62.8 ± 10 | 678.2 ± 40.5 | Gly | Met | Lys |
| | Mer-6 | 1 | 81.6 | 503 | Gly | Lys | Lys |
| | | | | | | **non-polar→polar** | |

| **AT3G02520.1** | | | | | **238** |
|---|---|---|---|---|---|
| H1 | Col-0 | 133 | 43.0 ± 2.2 | 783.8 ± 25.8 | Ile |
| H2 | Ema-1 | 9 | 66.9 ± 10.2 | 699.8 ± 38.3 | Thr |
| | | | | | **non-polar→polar** |

| **AT4G26990.1** | | | | | **60** | **74** | **106** | **110** | **139** | **151** | **166** | **189** | **207** | **208** | **219** | **221** | **242** | **280** | **287** | **301** | **303** | **323** | **325** | **353** | **365** | **390** | **397** | **463** |
|---|---|---|---|---|---|---|---|---|---|---|---|---|---|---|---|---|---|---|---|---|---|---|---|---|---|---|---|---|
| H1 | Kz-9 | 1 | 24.3 | 352 | Ile | Ile | Lys | Gly | Glu | Pro | Glu | Glu | Ala | Lys | Ser | Pro | Pro | Gly | Ser | Glu | Val | Val | Ala | Ala | Val | Val | Ser | Val |
| H2 | Chat-1 | 1 | 29.4 | 626.6 | Ile | Ile | Lys | Gly | Glu | Pro | Glu | Asp | Glu | Lys | Asn | Thr | Ser | Glu | Asn | Gln | Gly | Thr | Thr | Ser | Ile | Ile | Pro | Phe |
| H3 | Ru3.1-31 | 2 | 34.8 ± 4.9 | 735.2 ± 43.8 | Ile | Lys | Asn | Ala | Gln | Ala | Gln | Glu | Ala | Lys | Ser | Pro | Pro | Gly | Ser | Glu | Val | Val | Ala | Ala | Val | Val | Ser | Val |
| H4 | Sei-0 | 1 | 35.1 | 974 | Ile | Ile | Lys | Ala | Gln | Ala | Glu | Asp | Glu | Ile | Asn | Thr | Ser | Glu | Asn | Gln | Gly | Thr | Thr | Ser | Ile | Ile | Pro | Phe |
| H5 | Baa-1 | 4 | 35.2 ± 15.3 | 708.5 ± 25.1 | Ile | Ile | Lys | Gly | Glu | Pro | Glu | Asp | Glu | Ile | Asn | Thr | Ser | Glu | Asn | Gln | Gly | Thr | Thr | Ser | Ile | Ile | Pro | Phe |
| H6 | Gel-1 | 1 | 37.4 | 734.9 | Ile | Ile | Lys | Gly | Glu | Pro | Glu | Asp | Glu | Ile | Ser | Thr | Ser | Glu | Asn | Gln | Gly | Thr | Thr | Ser | Ile | Ile | Pro | Phe |
| H7 | Col-0 | 90 | 41.3 ± 2.5 | 804.0 ± 35.8 | Ile | Ile | Asn | Ala | Gln | Ala | Gln | Glu | Ala | Lys | Ser | Pro | Pro | Gly | Ser | Glu | Val | Val | Ala | Ala | Val | Val | Ser | Val |
| H8 | Zdr-1 | 31 | 43.9 ± 5.3 | 776.6 ± 28.1 | Val | Lys | Asn | Ala | Gln | Ala | Gln | Glu | Ala | Lys | Ser | Pro | Pro | Gly | Ser | Glu | Val | Val | Ala | Ala | Val | Val | Ser | Val |
| H9 | Sha | 1 | 47.2 | 652 | Ile | Ile | Lys | Gly | Glu | Pro | Glu | Asp | Glu | Ile | Asn | Thr | Ser | Glu | Asn | Glu | Gly | Thr | Thr | Ser | Ile | Ile | Pro | Phe |
| H10 | Sorbo | 1 | 48.1 | 464 | Ile | Ile | Lys | Gly | Glu | Pro | Glu | Asp | Glu | Ile | Asn | Thr | Ser | Glu | Asn | Gln | Gly | Thr | Thr | Ser | Ile | Val | Ser | Phe |
| H11 | Stepn-1 | 1 | 48.2 | 469 | Ile | Ile | Asn | Gly | Glu | Pro | Glu | Glu | Ala | Lys | Ser | Pro | Pro | Gly | Ser | Glu | Val | Val | Ala | Ala | Val | Val | Ser | Val |
| H12 | Ei-2 | 1 | 56 | 1026 | Ile | Ile | Lys | Gly | Glu | Pro | Glu | Asp | Glu | Ile | Ser | Pro | Ser | Glu | Asn | Gln | Gly | Thr | Thr | Ser | Ile | Ile | Ser | Phe |
| H13 | Mammo-1 | 1 | 66.6 | 895.4 | Ile | Ile | Lys | Gly | Glu | Pro | Glu | Asp | Ala | Lys | Asn | Thr | Ser | Glu | Asn | Gln | Gly | Thr | Thr | Ser | Ile | Ile | Pro | Phe |
| H14 | Je-0 | 1 | 71.6 | 643.7 | Ile | Ile | Lys | Gly | Glu | Pro | Glu | Asp | Glu | Ile | Asn | Thr | Ser | Glu | Asn | Gln | Gly | Thr | Thr | Ala | Ile | Ile | Pro | Phe |
| H15 | Bor-4 | 1 | 74.6 | 587 | Ile | Ile | Lys | Gly | Glu | Pro | Glu | Asp | Glu | Ile | Ser | Pro | Ser | Glu | Asn | Gln | Gly | Thr | Thr | Ser | Ile | Ile | Pro | Phe |
| H16 | Condara | 1 | 92 | 464 | Ile | Ile | Lys | Gly | Glu | Ala | Glu | Asp | Glu | Ile | Asn | Thr | Ser | Glu | Asn | Gln | Gly | Thr | Thr | Ser | Ile | Val | Pro | Phe |
| H17 | Jl-3 | 1 | 94.2 | 586 | Ile | Ile | Lys | Gly | Glu | Ala | Glu | Asp | Glu | Ile | Asn | Thr | Ser | Glu | Asn | Gln | Gly | Thr | Thr | Ser | Ile | Ile | Pro | Phe |
| H18 | Da(1)-12 | 1 | 94.6 | 691 | Ile | Ile | Lys | Gly | Glu | Pro | Gln | Asp | Glu | Ile | Asn | Thr | Ser | Glu | Asn | Gln | Gly | Thr | Thr | Ser | Ile | Ile | Pro | Phe |
| H19 | Mnz-0 | 1 | 99.9 | 643 | Ile | Ile | Lys | Gly | Glu | Pro | Glu | Asp | Glu | Ile | Asn | Thr | Pro | Glu | Asn | Gln | Gly | Thr | Thr | Ser | Ile | Ile | Pro | Phe |
| | | | | | | **polar→ non-polar** | **uncharged →positive** | | **uncharged→negative** | **non-polar→ polar** | **uncharged→negative** | | **non-polar→ polar** | **non-polar→ polar** | | **ring→ open-chain** | **ring→ open-chain** | **non-polar→ polar** | | **negative→uncharged** | | **polar →non-polar** | **polar →non-polar** | | | | **ring→ open-chain** | **aliphatic→aromatic** |

| **AT4G27010.1** | | | | | **209** | **428** | **465** | **577** | **867** | **869** | **937** | **962** | **987** | **1011** | **1067** | **1082** | **1529** | **1613** | **1788** | **2011** |
|---|---|---|---|---|---|---|---|---|---|---|---|---|---|---|---|---|---|---|---|---|
| H1 | TueV-13 | 1 | 10.7 | 691 | Val | Lys | Val | Ser | Pro | Leu | Ile | Leu | Ala | Thr | Glu | Ala | Asn | Ser | Leu | His |
| H2 | Rennes-1 | 1 | 11 | 815 | Val | Lys | Val | Ser | Pro | Leu | Ile | Leu | Ala | Thr | Asp | Ala | Asn | Ser | Leu | His |
| H3 | Ciste-1 | 1 | 14.7 | 875 | Phe | Asn | Val | Ala | Pro | Phe | Leu | Pro | Glu | Thr | Glu | Val | Asp | Leu | Leu | Pro |
| H4 | Me-0 | 1 | 14.8 | 870 | Phe | Lys | Ile | Ala | Pro | Phe | Leu | Pro | Glu | Thr | Glu | Val | Asp | Leu | His | |
| H5 | Ag-0 | 1 | 17 | 979 | Val | Lys | Val | Ala | Pro | Leu | Ile | Leu | Glu | Met | Asp | Ala | Asn | Ser | His | |
| H6 | Wc-1 | 1 | 20.2 | 681.8 | Val | Lys | Val | Ser | Ser | Leu | Ile | Leu | Ala | Thr | Asp | Ala | Asn | Ser | Leu | His |
| H7 | Zdr-1 | 1 | 25.2 | 587 | Val | Asn | Val | Ala | Ser | Leu | Ile | Leu | Ala | Thr | Asp | Ala | Asn | Ser | Leu | His |
| H8 | Timpo-1 | 1 | 26.2 | 929 | Phe | Asn | Val | Ala | Pro | Phe | Leu | Pro | Glu | Met | Glu | Val | Asp | Leu | His | |
| H9 | Voeran-1 | 1 | 29.9 | 779 | Val | Lys | Val | Ser | Ser | Leu | Leu | Pro | Ala | Thr | Asp | Ala | Asn | Ser | His | |
| H10 | An-1 | 1 | 30.1 | 780.8 | Val | Asn | Val | Ser | Pro | Leu | Ile | Leu | Ala | Thr | Asp | Ala | Asn | Ser | His | |
| H11 | Rovero-1 | 1 | 33 | 778.6 | Phe | Asn | Val | Ala | Pro | Phe | Leu | Pro | Glu | Thr | Glu | Val | Asp | Leu | His | |
| H12 | Petro-1 | 1 | 33.3 | 721.6 | Val | Lys | Val | Ala | Ser | Leu | Ile | Leu | Ala | Thr | Asp | Ala | Asn | Ser | His | |
| H13 | Bolin-1 | 1 | 34.9 | 607.1 | Phe | Lys | Val | Ala | Pro | Phe | Leu | Pro | Glu | Met | Glu | Val | Asp | Leu | His | |
| H14 | Br-0 | 24 | 36.76 ± 3.5 | 875.37 ± 89.6 | Phe | Asn | Ile | Ala | Pro | Leu | Leu | Pro | Glu | Met | Glu | Val | Asp | Leu | His | |
| H15 | Slavi-1 | 1 | 38.4 | 627.7 | Val | Lys | Val | Ser | Pro | Leu | Leu | Leu | Glu | Thr | Asp | Ala | Asp | Ser | Leu | His |
| H16 | Ts-1 | 12 | 38.84 ± 7.5 | 759.86 ± 43.7 | Phe | Lys | Val | Ala | Pro | Phe | Leu | Pro | Glu | Thr | Glu | Val | Asp | Leu | Leu | Pro |
| H17 | Ru3.1-31 | 1 | 39.7 | 691.4 | Phe | Lys | Val | Ala | Pro | Leu | Leu | Pro | Glu | Thr | Glu | Val | Asp | Leu | Leu | His |

| | | | | | | | | | | | | | | | | | | | | |
|---|---|---|---|---|---|---|---|---|---|---|---|---|---|---|---|---|---|---|---|---|
| H18 | Ga-0 | 28 | 40.63 ± 4.7 | 792.11 ± 72.2 | Phe | Lys | Val | Ala | Pro | Phe | Leu | Pro | Glu | Thr | Glu | Val | Asp | Leu | His | |
| H19 | Gr-1 | 11 | 45.9 ± 9.6 | 814.7 ± 35.2 | Val | Lys | Val | Ser | Ser | Leu | Ile | Leu | Ala | Thr | Asp | Ala | Asn | Ser | His | |
| H20 | Np-0 | 1 | 48 | 639.5 | Phe | Lys | Val | Ala | Pro | Leu | Leu | Pro | Glu | Met | Glu | Val | Asp | Leu | His | |
| H21 | Sorbo | 1 | 48.1 | 464 | Phe | Asn | Val | Ala | Pro | Leu | Leu | Pro | Glu | Thr | Glu | Val | Asp | Leu | His | |
| H22 | HR-5 | 8 | 51.3 ± 11.3 | 735.6 ± 70.9 | Val | Lys | Val | Ser | Pro | Leu | Ile | Leu | Ala | Thr | Asp | Ala | Asn | Ser | His | |
| H23 | Bs-1 | 9 | [illegible] | 714.8 ± 53.3 | Phe | Lys | Val | Ala | Pro | Leu | Leu | Pro | Glu | Thr | Glu | Val | Asp | Leu | Leu | Pro |
| H24 | Col-0 | 8 | [illegible] | 972.5 ± 103.6 | Phe | Asn | Ile | Ala | Pro | Leu | Leu | Pro | Glu | Met | Glu | Val | Asp | Leu | Leu | His |
| H25 | Ciste-2 | 1 | 59.1 | 874.5 | Phe | Lys | Val | Ala | Pro | Phe | Leu | Pro | Glu | Met | Glu | Val | Asp | Leu | Leu | Pro |
| H26 | Bor-4 | 19 | 59.2 ± 6.1 | 628.7 ± 33.6 | Phe | Lys | Val | Ala | Pro | Leu | Leu | Pro | Glu | Thr | Glu | Val | Asp | Leu | His | |
| H27 | Bor-1 | 1 | 61.8 | 587 | Val | Lys | Val | Ser | Pro | Leu | Ile | Pro | Ala | Thr | Asp | Ala | Asn | Ser | His | |
| H28 | Uod-1 | 2 | 64.4 ± 36 | 833.7 ± 112.7 | Val | Lys | Val | Ser | Ser | Leu | Ile | Leu | Ala | Thr | Asp | Ala | Asn | Ser | Leu | His |
| H29 | Fei-0 | 1 | 69.6 | 1088 | Val | Lys | Val | Ser | Pro | Leu | Ile | Pro | Ala | Thr | Glu | Ala | Asn | Leu | Leu | His |
| H30 | Mer-6 | 1 | 81.6 | 503 | Phe | Lys | Ile | Ala | Pro | Leu | Leu | Pro | Glu | Thr | Glu | Val | Asp | Leu | His | |
| | | | | | **aromatic→ aliphatic** | **negative→p ositive** | | **non-polar→** | **ring→ open-** | **aliphati c→aro** | | **polar→ non-** | **non-polar→** | **non-polar→** | | | **negative →uncha** | **non-polar→** | **frameshift** | **positive→non-polar ring** |

**AT5G35695.1**

| | | | | | 43 | 47 | 51 | 70 | 80 | 100 | 118 | 125 | 161 | 177 | 181 | 184 | 193 | 209 | 212 |
|---|---|---|---|---|---|---|---|---|---|---|---|---|---|---|---|---|---|---|---|
| H1 | Yeg-1 | 1 | 9.7 | 439 | Ser | Arg | Leu | Arg | Gly | Val | Ala | Lys | Ala | Glu | Glu | Glu | Trp | Glu | * |
| H2 | Bsch-0 | 1 | 9.9 | 643 | Ser | Arg | Leu | Arg | Gly | Val | Ala | * | | | | | | | |
| H3 | Star-8 | 1 | 10.4 | 940 | Ser | Arg | Leu | Arg | Gly | Val | Val | Lys | Ala | Glu | Asp | Glu | Trp | Glu | * |
| H4 | Me-0 | 1 | 14.8 | 870 | Ser | Arg | Leu | Arg | Arg | Ala | Val | Lys | Ala | Glu | Asp | Glu | Trp | Glu | * |
| H5 | Aa-0 | 1 | 18.7 | 709 | Ser | Arg | Leu | Arg | Gly | Ala | Ala | Lys | Ala | Glu | Asp | Glu | Trp | Glu | * |
| H6 | Sij2 | 1 | 27.8 | 777.5 | Ser | Arg | Leu | Arg | Gly | Val | Ala | Lys | Val | Glu | Glu | Lys | Trp | Glu | * |
| H7 | An-1 | 3 | 32.53 ± 8.4 | 1132.73 ± 374.9 | Ser | Arg | Leu | Arg | Gly | Val | Ala | Lys | Val | Asp | Asp | Glu | * | | |
| H8 | Rovero-1 | 1 | 33 | 778.6 | Ser | Arg | Leu | * | | | | | | | | | | | |
| H9 | Bolin-1 | 1 | 34.9 | 607.1 | Ser | Arg | Leu | Arg | Gly | Val | Ala | Lys | Val | Glu | Asp | Lys | Trp | Glu | * |
| H10 | HKT2-4 | 9 | 35.9 ± 7.2 | 985.4 ± 187.4 | Ser | Arg (FS) | Trp | | | | | | | | | | | | |
| H11 | Gr-1 | 11 | 39 ± 4.96 | 806.94 ± 61.3 | Asn | Arg | Leu | Arg | Gly | Val | Ala | Lys | Ala | Glu | Asp | Glu | Trp | Glu | * |
| H12 | Gu-0 | 15 | 41.16 ± 6.1 | 833.49 ± 130.2 | Ser | Arg | Leu | Arg | Gly | Val | Ala | Lys | Ala | Glu | Asp | Lys | Trp | Glu | * |
| H13 | Br-0 | 18 | 41.23 ± 6.5 | 785.02 ± 57.9 | Ser | Arg | Leu | Arg | Gly | Ala | Val | Lys | Ala | Glu | Asp | Glu | Trp | Glu | * |
| H14 | CIBC-5 | 1 | 41.6 | 757 | Ser | Arg | Leu | Arg | Gly | Val | Ala | Lys | Val | Asp | Asp | Glu | Trp | Glu | * |
| H15 | Bor-4 | 20 | 42.26 ± 5.4 | 667.01 ± 38 | Ser | Arg | Leu | Arg | Gly | Val | Ala | Lys | Ala | Glu | Asp | Glu | Trp | Glu | * |
| H16 | Bor-1 | 12 | 44.23 ± 6.3 | 654.85 ± 25.3 | Ser | Arg | val | Arg | Gly | Val | Ala | Lys | Ala | Glu | Asp | Glu | Trp | Glu | * |
| H17 | Baa-1 | 4 | 47.15 ± 13.1 | 725.62 ± 35.2 | Ser | Arg | Leu | Arg | Gly | Val | Ala | Lys | Ala | Glu | Asp | Glu | Trp | Glu | Arg |
| H18 | Ak-1 | 3 | 48.07 ± 24.2 | 755.9 ± 101.4 | Ser | Arg | Val | Arg | Arg | Val | Ala | Lys | Ala | Glu | Asp | Glu | Trp | Glu | * |
| H19 | HR-5 | 2 | 51.9 ± 12.8 | 939 ± 182 | Ser | Arg | Leu | Arg | Gly | Val | Ala | Lys | Val | Glu | Asp | Glu | Trp | Glu | * |
| H20 | Tscha-1 | 3 | 52.8 ± 6.0 | 904.8 ± 75.5 | Ser | Arg | Leu | Arg | Arg | Val | Ala | Lys | Ala | Glu | Asp | Lys | Trp | Glu | * |
| H21 | Ll-0 | 18 | 56.5 ± 7.0 | 783.08 ± 33.6 | Ser | Arg | Leu | Arg | Arg | Val | Ala | Lys | Val | Glu | Asp | Lys | Trp | Glu | * |
| H22 | Kz-9 | 3 | 57.3 ± 22.5 | 575 ± 113.3 | Ser | Arg | Leu | Arg | Gly | Val | Ala | Lys | Ala | Glu | Asp | Lys | * | | |
| H23 | Borsk-2 | 3 | 58.17 ± 10.1 | 595.57 ± 32.3 | Ser | Arg | Leu | Arg | Gly | Val | Ala | Lys | Ala | Glu | Glu | Lys | Trp | Glu | * |
| H24 | Col-0 | 8 | 58.34 ± 14.0 | 824.44 ± 74.9 | Ser | Arg | Leu | Arg | Arg | Val | Ala | Lys | Val | Glu | Glu | Lys | Trp | Glu | * |
| H25 | Fei-0 | 1 | 69.6 | 1088 | Ser | Arg | Leu | Arg | Gly | Ala | Val | Lys | Ala | Glu | Glu | Lys | Trp | Glu | * |
| | | | | | | **frameshift** | | **stop gain** | **positive→non-polar** | | | **stop gain** | | | | **positive →negati ve** | **stop gain** | | **stop loss** |

**AT5G35698.1**

| | | | | | 44 | 62 | 63 | 77 |
|---|---|---|---|---|---|---|---|---|
| H1 | Star-8 | 1 | 10.4 | | * | | | |
| H2 | Col-0 | 118 | 43.84 ± 2.4 | 779.6 ± 27.8 | Cys | Glu | Lys | Lys |
| H3 | Ru3.1-31 | 22 | 46.16 ± 5.4 | 777.8 ± 47.9 | Cys | Glu | Lys | Glu |
| H4 | Mer-6 | 1 | 81.6 | | Cys | Glu | Asn | |

**frameshift** **positive→negative**

**Table S4.** Amino acid haplotypes in genes associated with the top 10 most-significant SNPs in the GWAS for drought tolerance in *Arabidopsis thaliana* . The variants were analyzed with the denser SNP information for 142 out of the 207 ecotypes available in the 1001 genomes database (See Table 3). The relative root length (RRL; %) for ecotypes containing a particular haplotype and the annual rainfall (mm) of the native geographical location of the ecotype are shown as averages with standard errors. The effects of the polymorphisms, viz., drastic changes in the protein (frameshift, stop gain and stop loss) and moderate effects like change in the chemical nature of amino acids, are indicated below. Asterisks indicate a stop codon. Grey cells indicate deviations from the reference haplotype (Col-0).
